# A unified evolutionary origin for the ubiquitous protein transporters SecY and YidC

**DOI:** 10.1101/2020.12.20.422553

**Authors:** Aaron J. O. Lewis, Ramanujan S. Hegde

## Abstract

Cells use transporters to move protein across membranes, but the origins of the most ancient transporters are unknown. Here, we analyse the ubiquitous protein-conducting channel SecY. Features conserved by its two duplicated halves suggest that their common ancestor was an antiparallel homodimeric channel. Structural searches with SecY’s halves detect exceptional similarity with the only other ubiquitous protein transporter, YidC. Their shared fold comprises a three-helix bundle interrupted by a helical hairpin. In YidC this hairpin is cytoplasmic and facilitates substrate delivery, whereas in SecY it is transmembrane and forms the substrate-binding lateral gate helices. In both, the three-helix bundle forms a protein-conducting hydrophilic groove, delimited by a conserved hydrophobic residue. We propose that SecY originated as a homodimeric YidC homolog. Many YidC homologs now use this interface to heterodimerise with a conserved partner. Unification of the two ubiquitous protein transporters would reconstruct a key step in the evolution of cells.

## Introduction

By the time of the last universal common ancestor (cenancestor), cells had already evolved a hydrophobic membrane and integral membrane proteins (IMPs) which carried out core metabolic functions (Coleman et al., 2021; Williams et al., 2017). Among those IMPs was SecY, a protein-conducting channel (Park and Rapoport, 2012). As is typical for channels, SecY (termed Sec61 in eukaryotes) catalyses the translocation of hydrophilic substrates across the hydrophobic membrane by creating a conducive hydrophilic environment inside the membrane. The substrates which it translocates are secretory proteins and the extracytoplasmic segments of IMPs.

SecY typically requires that its hydrophilic translocation substrates be connected to a hydrophobic α-helix (von Heijne, 1985; Krogh et al., 2001; Petersen et al., 2011). These hydrophobic helices serve as signals which open the SecY channel (Jungnickel and Rapoport, 1995; Li et al., 2016; Voorhees and Hegde, 2016). SecY is comprised of two separate halves (Van den Berg et al., 2004) which open like a clamshell when a helix binds to the lipid interface between them (Figure 1a).

**Figure 1.**
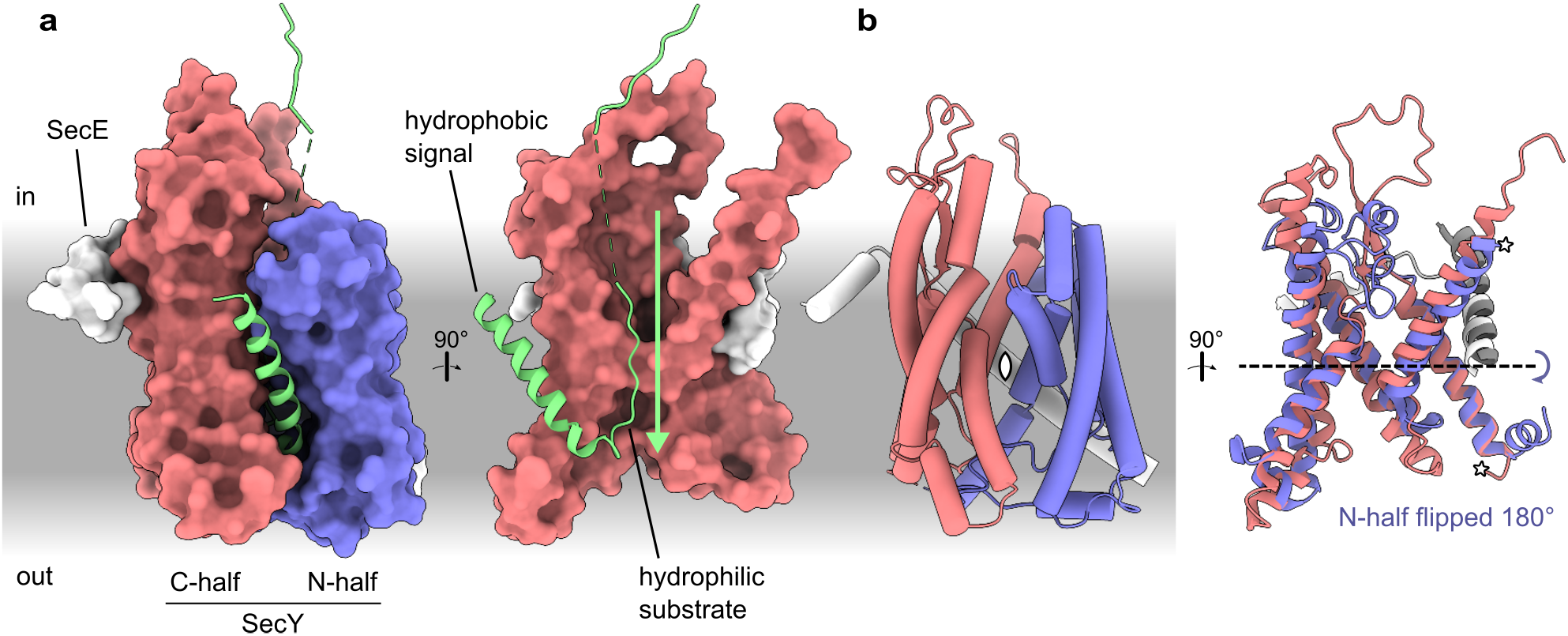
Structure and pseudosymmetry of the protein-conducting channel SecY. **a** Left: Structure of the channel engaged by a secretory substrate: *Geobacillus thermodenitrificans* SecYE engaged by proOmpA (Protein Data Bank ID [PDB] 6itc). The cytoplasmic ATPase SecA is present in the model but not shown. Right: Rotated view of only the SecY N-half and substrate. **b** Pseudosymmetry of the N- and C-halves. Left: SecYE shown as tubes with the pseudo-*C*_*2*_ symmetry axis denoted by a pointed oval. Right: Rotated view in ribbon representation. The N-half has been rotated 180° around the pseudo-*C*_*2*_ symmetry axis and aligned to the C-half. SecE is divided where it intersects the symmetry axis into N-terminal (white) and C-terminal (grey) segments. A dashed black line indicates the same pseudo-*C*_*2*_ symmetry axis shown at left after a 90° rotation. Stars indicate where the halves were split.

Spreading the halves apart destabilises a plug which sits between them, opening a hydrophilic pore that spans the width of the membrane. By binding at this site, the signal also threads one of its hydrophilic flanking regions through the hydrophilic pore, thereby initiating its translocation.

A set of conserved hydrophobic residues that line the narrowest part of the translocation pore form a gasket-like seal around the translocating chain (Ma et al., 2019). These residues, which are contributed by both halves of SecY, are known collectively as the pore ring. They not only maintain the ion permeability barrier across the membrane (Dalal and Duong, 2009; Park and Rapoport, 2011), but also bind the plug when the channel is closed (Van den Berg et al., 2004).

The site between SecY’s halves where signals bind is called the lateral gate. After binding and initiating translocation, sufficiently hydrophobic signals can diffuse away from the lateral gate into the surrounding hydrophobic membrane (Hessa et al., 2005). Many signals, particularly those at the N-terminus of secretory proteins, are ultimately cleaved off by signal peptidase, a membrane-anchored protease whose active site resides on the extracytoplasmic side of the membrane (Paetzel et al., 2002). Longer and more hydrophobic signals that are not cleaved serve as the transmembrane helices (TMHs) of IMPs (White and von Heijne, 2005).

SecY is the only ubiquitous transporter for protein secretion. There is however a second ubiquitous superfamily of protein transporters which is specialised for IMP integration, Oxa1 (Anghel et al., 2017). The Oxa1 superfamily consists of four member families, each of which now has a known atomic structure. One, YidC, is found in the prokaryotic plasma membrane (Borowska et al., 2015; Kumazaki et al., 2014), whereas the other three are paralogs located in the eukaryotic endoplasmic reticulum (ER): TMCO1 (McGilvray et al., 2020), EMC3 (Bai et al., 2020; Miller-Vedam et al., 2020; O’Donnell et al., 2020; Pleiner et al., 2020), and GET1 (McDowell et al., 2020). All share a conserved core of three TMHs and a cytoplasmic helical hairpin. With YidC also present in the plastid (Alb3; Sundberg et al., 1997) and mitochondrial inner membranes (Oxa1; Bauer et al., 1994; Bonnefoy et al., 1994), it appears that every membrane equivalent to the plasma membrane of the cenancestor contains Oxa1 superfamily proteins. As with SecY, archaeal and bacterial YidC are monophyletic and highly divergent (Anghel et al., 2017; Yen et al., 2001), suggesting that YidC was present alongside SecY in the cenancestor.

Like SecY, YidC facilitates IMP integration by translocating extracytoplasmic segments across the membrane (Borowska et al., 2015; Cymer et al., 2015; Hell et al., 2001; Samuelson et al., 2000). Unlike SecY substrates however, YidC substrates are limited in the length of polypeptide translocated, typically to less than 30 amino acids (Shanmugam et al., 2019). This limitation may be due to YidC’s lack of a membrane-spanning hydrophilic pore; instead, YidC structures show a membrane-exposed hydrophilic groove that only penetrates partway into the membrane (Kumazaki et al., 2014). YidC thus forms a partial channel, and may also thin and distort the adjacent membrane (Chen et al., 2017).

The two halves of SecY are structurally similar and related by a two-fold rotational (*C*_*2*_) pseudosymmetry axis parallel to the membrane plane (Van den Berg et al., 2004; Figure 1b). Such pseudosymmetry is common among membrane proteins and arises when the gene encoding an asymmetric progenitor undergoes duplication and fusion (Forrest, 2015). Channels are particularly likely to have a membrane-parallel *C*_*2*_ axis of structural symmetry because they have the same axis of functional symmetry: they facilitate substrates’ bidirectional diffusion across the membrane.

Indeed, polypeptides can slide through SecY bidirectionally (Ooi and Weiss, 1992), with unidirectionality arising from other factors (Erlandson et al., 2008; Matlack et al., 1998). Membrane-parallel *C*_*2*_ pseudosymmetry requires that the two fused domains be antiparallel, and thus those domains typically derive from progenitors that existed as antiparallel homodimers (Lolkema et al., 2008; Rapp et al., 2006).

The ubiquity and essentiality of the SecY channel motivated us to investigate how it might have evolved. We identify several structural elements that are conserved between its two halves, which suggest that the SecY progenitor was an antiparallel homodimer featuring a symmetric pore ring at its dimerisation interface. Automated database searches for structures similar to the SecY halves show that they are uniquely similar to the Oxa1 superfamily, of which YidC is the prokaryotic member. Structural alignments indicate that key residues of YidC’s hydrophilic groove and its capping hydrophobic residue are homologous to key residues in SecY’s hydrophilic funnels and its pore ring, respectively.

In light of this new correspondence, we re-evaluate the extensive mechanistic literature on SecY and the Oxa1 superfamily, identifying surprising similarities and specific structural bases for their differences. Based on this analysis, we propose that SecY evolved from a dimeric Oxa1 superfamily member by gene duplication and fusion. We compare the range of substrates that can be translocated by YidC to the prokaryotic membrane proteome, and find that a YidC-dependent,

SecY-independent cell is plausible. We discuss the implications of this model for the evolution of YidC itself and other components of the general secretory pathway.

## Results

### Conserved pre-duplication features in SecY

Features shared by both of SecY’s halves are likely to have been present in their last common ancestor, which we term proto-SecY. In an attempt to identify conserved sequence features of proto-SecY, we aligned the amino acid (a.a.) sequences of a set of N- and C-halves. However, their pairwise identities are just 12.5 ± 2.2% (s.d.), compared to 9.3 ± 4.3% between randomly shuffled sequences, an excess identity of only 6 a.a. per 200 a.a. half. By pairwise HHpred (Zimmermann et al., 2018), the halves have similarity *p* = 0.02, where *p* estimates the likelihood of observing as much similarity between a random pair of unrelated proteins (Remmert et al., 2012; Söding, 2005). For context, this means that in searching a typical whole-proteome database of ∼10^4^ entries with one half of SecY, one would expect to find ∼200 unrelated proteins just as similar as the other half of SecY. Reconstructing a cenancestral SecY sequence using methods previously successful for a different internally duplicated protein (Longo et al., 2020) yielded no increase in similarity between the SecY halves (see Supplementary Note). Thus, the two halves of SecY have diverged too far from one another to reliably reconstruct proto-SecY’s primary sequence.

Unlike primary sequence, a five-TMH tertiary structure is conserved by both halves of SecY (Figure 1b; Van den Berg et al., 2004). To facilitate comparisons, we label these five consensus helices H1-H5 (Figure 2a). A prefix of N or C is used when referring to a specific instance of a consensus element in the N- or C-half of SecY. For example, TM6 of SecY is labelled C.H1 because it is located in the C-half and corresponds to H1 of proto-SecY, as does TM1 (N.H1) in the N-half. Flanking and intervening segments are labelled using lower-case references to the nearest consensus elements. For example, the ribosome-binding loop between C.H1 and C.H2 is C.h1h2. The N-terminal peripheral helix of each half, which we argue later was probably also present in proto-SecY, is named H0.

**Figure 2.**
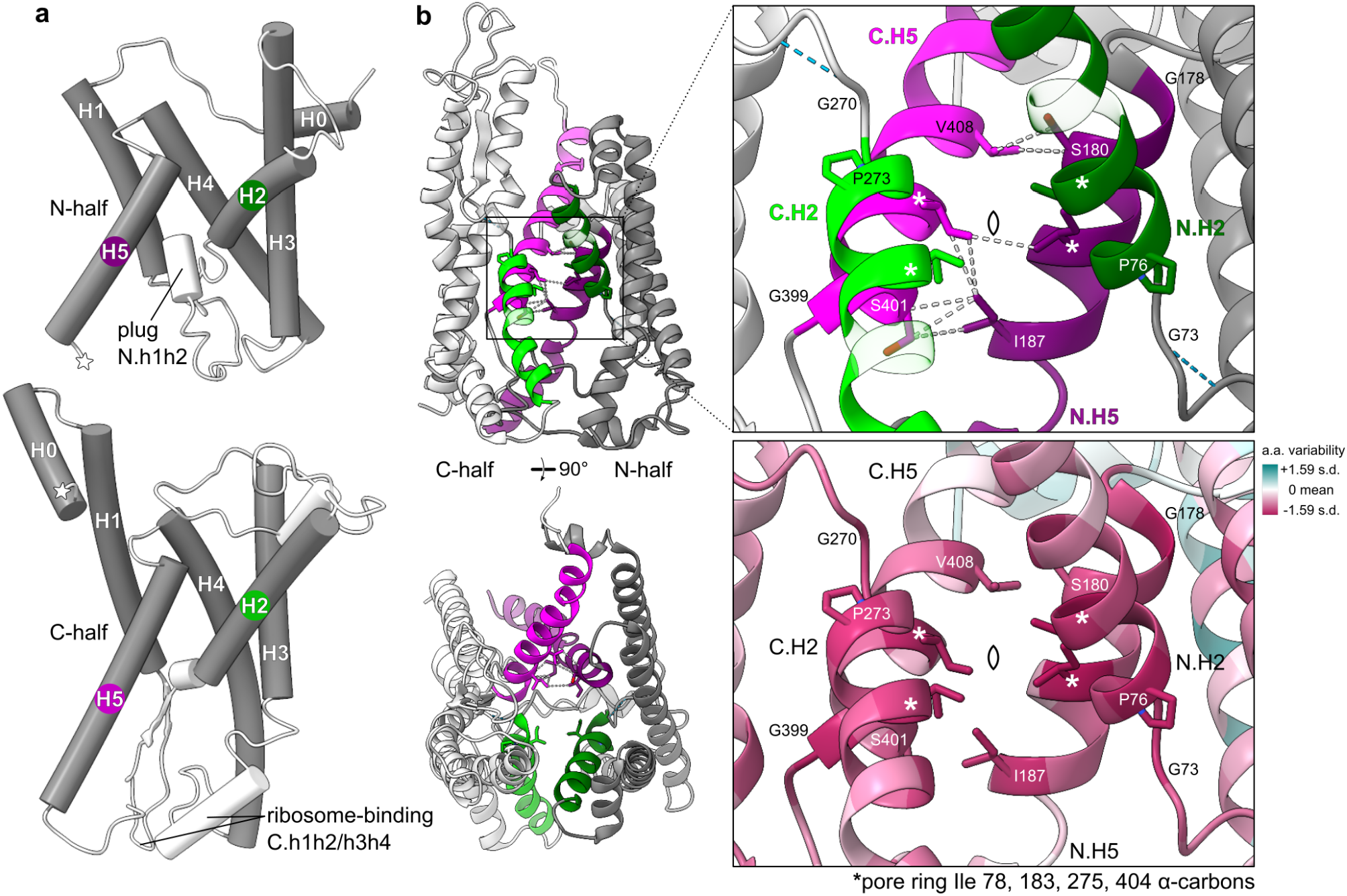
Features conserved between SecY’s halves. **a** Consensus secondary structure elements (grey) in *Methanocaldococcus jannaschii* SecY (1rhz). Stars indicate where the halves were split. **b** Symmetric features in the most self-similar SecY structure (6itc). A pointed oval indicates the pseudo-*C*_*2*_ symmetry axis. Dashed lines indicate hydrogen bonds (blue) and Van der Waals contacts (white). ConSurf variability scores for an alignment of the N- and C-half sequences are shown mapped onto each half’s model. The colour scale encompasses the minimum but not maximum score. The most conserved residues are shown as sticks and labelled. **Figure 2-Figure supplement 1**. Structural similarity and symmetry breaking between SecY halves. **Figure 2-Source data 1**. Structure-guided sequence alignment of the SecY halves.

To identify more detailed conserved features, we pursued a precise structural alignment of the SecY halves. We collected a representative set of seven SecY structures: closed (Itskanov et al., 2021; Tanaka et al., 2015), primed (Braunger et al., 2018; Zimmer et al., 2008), and open (Ma et al., 2019; Voorhees and Hegde, 2016) models for eukaryotic and bacterial SecY, and closed archaeal SecY (Van den Berg et al., 2004). Alignments between these halves generated by mTM-align (Dong et al., 2018a) vary widely in accuracy and extent (Figure 2-Figure supplement 1a), but display one clear trend: the C-halves in the closed and primed structures are least like the N-halves of any structure. This is because closure induces symmetry-breaking tilts in C.H2 and C.H5 (Figure 2-Figure supplement 1b) whereas the N-halves remain relatively unchanged.

Intriguingly, the stability of this asymmetrical closed conformation depends on another asymmetrical feature, the plug (Figure 2a; Li et al., 2007). This suggests that neither the plug nor the closed conformation may have been present in proto-SecY. A plugless proto-SecY is plausible, given that plug-deletion mutants of SecY are tolerated (Junne et al., 2006; Li et al., 2007; Park and Rapoport, 2011). If proto-SecY did lack SecY-like gating or a plug, it would then more closely resemble other protein transporters like YidC or TatC, which are not gated (Kumazaki et al., 2014; Rollauer et al., 2012).

The most similar halves, 6fti N (Braunger et al., 2018) and 6itc C (Ma et al., 2019), share a common core (< 4 Å deviation) of 121 a.a. across all 5 helices with 1.9 Å RMSD (Figure 2-Figure supplement 1c). This is precise enough that all 5 helices can be registered confidently. Their alignment shows that the four functionally important pore ring residues (Dalal and Duong, 2009) are located at the same two homologous sites in each half. To identify other conserved sites, this structural alignment was then used to align a diverse set of N- and C-half sequences. Their sequence conservation at each site was then scored and mapped to the most self-similar SecY structure (6itc).

This scored structure shows that the interface between halves is symmetrical and conserved (Figure 2b). N.H5 and C.H5 contact each other via the H5 pore ring residue, which coincides with the symmetry axis, and also via residues −3 and +4 a.a. from the pore ring. Although SecY today is a pseudodimer, split mutants show that it remains able to form true dimers via this interface (Wilkinson et al., 1997). Less conserved than the pore ring but still notable are two helix-breaking residues which N-terminate H5 (glycine) and H2 (proline), and a glycine near H2 which bonds its α-hydrogen with the -3 backbone oxygen, thereby stabilising a small bulge. These conserved residues are all within 5 a.a. of the pore ring, underscoring the structural conservation of this central region. Altogether, these features suggest that while proto-SecY may not have had SecY-like gating or a plug, it did form antiparallel homodimers centred on a pore very similar to SecY’s. Thus proto-SecY likely functioned as a protein-conducting channel.

### SecY is uniquely similar in structure to the Oxa1 superfamily

With this information about proto-SecY, we sought to identify distant homologs from before its duplication. For this we used Dali, which measures structural similarity between protein backbones. Dali is competitive with other top methods for accurate homolog detection, and outperforms them when the relationships in question are particularly distant (Holm, 2020). Other methods construct 3-D superpositions with better geometric properties like RMSD, but Dali nonetheless outperforms them in detection accuracy (Kolodny et al., 2005). Thus we use Dali here, whereas a method optimised for 3-D superposition, mTM-align (Dong et al., 2018a), was used above to align the SecY halves.

Queries of the PDB with the N- or C-half of SecY yielded a match correlation matrix (Tai et al., 2011) that indicates the possible presence of two separate subdomains (Figure 3a). The three-helix bundle of H1/4/5 showed positive self-correlation, but anti-correlation with the H2/3 two-helix hairpin. Because Dali measures global similarity, including both subdomains in our searches would tend to obscure distant homologs which share only one subdomain. We therefore performed searches with not only the whole N- and C-halves, but also their H1/4/5 subdomains (Figure 3b). We queried a non-redundant subset of the PDB filtered at 25% pairwise identity (PDB25).

**Figure 3.**
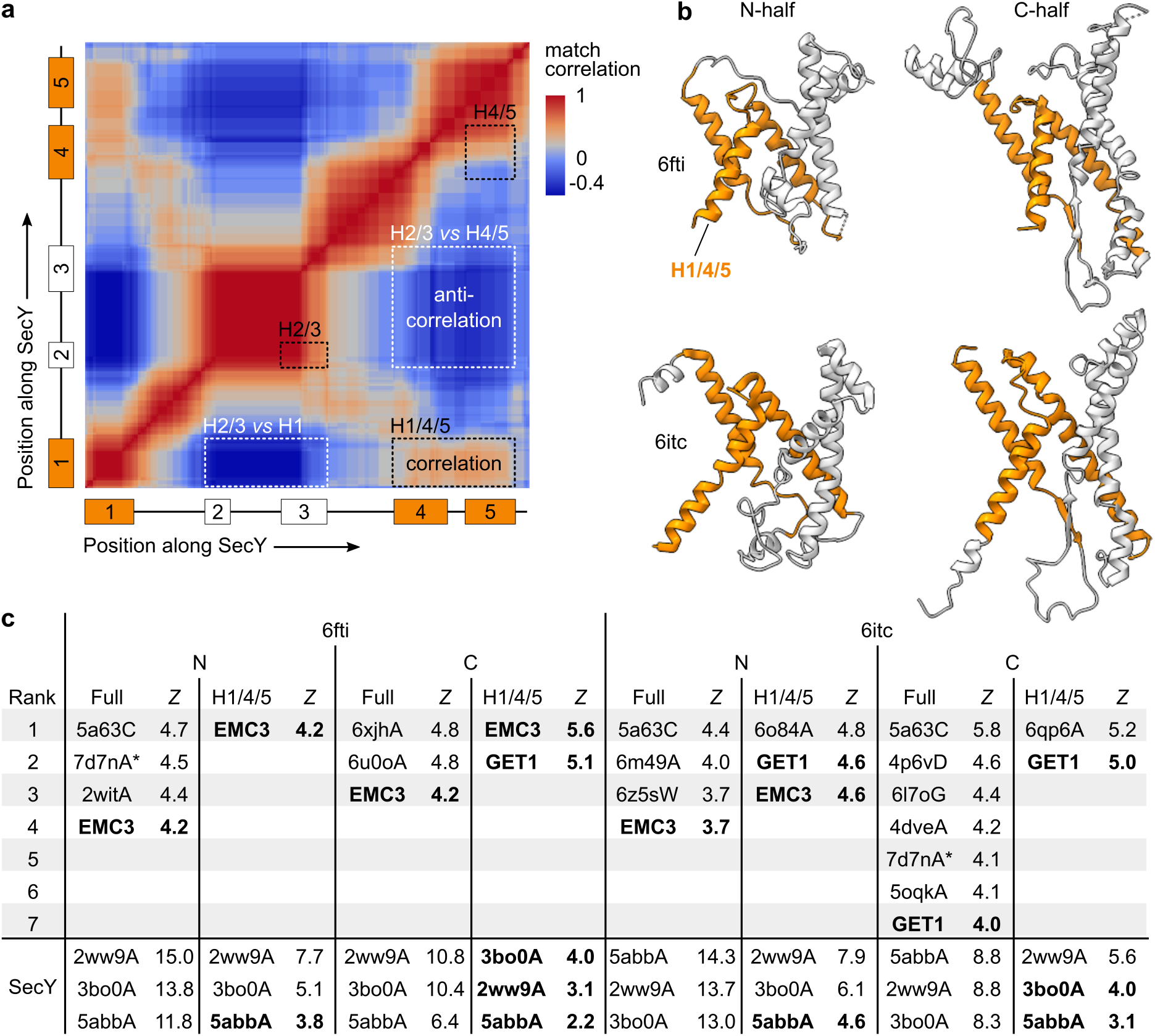
SecY’s halves are uniquely similar in structure to the Oxa1 superfamily. **a** Match correlation matrix returned by Dali for a half-SecY query (6itc C). The axes are labeled by a diagram of the SecY transmembrane helices. **b** The structural models used as Dali queries. The full models and the H1/4/5 subdomains (orange) were used. **c** Results from querying the PDB25. The top-ranking hits for each query are shown, and any lower-ranking hits that rank higher than the first Oxa1 superfamily hit. Asterisks mark 7d7nA because although it appears twice, those hits are with two non-overlapping parts of the model. Oxa1 superfamily hits are shown by name (EMC3, GET1) instead of PDB code (6ww7C, 6so5C). At bottom are the scores for the SecY hits, which were excluded from the ranking. SecY hits in boldface scored lower than an Oxa1 superfamily hit for that query. **Figure 3-Figure supplement 1**. Structures of non-Oxa1 superfamily top Dali hits. **Figure 3-Source data 1**. Results from Dali queries of the PDB25.

After excluding SecY and soluble hits, the most consistently high-ranking hits were members of the Oxa1 superfamily (Figure 3c). Moreover these hits link multiple Oxa1 families (GET1 and EMC3) to both SecY halves, suggesting that their similarity is due to conserved characteristics of the Oxa1 superfamily and proto-SecY rather than idiosyncrasies of any one structure. By contrast, almost all other hits were as highly ranked in only a single query. Manual review of these isolated hits shows them to be obviously dissimilar (Figure 3-Figure supplement 1a) due to features ignored by Dali’s distance matrix metric, such as gaps, context, and handedness. There is one non-Oxa1 hit that tops multiple queries, APH-1 (5a63C; Figure 3-Figure supplement 1b), but only two of the four aligned TMHs are conserved by the prokaryotic proteases from which APH-1 descends (Pei et al., 2011; Schaeffer et al., 2019), so the relevant part of this alignment is negligible. To test the sensitivity of these results to our choice of queries (6fti N and 6itc C), selected above for maximum symmetry, we repeated them with the opposite half of each structure (6fti C and 6itc N), with similar results.

In the H1/4/5 queries, the Oxa1 superfamily hits rank even higher than some SecY hits, and have Dali *Z-*scores 4.2 to 5.6 standard deviations above the mean, *i*.*e. p* = 0.0081 to 0.0014. This means that Dali predicts one would find an unrelated cenancestral protein this similar if the cenancestor contained 0.0014^-1^ ≈ 700 or more homology candidates (multi-pass helical IMPs non-redundant at 25% identity). For scale, *E. coli* contains ∼550 such proteins. Fewer such proteins can be confidently assigned to the cenancestor (Coleman et al., 2021; Williams et al., 2017), but the uncertainties involved are large. Thus in absolute terms, these *p*-values do not provide strong evidence for homology. But they do show that in relative terms, the SecY halves are more similar to the Oxa1 superfamily than any other.

Moreover, each consensus helix from proto-SecY can be matched to a consensus helix from the Oxa1 superfamily and linked with the same connectivity (Figure 4, Table 1). The SecY/Oxa1 fold comprises a right-handed three-helix bundle (H1/4/5) interrupted after the first helix by a helical hairpin (H2/3) and prefixed by an N-terminal peripheral helix (H0) which abuts H4 (Figure 4-Figure supplement 1). Thus in addition to sharing a universally conserved core three-TMH bundle, proto-SecY and Oxa1 proteins share a similar composition and connectivity across their full ∼200 a.a. lengths.

**Figure 4.**
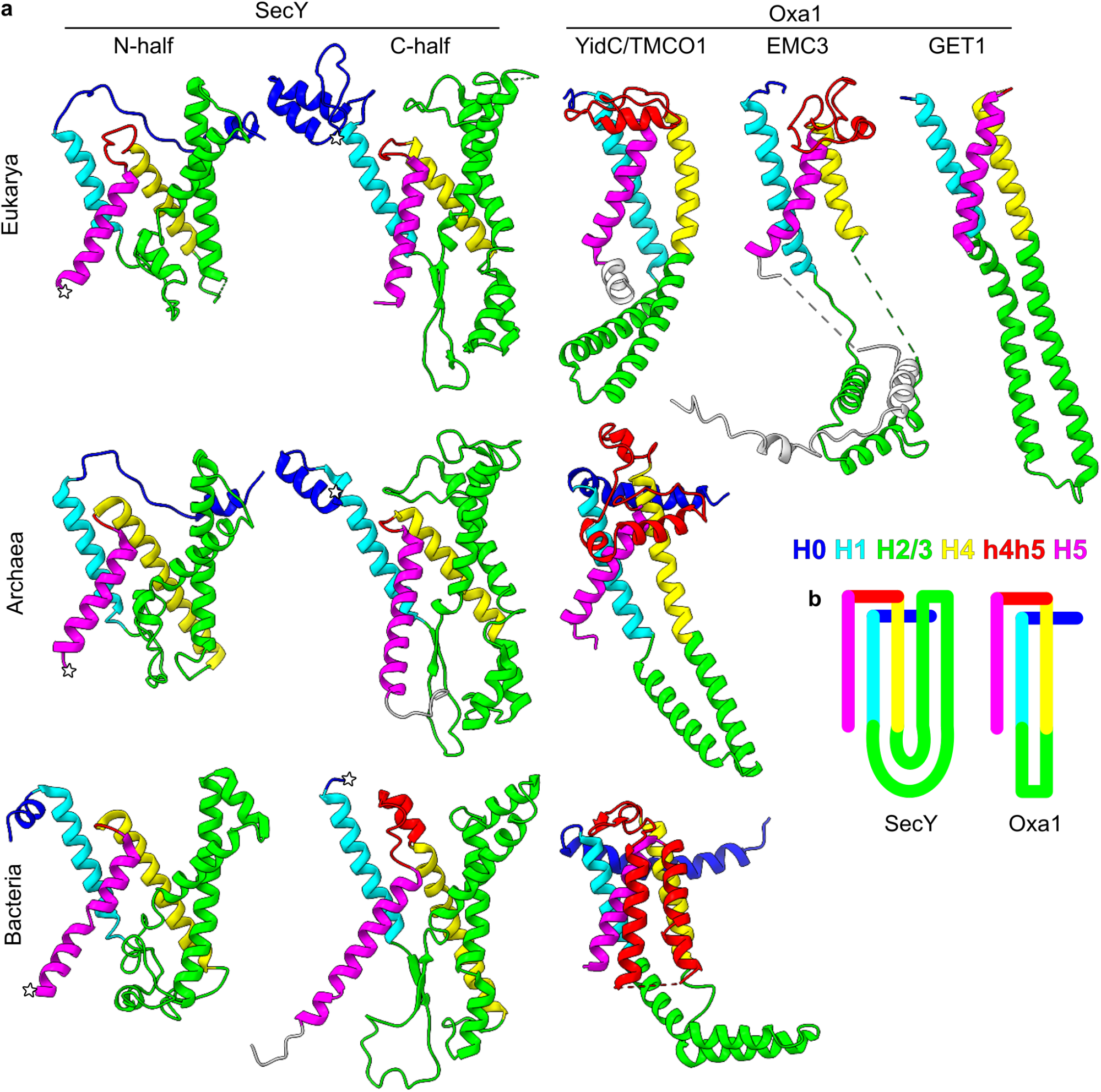
Correspondence between structural elements of SecY and the Oxa1 superfamily. Consensus elements and the intervening element h4h5 are coloured according to the key shown. Other intervening elements are coloured to match a neighbouring consensus element, and flanking elements are coloured white. The models are, from left to right and then top to bottom, *Canis lupus familiaris* SEC61A1 (6fti), *Homo sapiens* TMCO1 (6w6l), *H. sapiens* EMC3 (6ww7), *H. sapiens* GET1 (6so5), *M. jannaschii* SecY (1rhz), *M. jannaschii* MJ0480 (5c8j, extended by structure prediction to include the originally unmodelled H2/3; see Figure 4-Figure supplement 4), *G. thermodenitrificans* SecY (6itc), *Bacillus halodurans* YidC2 (3wo6). **Figure 4-Figure supplement 1**. Structural comparison of H0 in SecY and YidC. **Figure 4-Figure supplement 2**. Structure of the acquired transmembrane hairpin in SecE. **Figure 4-Figure supplement 3**. Amino acid conservation in bacterial YidC. **Figure 4-Figure supplement 4**. Crystallised and predicted structures of archaeal YidC. **Figure 4-Source data 1**. Predicted structures of archaeal YidC.

**Table 1.**
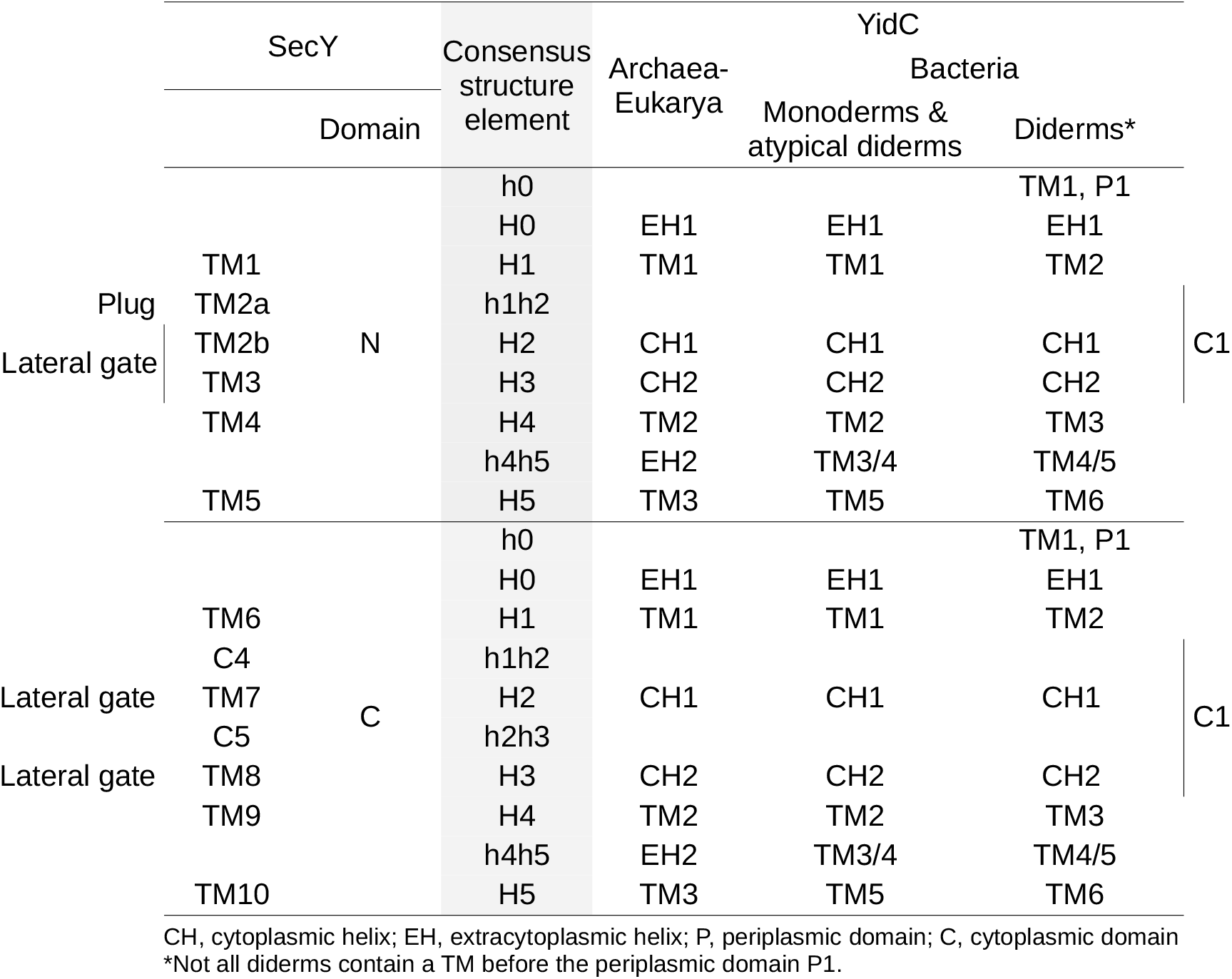
Consensus nomenclature for SecY and YidC.

There is one conspicuous difference between these groups’ structures: in SecY, the helical hairpin H2/3 is transmembrane, but in the Oxa1 superfamily it is cytoplasmic (Figure 4b). If SecY derived from an Oxa1 superfamily ancestor, this would suggest that an initially cytoplasmic H2/3 evolved to be transmembrane in the proto-SecY stem lineage. Transmembrane hairpins are indeed known to be acquired during membrane protein evolution; convenient examples are provided by the transmembrane hairpins in bacterial YidC h4h5 (Figure 4) and in some SecE (Figure 4-Figure supplement 2).

Starting from a YidC-like H2/3, more membrane-penetrating conformations could have been induced by hydrophobic substitution mutations around the hairpin tip, which lacks conserved hydrophilics (Figure 4-Figure supplement 3). SecY H2/3 could also derive to some degree from indel mutations, particularly since the segment between H1 and H4 is 10 a.a. longer in SecY N than in YidC, and 60 a.a. longer in SecY C. Mutant H2/3 would readily sample membrane-penetrating conformations because it rests at the lipid-water interface (Chen et al., 2017) and is flexibly connected to H1/4/5, as evident in simulations (Chen et al., 2017; Kumazaki et al., 2014) and in the archaeal and bacterial crystal structures where H2/3 was too mobile to be modelled (Borowska et al., 2015; Xin et al., 2018). Because SecY H2/3 is stabilised in the membrane by H1/4/5, it need not have become particularly hydrophobic; for example, most of the H2/3 helices in *G. thermodenitrificans* SecY are predicted to prefer the aqueous phase (N to C: Δ*G*_*app*_ = 1.7, 0.7, 1.0, −1.4 kcal/mol; Hessa et al., 2007).

Late acquisition of the transmembrane H2/3 would explain a curious feature of SecY’s structure. H2/3 does not pack against H1 (Figure 4-Figure supplement 1), despite the fact that during co-translational membrane insertion H1 would be exposed to H2/3 without competition. It is reasonable to expect that these elements would interact if their folding pathway had juxtaposed them throughout evolution. This is thought to be why most transmembrane helices pack sequentially against one another (Bowie, 1997; Gimpelev et al., 2004). In SecY however, H1 and H2/3 are separated by H4/5. This suggests that H1/4/5 was the original, sequentially packed transmembrane bundle, and H2/3 only later became transmembrane and packed against its surface. The transmembrane hairpins in bacterial YidC h4h5 and SecE likewise break sequential packing, suggesting that a similar process of transmembrane hairpin acquisition may explain non-sequential TMH packing in other proteins. This is analogous to how RNA branch acquisition left structural fingerprints in the ribosome (Petrov et al., 2014).

Thus the SecY halves and the Oxa1 superfamily have backbone structures that not only are uniquely similar by standard measures, but also could plausibly descend from a common ancestor. This identifies the Oxa1 superfamily as the best candidate for the origin of SecY. The following sections analyse their similarities and differences in mechanistic and functional terms. We focus on archaeal and bacterial YidC and not their eukaryotic homologs, since eukaryotes derive from archaea (Williams et al., 2020).

### Corresponding elements of SecY and YidC are mechanistically similar

YidC’s hydrophilic groove is functionally similar to those recently observed in components of the retrotranslocation machinery for ER-associated degradation (ERAD; Wu et al., 2020). There, the membrane proteins Hrd1 and Der1 each display hydrophilic grooves, which are open to the cytosol and ER lumen, respectively. The juxtaposition of these two partial channels forms a nearly continuous hydrophilic pore, interrupted by only a thin membrane through which polypeptide translocation is thought to occur. A YidC homolog that formed antiparallel homodimers could similarly create a near-complete channel by juxtaposing two grooves, one on each side of the membrane. Adaptation could then yield the complete channel of proto-SecY.

In both YidC and SecY, the hydrophilic translocation interface is lined by H1/4/5 (Figure 5a). Both three-TMH bundles have a right-handed twist, with H1 and H4 near parallel and H5 packing crossways against them. Of the three helices, it is this crossways H5 that makes the closest contacts with the translocating hydrophilic substrate in both SecY (Figure 5b) and YidC, as determined by chemical crosslinking experiments in *E. coli* (He et al., 2020). These crosslinking data also indicate that YidC’s substrates initiate translocation as a hairpin with both termini in the cytoplasm (Figure 5a), just as SecY’s substrates do (Mothes et al., 1994; Shaw et al., 1988). From this intermediate state, TMHs can integrate into the membrane, and their propensity to integrate is a very similar function of the TMH’s sequence regardless of whether YidC or SecY is used (Xie et al., 2007).

**Figure 5.**
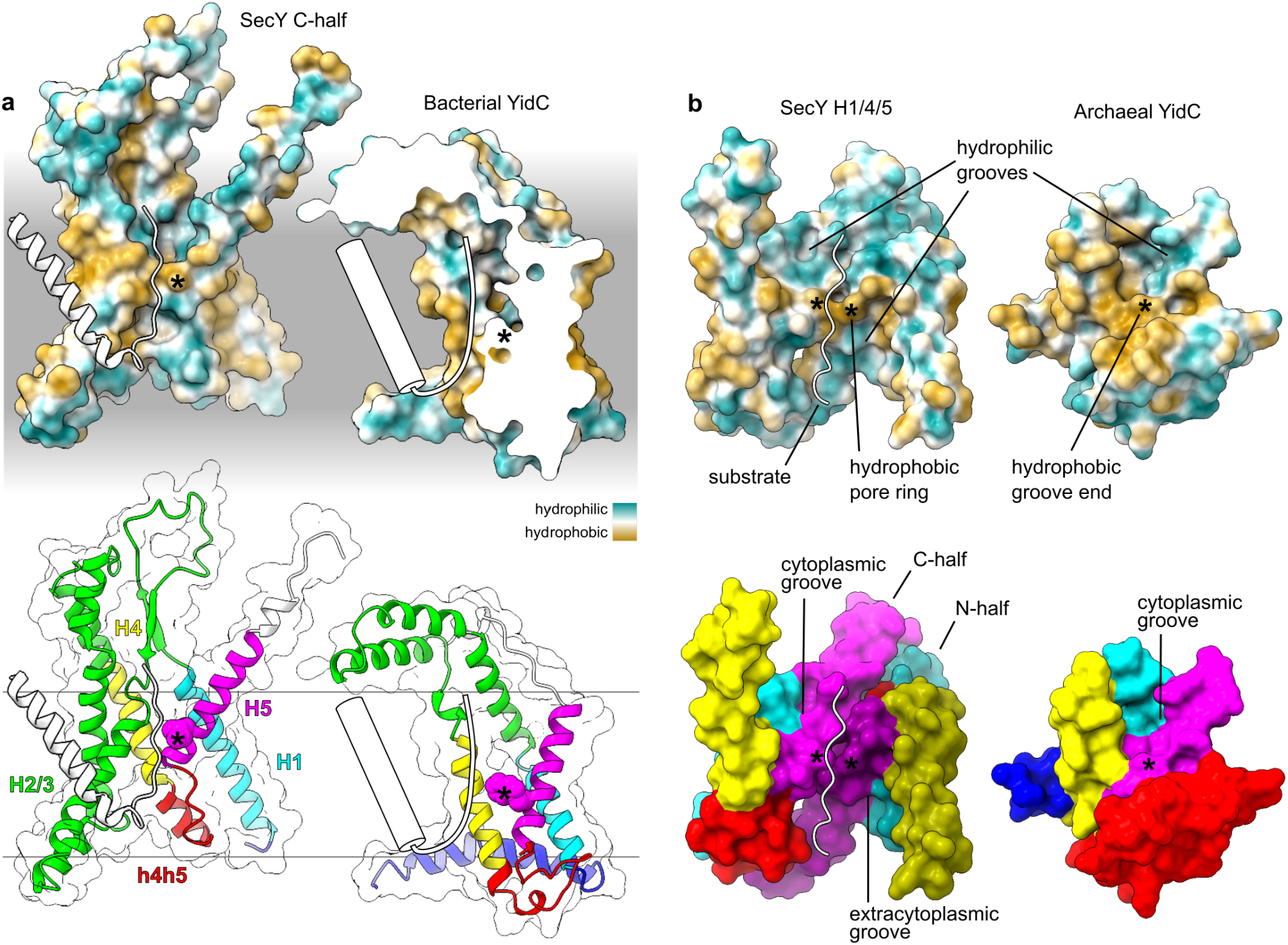
Structural and mechanistic comparison of SecY and YidC. The SecY/substrate model is *G. thermodenitrificans* SecY/proOmpA (6itc). **a** Comparison between the SecY C-half (left) and *B. halodurans* YidC2 (right; 3wo7A). The H5 pore ring residue and the groove end residue are both indicated by asterisks. The hydropathy of the aqueous and lipidic phases ranges from hydrophilic (white) to hydrophobic (grey). Models are oriented and positioned relative to the membrane as described in Methods. A cartoon substrate signal and translocating polypeptide is superimposed on YidC2 to indicate the experimentally determined interface across which substrates translocate and hairpin conformation in which they do so (He et al., 2020). The YidC2 surface and model are clipped to allow a lateral view of the hydrophilic groove which would otherwise be occluded by the bacteria-specific h4h5 transmembrane hairpin (*B. halodurans* YidC2 TM3/4). **b** Left: Lateral view of the SecY/substrate complex showing only the H1/4/5 core of each half. Right: archaeal YidC (*M. jannaschii* MJ0480, 5c8j) in the same relative orientation as SecY.

The YidC and SecY H1/4/5 bundles are structurally similar enough that they can be aligned confidently (Figure 6a). This alignment superimposes the pore ring residue in SecY H5 onto a conserved hydrophobic residue in YidC H5 that marks the end of the hydrophilic groove. In YidC and the SecY N-half this residue is positioned at a similar depth in the membrane, after accounting for the N-half’s inversion (Figure 6b), whereas the C-half is shifted toward the cytoplasm. In bacterial YidC the groove end residue is aromatic and intimately contacts the bacteria-specific h4h5 hairpin, but in archaeal YidC this residue is aliphatic and most often an isoleucine, just as it is in SecY (Figure 6c). Moreover, the same surrounding positions on H5 are polar (−3, +3, +7, +11) or polarisable aromatics (−1) in both YidC and SecY. Together with a conserved polar residue in H1, these comprise the entire hydrophilic groove of archaeal YidC, and thus that same groove is also hydrophilic in SecY. Finally, a conserved tryptophan is positioned at the lipid-water interface, tryptophan’s preferred environment (Yau et al., 1998), where it is thought to stabilise YidC’s particular transmembrane position (Chen et al., 2017).

**Figure 6.**
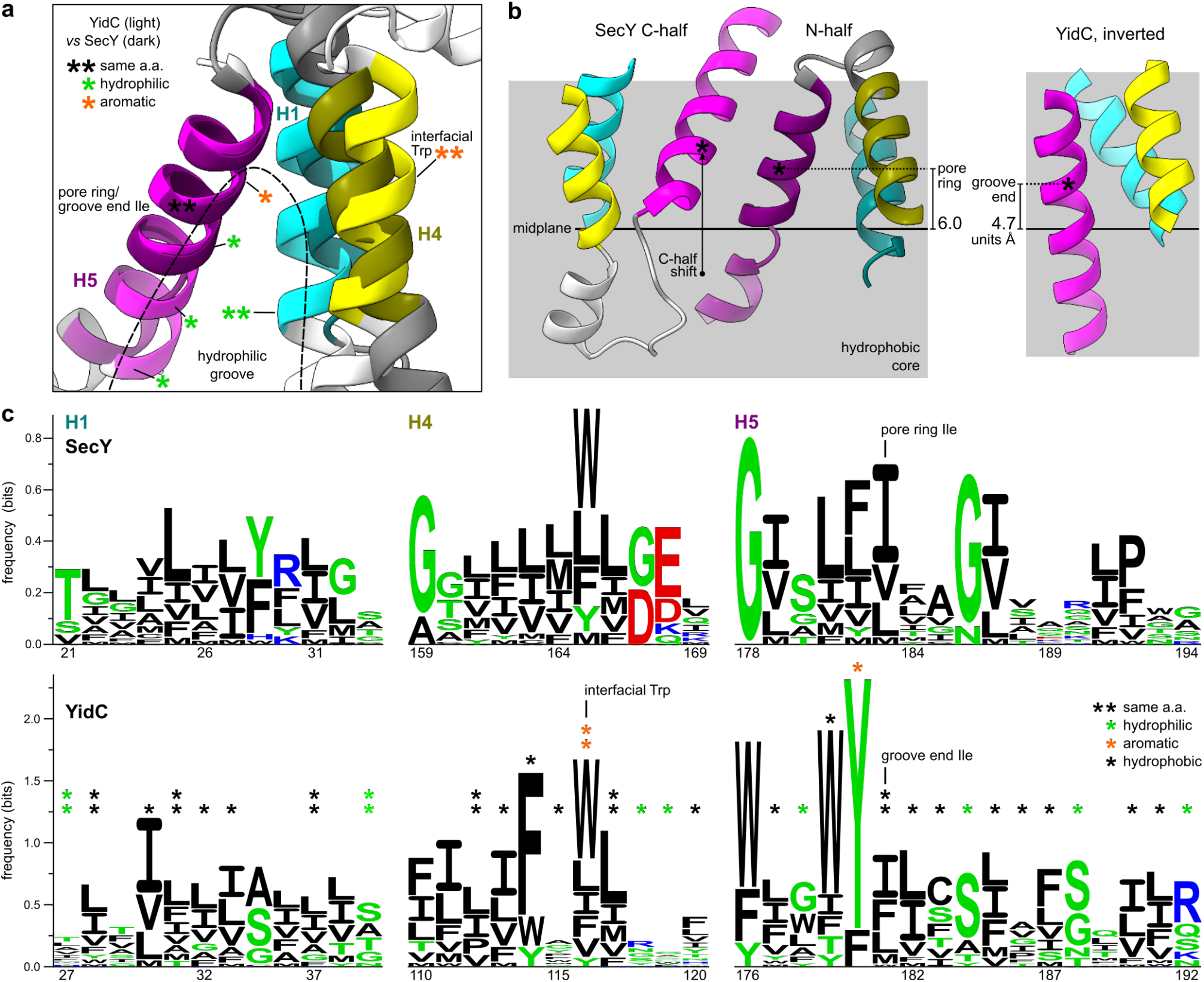
YidC and SecY have similar sequence profiles. **a** Superposition of the SecY N-half (*G. thermodenitrificans*, 6itc) and YidC (*M. jannaschii* MJ0480, 5c8j). Coloured segments correspond to the sequence logos in panel c. Some of the residues which are similar in both families are indicated by asterisks, with lines pointing to the corresponding α-carbon. **b** Transmembrane position of the H5 pore ring or groove end residue in the SecY N-half (*G. thermodenitrificans*, 6itc) and YidC (*B. halodurans* YidC2, 3wo7). The boundaries and midplane of the membrane were determined as described in the Methods. An arrow points from the transmembrane position of the groove end residue in YidC H5 to the pore ring residue in SecY C.H5. **c** Sequence logos for the structurally aligned regions of SecY and archaeal YidC. Column numbers correspond to positions in the proteins modelled in panel a.

Across the 40 structurally aligned sites, YidC and SecY have 22.5% identical consensus sequences, compared to 30.0% between the SecY halves at these same sites. This detailed similarity in both sequence and structure indicates that the residue at the end of YidC’s hydrophilic groove is homologous to the pore ring residue at the end of SecY’s hydrophilic funnel. Hydrophobic interactions between these residues in two antiparallel YidC-like monomers would have favoured dimers with a symmetry that juxtaposed them, allowing them to ultimately form the proto-SecY pore ring.

### SecY’s structural differences from YidC support its unique secretory function

Whereas the conserved cores of SecY and YidC are similar, their differences are concentrated in regions which are hypervariable among the Oxa1 superfamily: h4h5 and H2/3 (Figure 4). H2/3 forms a relatively compact cytoplasmic hairpin in YidC and TMCO1, is markedly elongated and rigid in GET1, and is tethered via long flexible loops in EMC3. By contrast, the H2/3 hairpin in SecY is folded back toward the H1/4/5 bundle and embedded in the membrane.

Despite their differences, H2/3 is a site for substrate signal recognition in both SecY (Figure 7a) and the Oxa1 superfamily. In YidC, TMCO1, and EMC3, the membrane-facing side of H2/3 is thought to interact with substrate TMHs before they reach the hydrophilic groove (Borowska et al., 2015; Kumazaki et al., 2014; McGilvray et al., 2020; Pleiner et al., 2020). In contrast to direct TMH interaction, the rigid and elongated H2/3 coiled coil of GET1 (McDowell et al., 2020) forms a binding site for the substrate targeting factor GET3 (Mariappan et al., 2011; Stefer et al., 2011; Wang et al., 2011). This adaptation may be due to the particularly hydrophobic TMHs inserted by this pathway (Guna and Hegde, 2018), warranting a specialised machinery to shield them in the cytosol.

**Figure 7.**
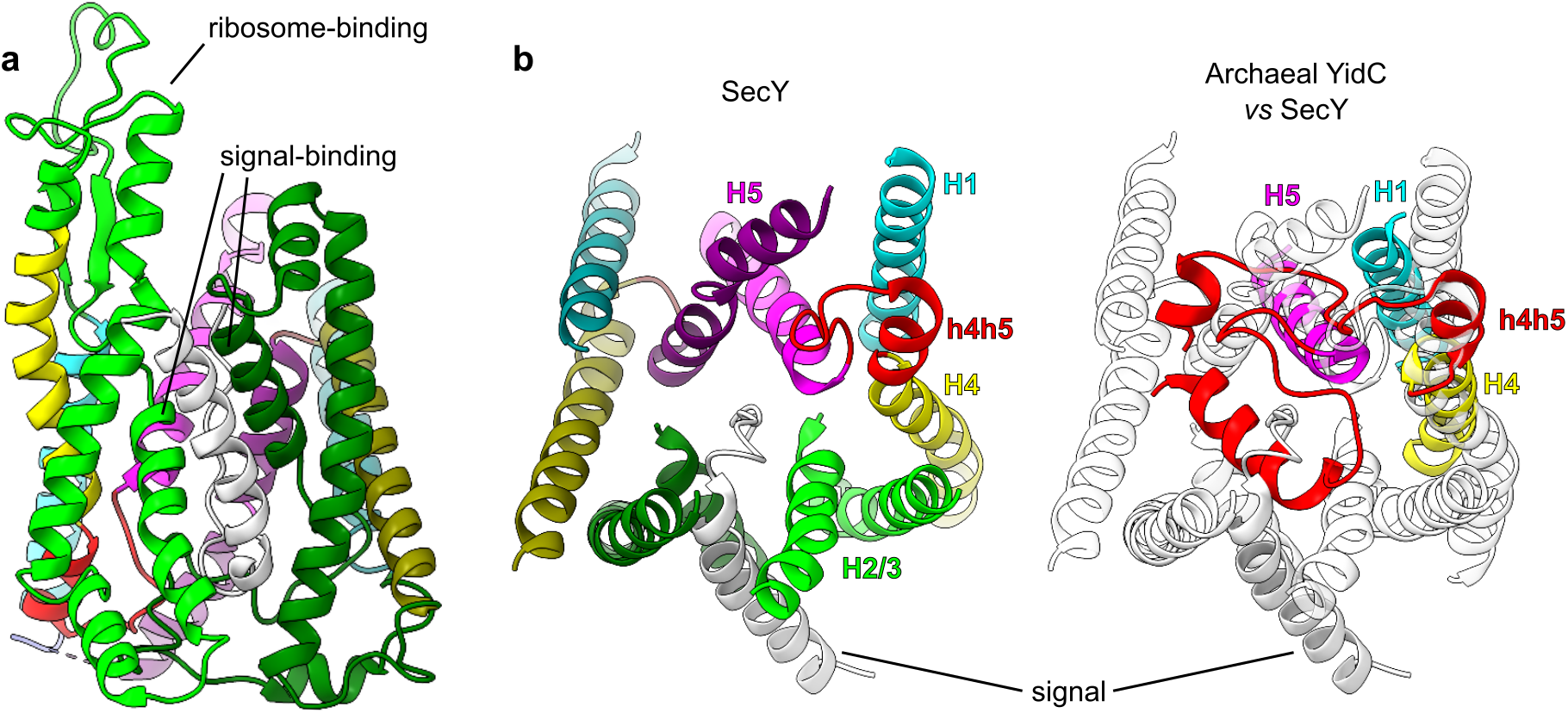
Structural features unique to SecY which enable signal binding and substrate translocation. SecY is *G. thermodenitrificans* SecY/proOmpA (6itc). **a** Signal-binding and ribosome-binding sites on SecY H2/3, viewed laterally. **b** The substrate translocation channel, viewed from its extracytoplasmic side. Only H1-5 and h4h5 of SecY are shown. SecY is colour-coded by consensus element as in Figure 4 (left), or rendered transparent and superimposed by the corresponding elements of archaeal YidC *(M. jannaschii* MJ0480, 5c8j), aligned to the SecY C-half (right).

The migration of H2/3 into the membrane in SecY encloses the translocation channel which in YidC is exposed to the membrane (Figure 7b). This allows SecY to create a more hydrophilic and aqueous environment for its hydrophilic substrates, facilitating their translocation. This is particularly important for SecY’s secretory function, which involves translocating much longer hydrophilic segments than those translocated by YidC.

As a secondary consequence, transmembrane insertion of H2/3 makes the site where signals initiate translocation more proteinaceous and hydrophilic (Figure 5a; Gogala et al., 2014; Park et al., 2014; Plath et al., 1998; Voorhees and Hegde, 2016; Weng et al., 2020). Because of this, translocation via SecY can be initiated via signals which are much less hydrophobic than the TMHs which initiate translocation via YidC (Xie et al., 2007). This, too, is important for SecY’s secretory function, because the signal peptides of secretory proteins are distinguished from TMHs by their relative hydrophilicity (von Heijne, 1985). This biophysical difference allows signal peptidase to specifically recognise and cleave them (Paetzel et al., 2002). Cleavage frees the translocated domain from the membrane to complete secretion.

After H2/3, the next most conspicuous difference between SecY and YidC is in h4h5, which is nearly absent from SecY (Figure 7b). Whereas the H2/3 transmembrane insertion differentiates how SecY and YidC receive and recognise hydrophobic domains, the absence of h4h5 clears the channel through which hydrophilic substrates translocate. As mentioned previously, h4h5 is, like H2/3, hypervariable in the Oxa1 superfamily, forming a peripheral helix in archaea and eukaryotes and a transmembrane hairpin in bacteria. If a more YidC-like h4h5 were present in proto-SecY, proto-SecY dimerisation would place h4h5 inside the hydrophilic groove of the opposite monomer, instead of in contact with the membrane. Thus a YidC-like h4h5 would be selected against in SecY, to maintain a membrane-spanning hydrophilic pore and facilitate translocation.

### Like proto-SecY, YidC uses the distal face of H5 for dimerisation

As shown above, it appears that proto-SecY formed antiparallel homodimers via the distal face of H5 (Figure 8a); here we consider whether this characteristic could have arisen in an ancient member of the Oxa1 superfamily. Antiparallel homodimerisation requires that the monomer possess two characteristics: a tendency to be produced in opposite topologies, and an interface suitable for dimerisation. Although dual topology is not evident in the Oxa1 superfamily, distant ancestors could easily have had this property with relatively few changes. Making only a few changes to basic amino acids (especially lysine and arginine) flanking the first TMH of an IMP can influence its topology, and an inverted first TMH can invert an entire IMP containing several TMHs (Beltzer et al., 1991; Brown et al., 2018; Rapp et al., 2006, 2007). Such changes in topology occur naturally in protein evolution (Rapp et al., 2006; Sääf et al., 1999), and YidC does not contain any conserved basic residues in its soluble segments that would impede this evolutionary process (Figure 4-Figure supplement 4). Moreover, the lysine and arginine bias in extant YidC is no greater than that previously observed in proteins which acquired divergent orientations (Sääf et al., 1999).

**Figure 8.**
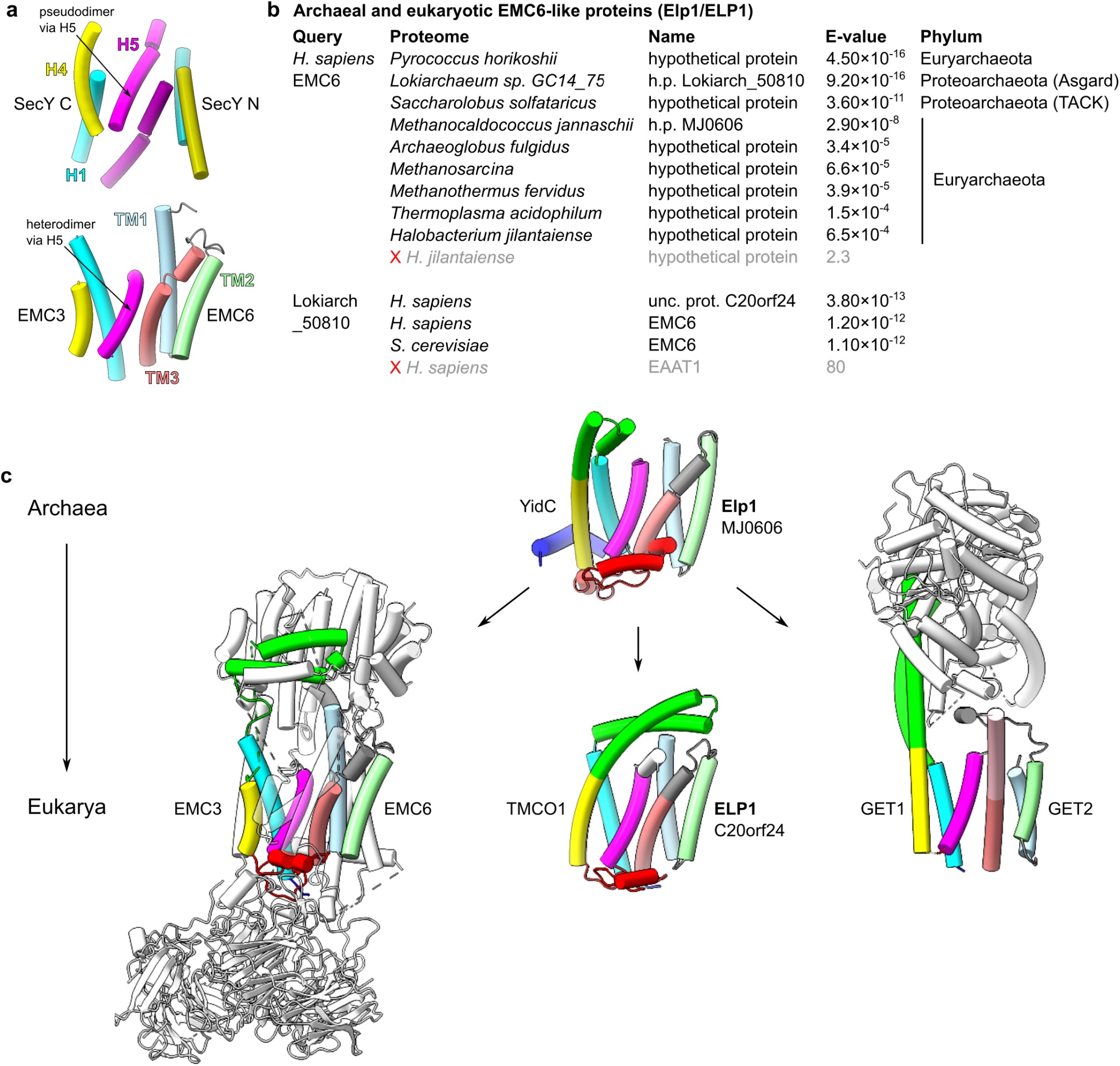
Oxa1 superfamily proteins form dimers via the same interface as proto-SecY. **a** Comparison of the EMC3/6 and proto-SecY dimerisation interfaces. Models show *S. cerevisiae* EMC3 H1/4/5 with EMC6 (6wb9) and *G. thermodenitrificans* SecY H1/4/5 (6itc). **b** Archaeal and eukaryotic HHpred hits. A red cross and grey text indicates the first rejected result. For sequence accession numbers, see Methods. **c** Structural models of the archaeal and eukaryotic heterodimers of YidC and EMC6/GET2 homologs. *S. cerevisiae* EMC (6wb9) and *H. sapiens* GET1/2/3 (6so5) are shown. The N-terminal half of GET2 TM3 which is identified as a sequence insertion by alignment to Elp1 is shown in pink. The *M. jannaschii* YidC/Elp1 (MJ0480/MJ0606) models were predicted by trRosetta and aligned to the EMC3/6 dimer. The *H. sapiens* TMCO1/ELP1 dimer is represented similarly, except the TMCO1 model is from the PDB (6w6l). **Figure 8-Source data 1**. Structure and contact predictions for the EMC6-like proteins. **Figure 8-Figure supplement 1**. Structure and contact prediction for archaeal and human EMC6-like proteins. **Figure 8-Figure supplement 2**. Structural models for nine diverse archaeal EMC6-like proteins.

Unlike dual topology, interactions via the distal face of H5 are known to occur in several Oxa1 superfamily members. This surface forms an intramolecular interaction with the h4h5 transmembrane hairpin in bacterial YidC but remains exposed in archaeal YidC and its eukaryotic homologs (Figure 4). There are no published data on YidC biochemistry in archaeal cells, but eukaryotic EMC3 and GET1 are known to form functionally important complexes, and structural models show that they use the distal face of H5 to do so (Figure 8a; Bai et al., 2020; McDowell et al., 2020; Miller-Vedam et al., 2020; O’Donnell et al., 2020; Pleiner et al., 2020) These interactions via H5 are heterodimeric, rather than homodimeric, but nonetheless demonstrate that EMC3 and GET1 can dimerise (with EMC6 and GET2, respectively) along the same interface as the proto-SecY homodimer without impeding their translocation activities.

To determine whether these H5 interactions are ancient or eukaryote-specific, we queried nine diverse archaeal proteomes for homologs of *H. sapiens* EMC6 or GET2 using HHpred (Zimmermann et al., 2018). Although none displayed significant similarity with GET2, every proteome queried contained exactly one protein similar to EMC6 (Figure 8b). Among these archaeal proteins, those most similar to eukaryotic EMC6 tend to come from the species most closely related to eukaryotes: the Asgard archaean, then the TACK archaean, and then the euryarchaeans. This phylogenetic concordance indicates that the archaeal proteins are homologs of the eukaryotic protein, and that their ubiquity is due to an ancient origin. We provisionally name them Elp1, for EMC6-like protein 1.

Reciprocal queries of *H. sapiens* and *S. cerevisiae* proteomes with the Asgard Elp1 (Lokiarch_50810) identified EMC6 in both cases as high-confidence hits. Unexpectedly, the *H. sapiens* search also identified an even more similar hit, C20orf24 (Figure 8b), which we provisionally name ELP1. These proteins all share a structurally similar three-TMH core (Figure 8-Figure supplement 1a; Figure 8-Figure supplement 2), as evident in both *de novo* and homology-templated structures predicted by trRosetta (Yang et al., 2020).

The patterns of co-evolution between archaeal YidC and Elp1 showed that their highest-probability contacts all occur along the same interface used for dimerisation by eukaryotic EMC3/6 (Figure 8-Figure supplement 1b). This indicates that the distal face of H5 is used for heterodimerisation not only by eukaryotic EMC3 and GET1 but also by archaeal YidC. The third eukaryotic Oxa1 family, TMCO1, displays a similar pattern of co-evolution with ELP1, indicating that they form a similar heterodimer (Figure 8-Figure supplement 1b). TMCO1-ELP1 interaction would be consistent with the absence of ELP1 from *S. cerevisiae* (Figure 8b), because *cerevisiae* also lacks TMCO1.

Although GET2 lacks strong sequence similarity with these EMC6 homologs, its structural similarity with EMC6 was immediately recognised (McDowell et al., 2020; Pleiner et al., 2020). Our identification of archaeal EMC6 homologs reveals a plausible origin for GET2. Consistent with this, although our GET2 query of the lokiarchaean proteome did not identify any very high-similarity proteins, the most similar membrane protein was indeed Elp1 (Lokiarch_50810, HHpred *p* = 0.0057). Moreover, the aligned columns between GET2 and Elp1 correspond exactly to their structurally similar transmembrane domains. The single large gap in this alignment spans the cytoplasmic extension of GET2 TM3, which brings it into contact with GET3 (Figure 8c). Thus, the major difference between GET2 and EMC6 can be explained as a functional adaptation for GET3 recognition, not unlike GET1’s elongation of H2/3.

The absence of a similar heterodimer in bacteria suggests that it may have been acquired in archaea after divergence from bacteria, which instead acquired the H5-occluding transmembrane hairpin in h4h5 (Figure 4). An archaeal origin for Elp1 would be consistent with its genomic location, which is distant from the widely conserved cluster of cenancestral ribosomal genes, SecY and YidC (Makarova et al., 2015). In the period prior to heterodimerisation with Elp1, a YidC homolog could have evolved to use H5 for homodimerisation, giving rise to proto-SecY. YidC’s universal tendency to cover the distal face of H5 supports this possibility.

### Reductive evolution in symbionts demonstrates the functional range of YidC

If proto-SecY originated in the YidC family, YidC might initially have been the cell’s only transporter for the extracytoplasmic parts of IMPs. But some IMPs cannot be integrated by YidC, and instead depend on SecY (Welte et al., 2011). Thus a cell with YidC and not SecY may have been constrained to express a more limited range of IMPs. The looser this constraint, the more plausible it is that such a cell would be viable, and that YidC could have preceded SecY.

Insight into this question of *in vivo* sufficiency can be obtained by inspection of the only cells known to have survived SecY deletion: the mitochondrial symbionts. SecY has been lost from all but one group of eukaryotes for which mitochondrial genome sequences are available, and it has not been observed to relocate to the nuclear genome (Janouškovec et al., 2017). The exceptional group is the jakobids, only a subset of which retain mitochondrial SecY. The incomplete presence of SecY in this group implies that SecY was lost multiple times from the jakobids and their sister groups. SecY deletion is therefore a general tendency of mitochondria, rather than a single deleterious accident.

Mitochondria retain two YidC family proteins, Oxa1 and Oxa2 (Cox18), the genes for which relocated from the mitochondrial genome to the nuclear genome (Bauer et al., 1994; Bonnefoy et al., 1994). As nuclear-encoded mitochondrial proteins, they are translated by cytoplasmic ribosomes and then imported into mitochondria via channels in the inner and outer mitochondrial membranes (Wiedemann and Pfanner, 2017). These channels are essential for the import of nuclear-encoded proteins, but are not known to function in the integration of mitochondrially encoded IMPs (meIMPs), which instead requires export from the matrix, where they are synthesized by mitochondrial ribosomes. This export is generally Oxa1-dependent (Hell et al., 2001).

The meIMPs have diverse properties, including 1 to 19 TMHs and exported parts of various sizes and charges (Figure 9a-c). Oxa1’s sufficiency for their biogenesis *in vivo* is consistent with *in vitro* results showing that *E. coli* YidC is sufficient for the biogenesis of certain 6- and 12-TMH model substrates (Serdiuk et al., 2019; Welte et al., 2011). Ectopically expressed EMC3/6 can rescue meIMP integration in the absence of Oxa1, indicating that Oxa1’s broad substrate spectrum is representative of the Oxa1 superfamily as a whole (Güngör et al., 2021). The only apparent constraint on the meIMPs is that they tend to have only short (∼15 a.a.) soluble segments. This is consistent with observations from *E. coli* that fusing long soluble segments to a YidC-dependent IMP can induce SecY dependence (Andersson and von Heijne, 1993; Kuhn, 1988; Shanmugam et al., 2019). Among the meIMPs, Cox2 is an exception which proves the rule, because Oxa1 cannot efficiently translocate its exceptionally long (∼140 a.a.) C-terminal tail; instead it is translocated by Oxa2 in cooperation with two accessory proteins (Saracco and Fox, 2002).

**Figure 9.**
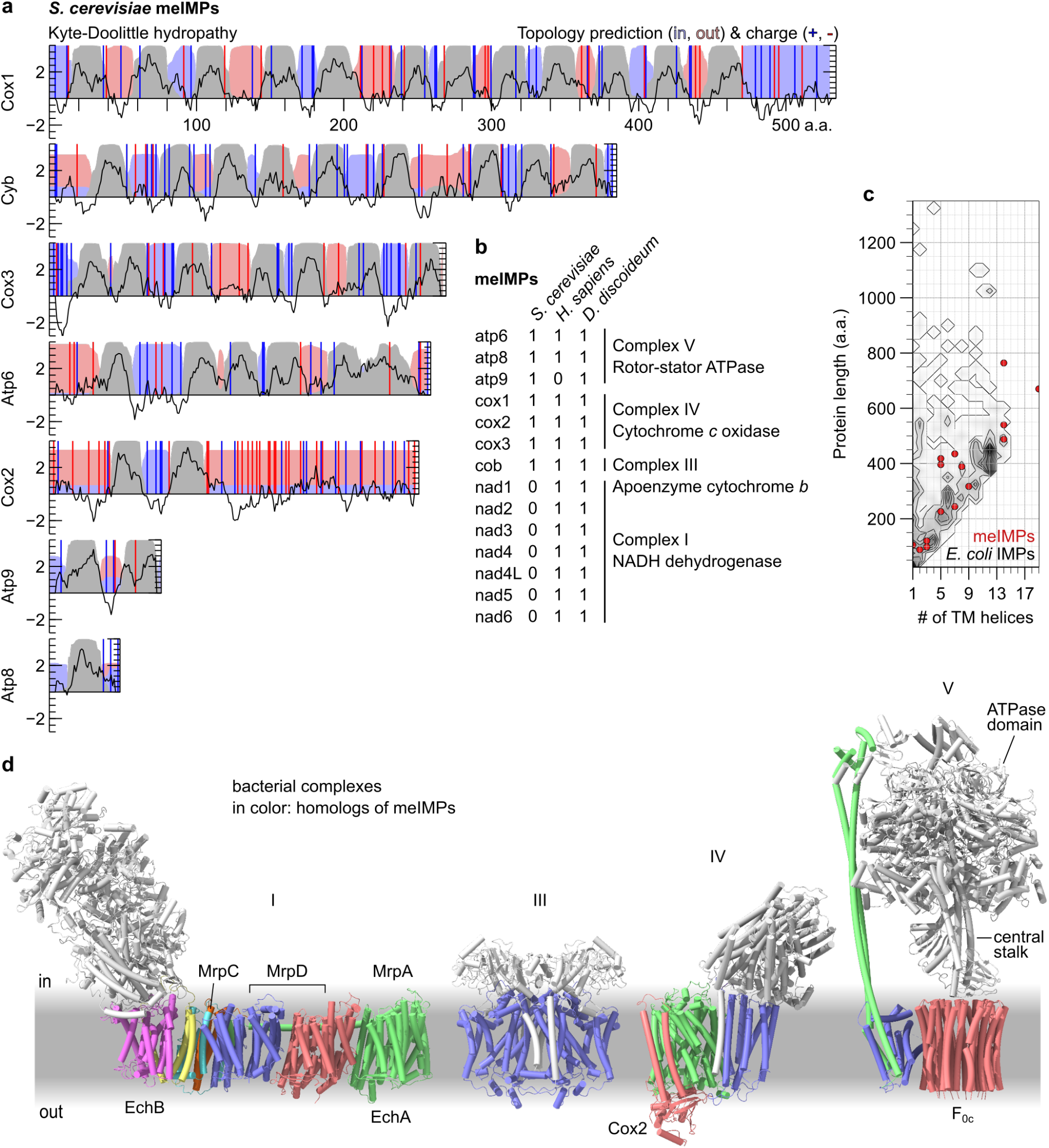
Substrates of the mitochondrial SecY-independent pathway for IMP integration. **a** Sequence characteristics of the mitochondrially-encoded IMPs (meIMPs) from *S. cerevisiae*. Kyte-Doolittle hydropathy (left axis) is averaged over a 9 a.a. moving window (black line). Topology predictions were computed by TMHMM (right axis) to indicate regions which are retained in the mitochondrial matrix (light blue field), inserted into the membrane (grey field), or exported to the intermembrane space (light red field). Positive (blue) and negative (red) residues are marked with vertical bars. **b** Table of all meIMPs in a fungus (*S. cerevisiae*), a metazoan (*H. sapiens*) and an amoebozoan (*Dictyostelium discoideum*). **c** Scatter plot of the length and number of TMHs in the meIMPs of a eukaryote (*D. discoideum*), superimposed on a contour plot and heat-map of all 910 IMPs from a proteobacterium (*E. coli*). Protein lengths were binned in 25 a.a. increments. Each contour represents an increase of 3 proteins per bin. **d** Structures of prokaryotic complexes homologous to meIMPs. Subunits not homologous to the meIMPs listed in panel b are shown in white. Homo-oligomers are represented by a single colour. From left: I, NADH dehydrogenase (*Thermus thermophilus*, 6y11; Gutiérrez-Fernández et al., 2020), III, cytochrome *bc*1, (*Rhodobacter sphaeroides*, 6nhh; Esser et al., 2019), IV, cytochrome *c* oxidase (*R. sphaeroides*, 1m57; Svensson-Ek et al., 2002), V, rotor-stator ATPase (*Bacillus* sp. PS3, 6n2y; Guo et al., 2019). The labelled subunits of NADH dehydrogenase (I) are homologous to the two IMP subunits of the energy-converting hydrogenase (EchA/B), and/or to subunits of the multiple-resistance and pH (Mrp) antiporters. The labelled subunits of IV and V indicate those referenced in the text.

This constraint on soluble segment length is less consequential than it may at first appear, because prokaryotic IMPs in general tend to have only short soluble segments (Figure 9c; Wallin and von Heijne, 1998). Thus, most prokaryotic IMPs may be amenable to SecY-independent, YidC-dependent biogenesis. Consistent with this, in *E. coli*, the signal recognition particle (SRP) has been found to target nascent IMPs to either SecY or YidC (Welte et al., 2011), and YidC is present at a concentration 1–2*×* that of SecY (Schmidt et al., 2016). By contrast, IMPs with large translocated domains became much more common in eukaryotes (Wallin and von Heijne, 1998) concomitant with YidC’s divergence into three niche paralogs, none of which are essential at the single-cell level (Guna et al., 2018; Jonikas et al., 2009; McGilvray et al., 2020).

Even without extrapolating from the meIMPs to other similar IMPs, it is clear that chemiosmotic complexes are amenable to YidC-dependent, SecY-independent biogenesis (Figure 9d). These complexes couple chemical reactions to the transfer of ions across the membrane and are sufficient for the membrane’s core bioenergetic function. Although the complexes shown participate in aerobic metabolism, which presumably post-dates the oxygenation of Earth’s atmosphere, they have homologs which enable chemiosmosis in anaerobes. In particular, chemiosmosis in methanogens and acetogens employs the rotor-stator ATPase, Mrp antiporters, and an energy-converting hydrogenase (Ech; Lane and Martin, 2012), all of which have homologs of their IMP subunits among the meIMPs (Figure 9d) and may have participated in primordial anaerobic metabolism (Weiss et al., 2016).

Thus if YidC had preceded SecY, it would have been sufficient for the biogenesis of diverse and important IMPs, but likely not the translocation of large soluble domains. This is supported by the results of reductive evolution in chloroplasts, which retain both SecY (cpSecY) and YidC (Alb3) (Xu et al., 2020). cpSecY imports soluble proteins across the chloroplast’s third, innermost membrane, the thylakoid membrane (Peltier et al., 2002). This thylakoid membrane was originally part of the chloroplast inner membrane (equivalent to the bacterial plasma membrane), much like the mitochondrial cristae, but subsequently detached and now forms a separate compartment (Vothknecht and Westhoff, 2001). Because the thylakoid membrane is derived from the plasma membrane, import across the thylakoid membrane is homologous to secretion across the plasma membrane. Thus, when symbiosis removed the need for secretion, SecY was eliminated from mitochondria, whereas it was retained in chloroplasts for an internal function homologous to secretion.

A primordial YidC-dependent cell may simply not have secreted protein or may instead have used a different secretion system. Notably one primordial protein secretion system has been proposed: a protein translocase homologous to the rotor-stator ATPases (Mulkidjanian et al., 2007). Translocases are transporters which use chemical reactions to drive translocation (Tipton, 2018), such as the translocase formed when the SecA ATPase acts in tandem with the SecYEG channel (Erlandson et al., 2008). The putative rotor-stator-like protein translocase used its ATPase subunit to unfold and feed substrates through the homo-oligomeric channel formed by F_0c_, now occupied by the central stalk (Figure 9d). The strict YidC-dependence of F_0c_ biogenesis in *E. coli* (Yi et al., 2003) hints that YidC and F_0c_ shared an early era of co-evolution, as a laterally closed channel for the secretion of soluble proteins (F_0c_) and a laterally open channel for the integration of membrane proteins (YidC), including F_0c_ itself. The subsequent advent of a laterally gated channel, SecY, would have facilitated the biogenesis of a hybrid class of proteins: IMPs with large translocated domains.

## Discussion

By comparing structures of the SecY N- and C-halves, we identified a maximum-symmetry pair, and thus an estimate of the structure of their last common ancestor, proto-SecY. Their alignment identifies homologous sites in each half, revealing that both the hydrophobic pore ring and the interface between halves are symmetric. The conservation of these features indicates that they were also present in proto-SecY, and thus that it formed antiparallel homodimers and functioned as a protein-conducting channel.

In automated database searches for structures similar to SecY’s halves, the top hit is the Oxa1 superfamily, of which YidC is the prokaryotic member. The SecY/Oxa1 fold consists of a right-handed three-helix bundle (H1/4/5), interrupted after the first helix by a helical hairpin (H2/3) and prefixed by an N-terminal peripheral helix (H0) that abuts H4. The H2/3 hairpin is cytoplasmic in the Oxa1 superfamily but transmembrane in SecY, where it forms the lateral gate helices. This suggests that H2/3 was originally cytoplasmic, then evolved to pack against the surface of H1/4/5 in the proto-SecY stem lineage. This sequence of events would explain the peculiar non-sequential packing arrangement of SecY’s transmembrane helices.

This unexpected correspondence motivates a re-evaluation of the literature on SecY and YidC. In both, H1/4/5 buries a hydrophilic groove inside the membrane to facilitate the translocation of hydrophilic polypeptide. Juxtaposing two grooves, one on each side of the membrane, allows SecY to open a membrane-spanning pore, whereas YidC has only a cytoplasmic groove. Structural alignments superimpose the hydrophobic residue in H5 that rings the SecY pore onto the hydrophobic residue that ends the YidC groove, and likewise the surrounding polar and aromatic groove residues. Both SecY and YidC recognise hydrophobic helices in their substrates via binding at the protein-lipid interface, and in doing so induce a hairpin conformation in the substrate’s hydrophilic flank which initiates its translocation. The SecY-specific lateral gate helices create a more hydrophilic environment for signal recognition and substrate translocation that is better suited to SecY’s specific secretory function.

Whereas proto-SecY formed homodimers via the distal face of H5, two of the three eukaryotic Oxa1 member families are known to use this interface for heterodimerisation. Homology would predict that this is an ancient tendency. We indeed found indications that H5-mediated heterodimers are formed by the third eukaryotic Oxa1 superfamily member, TMCO1, and by archaeal YidC. In bacterial YidC this interface instead makes intramolecular contacts with bacteria-specific TMHs. To gauge the plausibility of a YidC-dependent, SecY-independent primordial cell, we reviewed the range of substrates translocated by YidC in SecY-lacking mitochondria, and found that it spans most of the diversity of the prokaryotic membrane proteome. The surprising conclusion of our study is that a YidC homolog could have both preceded and evolved into proto-SecY, whose gene duplication and fusion then originated the present-day SecY family.

### Evaluation of the homology hypothesis

It is important to consider whether the similarities between SecY and YidC could arise by convergent evolution under shared constraints (making them analogs), rather than divergent evolution from a common ancestor (making them homologs). Deciding between the analogy hypothesis and the homology hypothesis requires an assessment of whether any plausible constraints could explain their similarities (Doolittle, 1994). We will weigh their functional, mechanistic, structural, and sequence similarities in turn.

Laterally open helical protein-conducting channels have arisen by functional convergence several times (Figure 10). Thus if the similarity between SecY and YidC were solely functional, the analogy hypothesis would be attractive. Analogy would also be plausible if the similarity between SecY and YidC were solely mechanistic, because their mechanism is common among amphiphile transporters. Many use a membrane-exposed hydrophilic groove to translocate the hydrophilic parts of an amphiphile while exposing its hydrophobic parts to the bilayer (Bakelar et al., 2016; Brunner et al., 2014; McKenna et al., 2020). Moreover the hairpin conformation which protein transporters induce in their substrates is a predictable result of physical constraints which disfavour head-first translocation (Engelman and Steitz, 1981).

**Figure 10.**
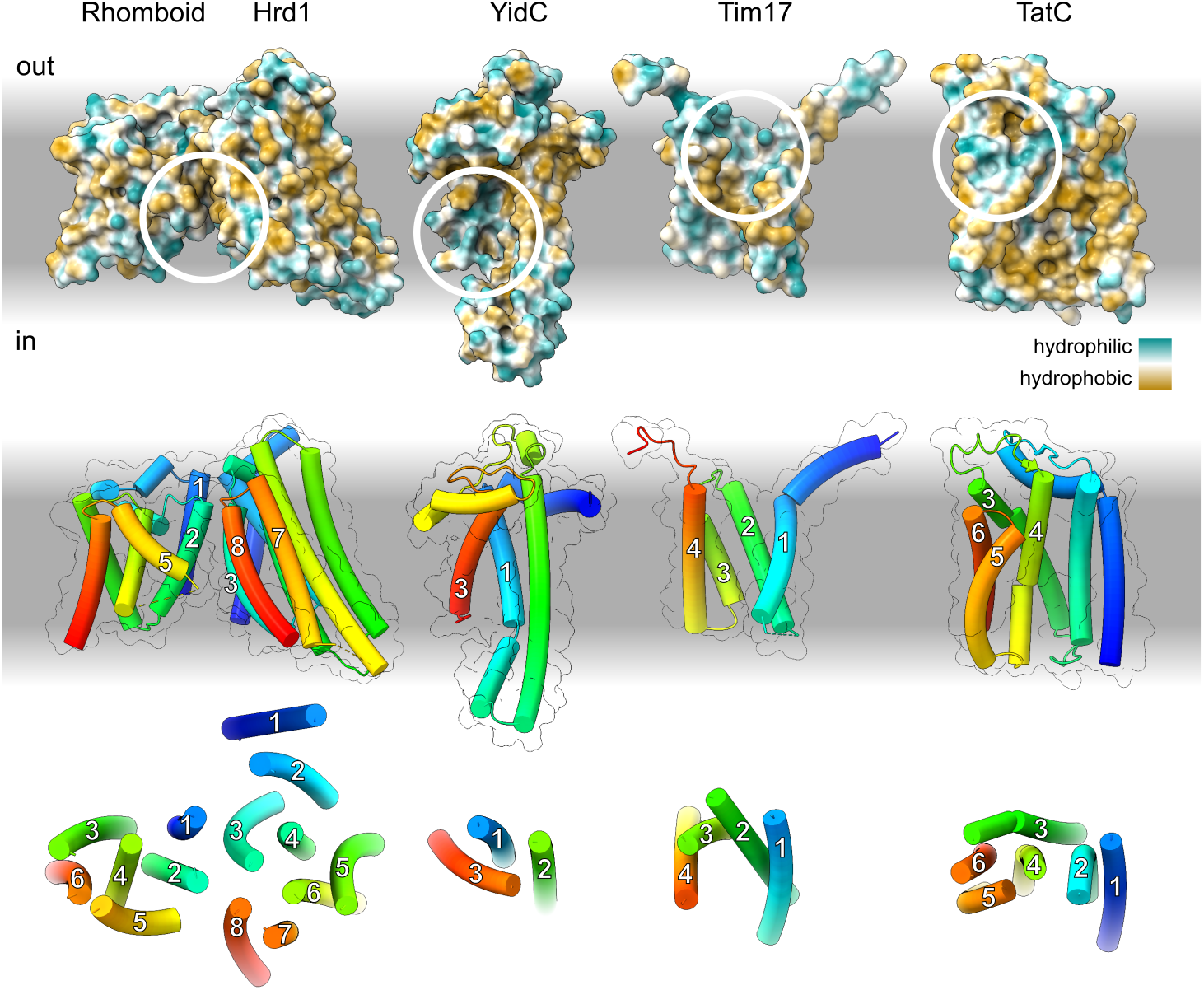
Structures of the known families of laterally open helical protein-conducting channels. Top: Structural models shown as solvent-excluded surfaces colour-coded by hydropathy. The hydropathy of the lipidic and aqueous phases represented on a separate scale, ranging from hydrophilic (white) to hydrophobic (grey). White circles indicate intramembrane hydrophilic grooves. Middle: models shown as tubes colour-coded by position. Transmembrane segments in the vicinity of the hydrophilic groove are numbered. Bottom: Axial views of each molecule showing only transmembrane helices. From left to right, the models representing each family are as follows. Rhomboid: *S. cerevisiae* Der1 (6vjz), Hrd1: *S. cerevisiae* Hrd1 (6vjz), YidC: *M. jannaschii* MJ0480 (5c8j, extended by structure prediction to include the originally unmodelled H2/3; see Figure 4-Figure supplement 4), Tim17: *S. cerevisiae* Tim22 (6lo8, Zhang et al., 2020), TatC: *Aquifex aeolicus* TatC (4b4a, Rollauer et al., 2012).

Thus there is precedent and a clear physical basis for SecY and YidC’s functional and mechanistic similarities arising by convergence. But the same is not true of their structural similarities. First, it would be unprecedented for structural similarity to arise by convergence within this functional and mechanistic class, given that all other known amphiphile transporters are grossly dissimilar from one another, including all other laterally open helical protein-conducting channels (Figure 10). This suggests that the space of mechanically sufficient folds is large, and thus the likelihood of convergence low.

Second, the extensive literature on SecY and YidC discussed throughout this paper suggests no physical reason why their mechanism would favour the SecY/Oxa1 fold. Thus attributing their structural similarity to mechanistic constraints would require one to assume that such a constraint exists. On the contrary, structural convergence due to mechanistic constraints typically occurs in only those parts of a protein with clear mechanistic roles, such as the catalytic dyads and triads of enzymes. For example, a comprehensive survey of convergence in analogous enzymes identified 267 pairs with similar dyads or triads, but none with similar folds (Gherardini et al., 2007). Fold space is evidently large enough that many folds are likely to be compatible with a given mechanism.

Perhaps the most extensive known case of structural convergence in functionally similar helical IMPs occurred among thiol oxidoreductases. Four analogous families all use four-helix bundles to bind their redox cofactors, despite two being IMPs and two being cytoplasmic (Li et al., 2018). But they are nonetheless easily distinguishable because they connect those four helices in different orders. This indicates that even an exceptionally tight constraint on the architecture of secondary structure elements does not comparably constrain the connectivity of those elements. Indeed, the seven TMHs of another IMP, rhodopsin, can be experimentally permuted while retaining activity (Mackin et al., 2014). 36 such permutations are possible for proto-SecY H0-5. Although some permutations would be more likely to evolve than others, analogy would be as likely as homology only if all 35 other permutations were forbidden. Thus even if the specific architectures of proto-SecY and YidC were favoured by some yet unknown mechanistic constraint, their identical connectivity would still weigh in favour of the homology hypothesis.

Without functional or mechanistic constraints, structural convergence can still occur in some cases due to folding constraints imposed by the intrinsic properties of polypeptide and solvent. One would expect such intrinsically preferred structures to occur frequently and in functionally unrelated contexts. For this reason the phylogeny of ubiquitous and functionally diverse folds is challenging to discern (Lupas et al., 2001). But it is implausible that folding constraints strongly favour the SecY/Oxa1 fold because it is not found in other proteins, as our database queries show.

Finally, we consider the most detailed similarity between SecY and YidC, which is in their H1/4/5 sequence profiles. If this bundle was in the same transmembrane position and orientation in both proteins, one might imagine that their sequence similarity was a product of mechanistic constraints. However, this similarity occurs despite topological inversion (Figure 6), and thus lends at least some weight to homology. Just how how much weight is unclear. Ideally one would compare SecY and YidC to analogous proteins with the same structure and function and see how exceptional their sequence similarity is among that set. Such a test is partly feasible for proteins with very common folds, like β-barrels (Remmert et al., 2010), but impossible here, because our database queries find no other proteins with the SecY/Oxa1 fold.

In sum, the dispositive evidence for homology between SecY and the Oxa1 superfamily is structural. It would be empirically unprecedented and theoretically improbable for their structural similarity to arise by convergence. We therefore conclude that they are more likely to be homologs than analogs, and describe them as homologs hereafter.

### Implications for the evolution of protein transport

Besides illuminating SecY’s origins, identifying YidC as its progenitor implies that YidC is the oldest known channel. This has implications for the evolution of IMPs generally, including YidC itself, and other ancient components of the general secretory pathway: SecEG (Cao and Saier, 2003; Kinch et al., 2002; Figure 11-Figure supplement 1), signal peptidase (Rawlings and Bateman, 2019), SRP and SRP receptor (SR; Gribaldo and Cammarano, 1998). We propose that the following stepwise model (Figure 11) is the simplest that is consistent with the available data.

**Figure 11.**
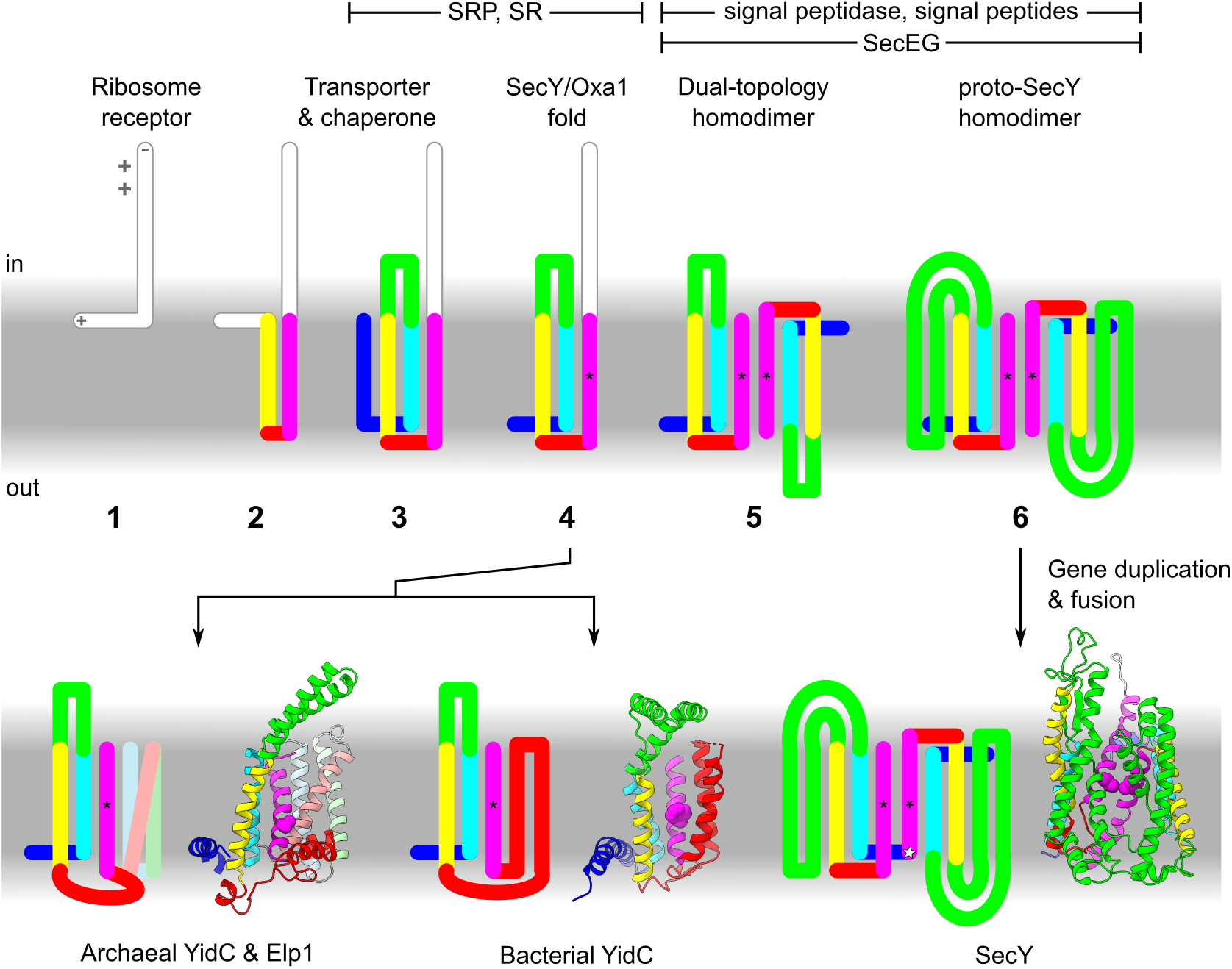
Model for the evolution of YidC and SecY. Charged side chains and termini are indicated only at stage 1, by grey symbols. Asterisks indicate the pore ring or groove end residue in H5. At top, additional components of the secretory pathway label a range of stages at which they they may have arisen. Models show archaeal YidC and Elp1 (*M. jannaschii* MJ0480 and MJ0606, trRosetta), bacterial YidC (*B. halodurans* YidC2, 3wo6), and SecY (*G. thermodenitrificans* SecY, 6itc). **Figure 11-Figure supplement 1**. Similarity between archaeal and bacterial SecG.

Step 1. An ancestor of YidC was a membrane-peripheral ribosome receptor. This is parsimonious insofar as both YidC and SecY are ribosome receptors, and like all IMPs presumably descend from peripheral proteins (Mulkidjanian et al., 2009). Ribosome receptor function can be achieved with just two low-complexity domains: a weakly hydrophobic anchor and a polybasic extension. This receptor would reduce aggregation of hydrophobic domains in the aqueous phase by creating a population of membrane-bound ribosomes, from which any nascent IMPs would be more likely to encounter the membrane. Similar polybasic C-terminal tails are known to occur in YidC and can compensate for deletion of SRP or SR (Seitl et al., 2014; Szyrach et al., 2003).

Step 2. The peripheral helix acquires a transmembrane hairpin, thereby integrating into the membrane. Uncatalyzed insertion of a hairpin is more efficient than that of a single TMH (Engelman and Steitz, 1981), making a hairpin the more likely initial membrane anchor. We infer that SRP/SR-dependent targeting did not evolve until after this and other minimal IMPs existed for it to target. The proximity of this hairpin to nascent IMPs emerging from the bound ribosome imposes a selective pressure on the hairpin to evolve membrane-buried hydrophilic residues that can facilitate IMP integration. Substrates would engage this YidC ancestor in the same hairpin conformation that is favoured during uncatalysed translocation, and this conformation remains how substrates engage SecY and YidC today.

Step 3. Acquisition of a second transmembrane hairpin produces a four-TMH protein containing the conserved three-helix bundle and hydrophilic groove. The segment between the first and second transmembrane hairpins becomes the cytoplasmic hairpin H2/3. The additional TMHs allow YidC to form a hydrophilic groove in the membrane, thereby further facilitating substrate translocation.

Step 4. The hydrophilic groove allows hydrophilic termini to efficiently translocate, including the N-terminus of the YidC ancestor itself, which acquires a new position as the extracytoplasmic peripheral helix H0. Thus the full SecY/Oxa1 fold is now attained. By this time, SRP/SR have evolved, and H2/3 evolves interactions with SR and the ribosome, features that are still evident in SecY, YidC, and TMCO1 (Kuhn et al., 2011; McGilvray et al., 2020; Petriman et al., 2018). At this stage, the YidC gene duplicated, allowing one paralog to seed the SecY lineage. Paralogous origin in a tandem duplication event would be consistent with the commonly observed juxtaposition of YidC and SecY in prokaryotic genomes (Makarova et al., 2015).

Step 5. The original ribosome-binding tail is lost due to its redundancy with SRP/SR for targeting and H2/3 for docking. Loss of this element and genetic drift yields a subpopulation of inverted proteins. Antiparallel dimerisation of the two subpopulations would be favoured because the monomers prefer a similarly thinned membrane, especially near the distal face of H5.

Hydrophobic interactions between the groove end residues would favour the particular dimer symmetry of SecY, which juxtaposes them. The non-SecY lineage of YidC (from step 4) evolves in archaea to heterodimerise with Elp1 via the distal face of H5; in bacteria, this same surface becomes covered by the h4h5 transmembrane hairpin.

Antiparallel homodimerisation in the SecY lineage positions hydrophilic grooves on both sides of the membrane, leaving at most a thin hydrophobic layer between them, as in the heterodimeric ERAD channel (Wu et al., 2020). This facilitates the translocation of IMPs with large soluble domains, including signal peptidase. In the presence of signal peptidase, signal-dependent secretion becomes possible, with the first cleavable signal peptides being the TMHs of IMPs which had previously anchored their now-secreted extracytoplasmic domains. Signal peptides originating as TMHs would explain why both engage SecY in a similar way.

At this stage or later, SecEG are acquired. SecE’s symmetrical binding to each half of the dimer would stabilise it, particularly when the monomers separate to accommodate substrates. Evolution of SecEG after YidC but before proto-SecY is consistent with evidence that their integration depends on other YidC homologs apart from SecY (Guna et al., 2018; Yi et al., 2003).

Step 6. Transmembrane insertion of H2/3 creates a lateral gate, and thus the proto-SecY fold. By inserting between the hydrophilic grooves and the membrane, H2/3 makes those grooves deeper and more hydrophilic, further facilitating translocation. As a secondary consequence, it also creates a more hydrophilic site for signal recognition. This allows cleavable signal peptides to become less hydrophobic than TMHs, and thus more easily distinguished by signal peptidase.

Duplication and fusion of the proto-SecY gene would allow each half of this initially symmetric protein to specialise for cytoplasmic and extracytoplasmic functions. For example, the C.h1h2 and C.h4h5 loops would continue to bind ribosomes, whereas these same loops in the N-half atrophy. One such loop was repurposed as the plug. We infer that gene duplication occurred after antiparallel dimerisation because this has precedent (Lolkema et al., 2008; Rapp et al., 2006) and because both halves of SecY conserve the transmembrane insertion of H2/3, which appears to be an adaptation to antiparallel dimerisation.

### Outlook

One might hope that the increasing diversity of known IMP structures will reveal the origins of other pseudosymmetric channels, which have been refractory to sequence searches (Hennerdal et al., 2010). But the detectability of SecY’s origins may be due to the unusual properties of protein as a transport substrate. Unlike most substrates, protein can be sufficiently hydrophobic to assist in its own translocation, making a partial channel like YidC functionally sufficient. Moreover YidC is thought to serve a second function as a chaperone for IMP folding, which makes it non-redundant to SecY. The same hydrophilic groove used for transport is thought to mediate this chaperone function (Kumazaki et al., 2014; Nagamori et al., 2004; Serdiuk et al., 2016). Other pre-fusion channel precursors may have exposed similar grooves for transport, but this non-redundant chaperone function is unique to protein substrates. Thus pre-fusion homologs of other channels may have had less reason to be conserved.

Although theories about early evolutionary transitions are not experimentally testable, experimental reconstructions can at least demonstrate their plausibility. Efforts to reconstruct the earliest cells, called protocells, could capitalise on the synergy detailed above between YidC and the putative rotor-stator-like protein-secreting translocase (Mulkidjanian et al., 2007). This protein translocase is itself thought to descend from an RNA translocase, in part because its ATPase domain descends from an RNA helicase. By facilitating the integration of such an RNA translocase, YidC would have indirectly facilitated gene transfer among protocells, thereby allowing recombination to continue despite cellularisation and accelerating this stage of evolution.

Since early studies on protein transport, it has been theorised that protocells were preceded by inside-out precursors, called obcells, which arose when macromolecules colonised the surface of a vesicle (Blobel, 1980). The obcell’s interior would then become the protocell’s periplasm after an involution akin to gastrulation. This stage would be the earliest that could have hosted protein transporters, but may have featured only a rudimentary genetic code (Cavalier-Smith, 2001).

Consistent with such an early origin, the conserved pore and groove residues identified here (proline, glycine, serine, branched aliphatics) are all abiotically generated (Weber and Miller, 1981) and thought to be among the first encoded (Trifonov, 2000). Moreover even the simplest ancestors of YidC modelled here served functions that would be useful during the colonisation of a membrane. Thus this stage is a reasonable early bound for the origin of YidC. More precise estimates may require more detailed contextual knowledge about protocells and their precursors.

## Supporting information

Figure 2-Source data 1. Structure-guided alignment of the SecY halves

Figure 3-Source data 1. Results from Dali queries of the PDB25

Figure 4-Source data 1. Predicted structures of archaeal YidC

Figure 8-Source data 1. Structure and contact prediction for archaeal and human EMC6-like proteins

## Acknowledgements

A.J.O.L. is funded by the UK Medical Research Council and the Cambridge Commonwealth, European and International Trust. R.S.H is funded by the UK Medical Research Council (MC_ UP_A022_1007). For critically reading the manuscript, we thank Robert J Keenan, John P O’Donnell, and Viswanathan Chandrasekaran.

## Methods

### Sequence similarity measures, datasets and queries

SecY sequence analyses used a recently published dataset of taxonomically diverse prokaryotic sequences (Harris and Goldman, 2021). To this dataset, we added the sequences for two structurally characterised SecY (*G. thermodenitrificans* and *M. jannaschii*) and removed 8 fully redundant sequences, 3 highly divergent Elusimicrobia sequences, and 4 N-terminally truncated sequences.

The resulting alignment contains 342 sequences, 263 bacterial and 79 archaeal. Sequences were aligned with MAFFT L-INS-i. This and subsequent MAFFT alignments used default parameters, except for using the alternative gap extension penalty --ep 0.123 that is standard for sequences without domain-scale indels.

Pairwise sequence identities within groups of sequences were calculated by re-alignment with ClustalOmega (Sievers and Higgins, 2021) on [the European Bioinformatics Institute server](https://www.ebi.ac.uk/Tools/msa/clustalo/) (Madeira et al., 2019). Clustal reports an all-against-all identity matrix and has previously been used to quantify long-term evolutionary trends in sequence identity (Konaté et al., 2019). Default parameters were used. The number of pairwise comparisons was 342^2^ for the SecY halves, 89×75 for the ComEA and UvrC (HhH)_2_ families, and 79×263 for archaeal and bacterial SecY. Sequences were shuffled to estimate excess identity using the [Sequence Manipulation Suite](https://www.bioinformatics.org/sms2/index.html) (Stothard, 2000).

HHpred pairwise comparisons and database queries used [the Max Planck Institute for Developmental Biology's server](https://toolkit.tuebingen.mpg.de/tools/hhpred) (Zimmermann et al., 2018). All used default parameters. The HHpred *p* between the full-length SecY halves was calculated using their subsequences from *M. jannaschii* SecY as input for automatic MSA generation. Database queries pertaining to EMC6/GET2 homologs used the *H. sapiens* EMC6 (NP_001014764.1), GET2 (NP_001736.1), or *Lokiarchaeum sp. GC14_75* Lokiarch_50810 (KKK40543.1) sequence.

The N- and C-half sequences were aligned using the structurally similar regions of H1-5 as a seed alignment, to which the 684 N- and C-half sequences were added using MAFFT L-INS-i --seed. This alignment of halves was used as input to [ConSurf](https://consurf.tau.ac.il/) (Ashkenazy et al., 2016) to score the conservation of each column across the two halves. Conservation scores for *B. halodurans* YidC2 were obtained from [ConSurf-DB](https://consurfdb.tau.ac.il/) (Chorin et al., 2020).

*E. coli* IMP annotations and sequences were fetched from UniProt (The UniProt Consortium, 2019). The sequences for proteins annotated as multi-pass IMPs and not beta-stranded were filtered at 25% identity using MMseqs2 (Steinegger and Söding, 2017), yielding 554 sequences.

Archaeal YidC sequences were collected from Pfam family PF01956. All UniProt sequences assigned to PF01956 were retrieved, and non-archaeal sequences (EMC3 and TMCO1) were excluded. To speed subsequent alignment, the archaeal sequences were filtered at 80% sequence identity using MMseqs2, in target-coverage mode so as to preferentially eliminate fragments. The resulting 871 sequences were aligned by MAFFT L-INS-i. Sequence logos were computed for columns from this alignment and the SecY alignment using DTU Health Tech’s Seq2Logo 2.0 server (Thomsen and Nielsen, 2012) with default parameters.

### Structural similarity measures and database queries

Each SecY model was split into N- and C-halves at an arbitrary point in the poorly conserved loop between them close to the C-terminus of N.H5. The resulting half-SecY structures were multiply aligned and compared by TM-score using the Zhang group’s mTM-align server (Dong et al., 2018b, 2018a), which also reports their number of common core a.a. and RMSD.

Structural searches of the PDB25 were performed using the Holm group’s [Dali server](http://ekhidna2.biocenter.helsinki.fi/dali/) (Holm, 2020). The Dali PDB25 is a subset filtered at 25% maximum pairwise sequence identity, and excludes some additional structures, including TMCO1 (6w6l), due to file format incompatibilities. It contained 21390 chains when queried. Results were manually reviewed to exclude hits with SecY proteins or regions that are not transmembrane. Dali *Z*-scores were equated to *p-*values by assuming an extreme value distribution of scores as in Sierk and Pearson, 2004,

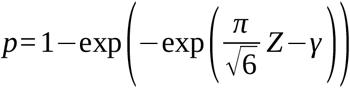

where *γ* is Euler’s constant.

The number of possible connectivities consistent with the architecture of proto-SecY was counted combinatorially, (N_in_ TMHs)!×(N_out_ TMHs)!×(N_in_ TMHs to which H0 could be prepended) = 3!×2! ×3 = 36.

### Co-evolutionary structure and contact prediction

Structural models were predicted from amino acid sequences using the trRosetta algorithm (Yang et al., 2020). End-to-end pipelines which automate the intermediate steps of multiple-sequence alignment generation and homology template selection were used, which reduces user input to only the single protein sequence of interest. [The Baker lab's server](https://robetta.bakerlab.org/) (for Ylp1) and [the Yang lab's server](https://yanglab.nankai.edu.cn/trRosetta/) (for the EMC6 family) were used.

NCBI RefSeq accession numbers for the EMC6/GET2 homolog sequences used as queries are as follows: *P. horikoshii* WP_010885465.1, *Lokiarchaeum sp. GC14_75* KKK40543.1, *S. solfataricus* WP_009990433.1, *M. jannaschii* WP_010870110.1, *A. fulgidus* WP_010878056.1, *Methanosarcina* WP_011032380.1, *M. fervidus* WP_013413780.1, *T. acidophilum* WP_010900743.1, *H. jilantaiense* WP_089668789.1, *H. sapiens* NP_061328.1 (C20orf24 isoform a, a.k.a. UniParc isoform 2, Q9BUV8-2). For the sequences most similar to eukaryotic EMC6, yeast and human EMC6 were automatically selected as homology templates (PDB 6wb9, 6z3w), as indicated in Figure 8-Figure supplement 2a.

Heterodimeric contacts were predicted from amino acid sequences using the RaptorX ComplexContact algorithm (Zeng et al., 2018) as provided by [the Xu group's web server](http://raptorx.uchicago.edu/ComplexContact/). The multiple-sequence alignments generated by RaptorX ComplexContact for MJ0606 and C20orf24 were reviewed to ensure that they did not include any proteins annotated as EMC6.

### Figure preparation

All models were aligned and rendered in UCSF ChimeraX (Pettersen et al., 2020). Surface hydrophobicity was computed in ChimeraX by its default method: pyMLP (Broto et al., 1984; Laguerre et al., 1997) with Fauchere propagation and lipophilicity values from Ghose et al., 1998. Models depicted relative to a membrane are positioned and oriented according to the prediction algorithm provided by [the Orientations of Proteins in Membranes server](https://opm.phar.umich.edu/) (Lomize et al., 2012).

OPM does not account for any anisotropy which lipid-exposed hydrophilic residues may induce, and thus none is shown. The OPM-predicted midplane for YidC was adjusted 1.8 Å toward the cytoplasm to agree with molecular dynamics simulations in which a conserved arginine in H1 (homologous to *B. halodurans* YidC2 R72) sits at the bilayer midplane (Chen et al., 2017). The membrane’s interfacial layers are shown as linear gradients half the width the hydrophobic layer, to approximate experimentally determined polarity profiles (White and Wimley, 1999).

Per-residue hydropathy and charge were computed from protein sequences using [EMBOSS Pepinfo](https://www.ebi.ac.uk/Tools/seqstats/emboss_pepinfo/) (Madeira et al., 2019), topology predicted using [TMHMM](https://services.healthtech.dtu.dk/service.php?TMHMM-2.0) (Krogh et al., 2001), and plotted in [Veusz](https://veusz.github.io/). The 2-D histogram of IMP length *vs* TMH count was likewise plotted in Veusz.

## Supplementary Note

We sought to estimate the amino acid sequences of cenancestral SecY and proto-SecY. A recent study of another internally duplicated cenancestral protein, the helix-hairpin-helix (HhH)_2_, demonstrated that maximum likelihood (ML) tree inference using IQ-TREE (Trifinopoulos et al., 2016) and empirical Bayesian ancestral sequence reconstruction (ASR) could yield a cenancestral sequence with dramatically increased sequence identity between its duplicated domains (from 21 ± 8% to 46%; Longo et al., 2020). Here we apply similar methods to a recently published dataset of taxonomically diverse prokaryotic SecY sequences (Harris and Goldman, 2021). The published alignment of these sequences is inconsistent with a structural alignment around C.H1 (Supplementary figure 1), but aligning the sequences using MAFFT L-INS-i yields results consistent with the structures, so we use this realignment for our analysis. A few sequences were added or removed to obtain a final full-length (FL) alignment, as described in the Methods.

**Supplementary figure 1.**
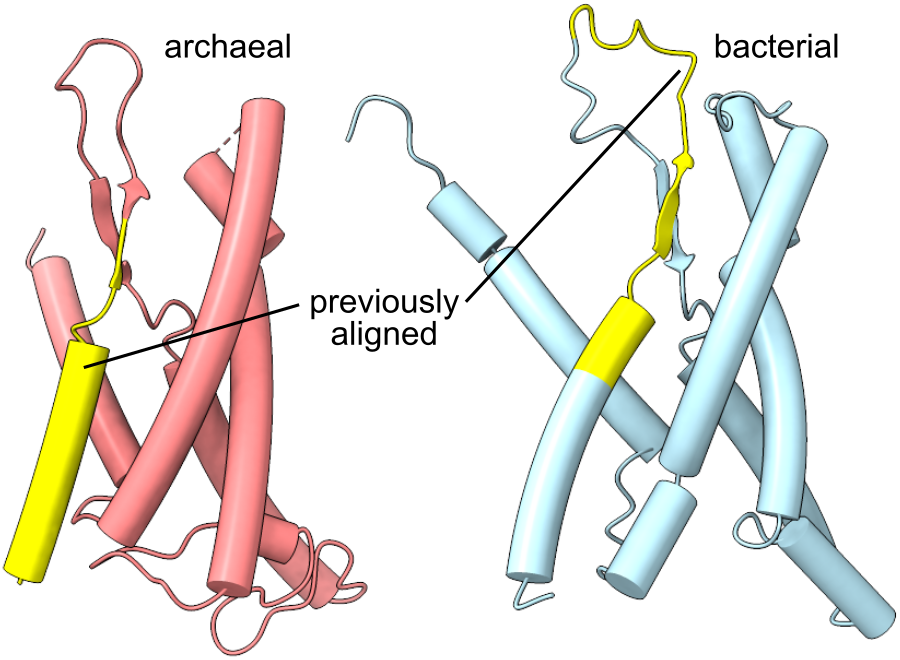
Discrepancy between a structural alignment of archaeal and bacterial C.H1 and the previous sequence alignment of Harris and Goldman, 2021. Archaeal (*M. jannaschii*, 1rh5) and bacterial SecY (*G. thermodenitrificans*, 6itc) C-halves are shown, with a previously sequence-aligned segment highlighted in yellow.

In addition to the FL alignment, we also prepared a structure-guided alignment of the N- and C-half subsequences from the same dataset, as described in the Methods, and used this NC alignment as a separate basis for ASR. Finally, we also used a version of the NC alignment which includes only the five structurally aligned blocks corresponding to H1-5 (NC-blocks).

Matched-pair tests (Naser-Khdour et al., 2019) did not reject the standard assumptions of symmetry and homogeneity (*p >* 5%), although the data in NC-blocks were insufficient for most tests. Automated phylogenetic model selection was performed with ModelFinder (Kalyaanamoorthy et al., 2017) by the Bayesian information criterion (BIC). In each case, the same model would have been selected by the Akaike information criterion (AIC), whereas the corrected AIC (AICc), which more strongly avoids overfitting, would select a less parameter-rich model. Likelihood mapping (Strimmer and von Haeseler, 1997) showed that while the phylogenetic signal in all three alignments is poor, it is strongest in the FL alignment, and not much degraded by reducing NC to NC-blocks. ML tree inference with IQ-TREE 2 (Minh et al., 2020) was performed with 1000 ultrafast bootstrap replicates (Hoang et al., 2018).

**Supplementary table 1.** Summary statistics from sequence alignment and phylogenetic inference. Abbreviations: LG, Le & Gascuel substitution matrix (Le and Gascuel, 2008); F, empirical amino acid frequencies; R, rate categories; G, gamma-distributed rate categories; Q.yeast, QMaker yeast-specific substitution matrix (Minh et al., 2021).

|  | FL | NC | NC-blocks |
| --- | --- | --- | --- |
| % gaps | 51 | 67 | 0 |
| **sites** | 914 | 678 | 102 |
| parsimony-informative | 630 | 423 | 102 |
| singleton | 118 | 104 | 0 |
| constant | 166 | 151 | 0 |
| **sequences** | 342 | 684 | 684 |
| composition $\chi^2$ test $p < 5\%$ | 9 | 10 | 2 |
| **matched pair test $p$** |  |  |  |
| symmetry | 0.23 | 0.56 | 0.62 |
| marginal symmetry | 0.32 | 0.88 | n/a |
| internal symmetry | 0.26 | 0.34 | n/a |
| **model selection** |  |  |  |
| BIC | LG+F+R9 | LG+F+R10 | Q.yeast+F+R9 |
| AIC | LG+F+R9 | LG+F+R10 | Q.yeast+F+R9 |
| AICc | LG+F+R8 | LG+G4 | LG |
| **likelihood mapping** |  |  |  |
| % fully resolved | 66 | 55 | 52 |
| % partially resolved | 22 | 3 | 3 |
| % unresolved | 12 | 42 | 45 |
| total tree length | 313 | 147 | 163 |

The FL tree accurately reflects most of the shallow relationships within phyla, as in Harris & Goldman 2021. But the deepest branch, between archaea and bacteria, is extraordinarily long, 5.9 expected substitutions per site (sps), which means the overwhelming majority of sites should be expected to differ in both archaea and bacteria from whatever their identity was in the cenancestor (Supplementary figure 2a). This length may be regarded as a lower bound on the true branch length, since excluding the fastest-evolving sites or using more parameter-rich models typically yields even longer archaeal-bacterial branches (Moody et al., 2021). It is thus unsurprising that ancestral sequences reconstructed at the root nodes of archaea or bacteria, using IQ-TREE 2’s implementation of the empirical Bayesian method, displayed halves with poor sequence similarity (15-20% identity across H1-5), not significantly different from that between the halves of extant sequences (16 ± 3% across H1-5).

The NC and NC-blocks alignments yielded trees with a long central branch separating the N-halves and C-halves (5.8, 6.8 sps; Supplementary figure 2b,c). The subtree for each of the two halves displayed some topological discrepancies with the other half’s subtree. These discrepancies were particularly acute in the NC-blocks tree, which puts the root of the N-half inside archaea, but the root of the C-half inside bacteria. Such extreme discrepancies are common in trees of universal paralogs, which are generally so divergent that outgroup rooting is inadvisable due to severe model violations (Gouy et al., 2015).

Unlike the NC-blocks tree, the NC tree places the root of both subtrees between archaea and bacteria. This difference is difficult to attribute to any meaningful signal in the data, since the NC-blocks alignment retains the high-confidence columns from structural alignment and nearly as much phylogenetic signal as the NC alignment (Supplementary table 1). It may instead be an artefact of long branch attraction (LBA), induced by the greater number of high variability and poorly aligned sites contained in the NC alignment. The indistinguishability of this tree topology from what LBA would induce makes the accuracy of universal paralog trees suspect in general (Gouy et al., 2015). Thus although one could estimate ancestral N- and C-half and proto-SecY sequences using these trees, one has little reason to think those estimates would be accurate.

**Supplementary figure 2.**
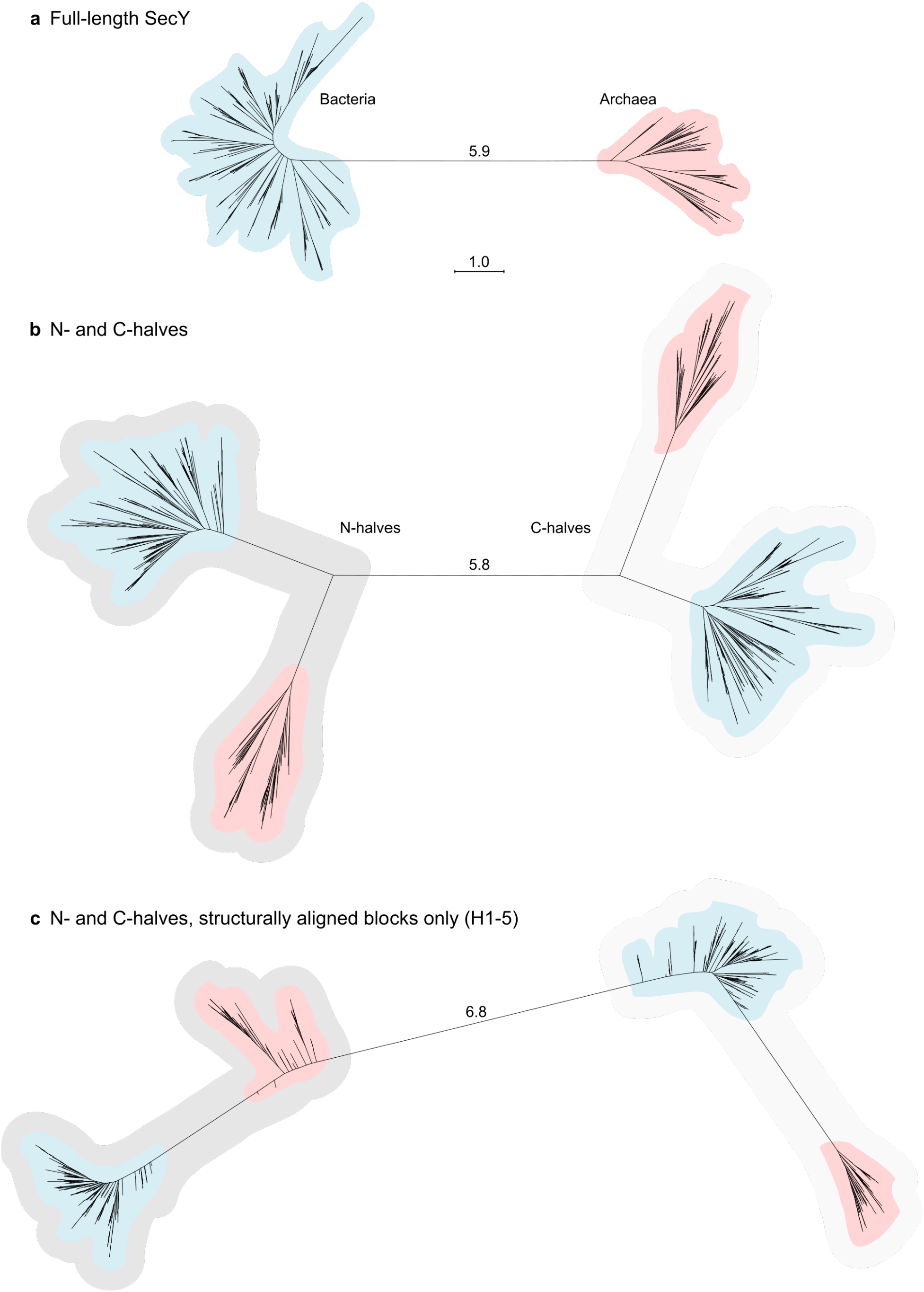
Maximum likelihood trees inferred for SecY and its halves. All trees are shown on the same scale. The length unit is the expected number of substitutions per site. **a** Tree inferred from the FL alignment. **b** Tree inferred from the NC alignment. **c** Tree inferred from the NC-blocks alignment.

The methods applied here may have proven less effective than when previously applied to HhH domains (Longo et al., 2020) because those domains are less divergent than the SecY halves. The HhH domains each conserve independent nucleotide-binding activity, and thus are just as conserved as are the (HhH)_2_ proteins as a whole (22.4 ± 7.8% between domains *vs* 19.5 ± 4.4% between UvrC and ComEA). By contrast, the SecY domains form an obligate complex around a single active site, and are much less conserved than SecY as a whole (12.5 ± 2.2% between domains *vs* 20.3 ± 1.6% between archaea and bacteria). It is also known that general-purpose substitution models like LG, used here, are a poor fit to substitutions occurring in the heterogeneous membrane environment (Jones et al., 1994). Regardless of the true cause, the sequences of SecY’s ancestors remain recondite. The main text therefore focuses on more stable characteristics, namely structure, mechanism, function, and a few functionally important, highly conserved residues.

## Supplementary figures

**Figure 2-Figure supplement 1.**
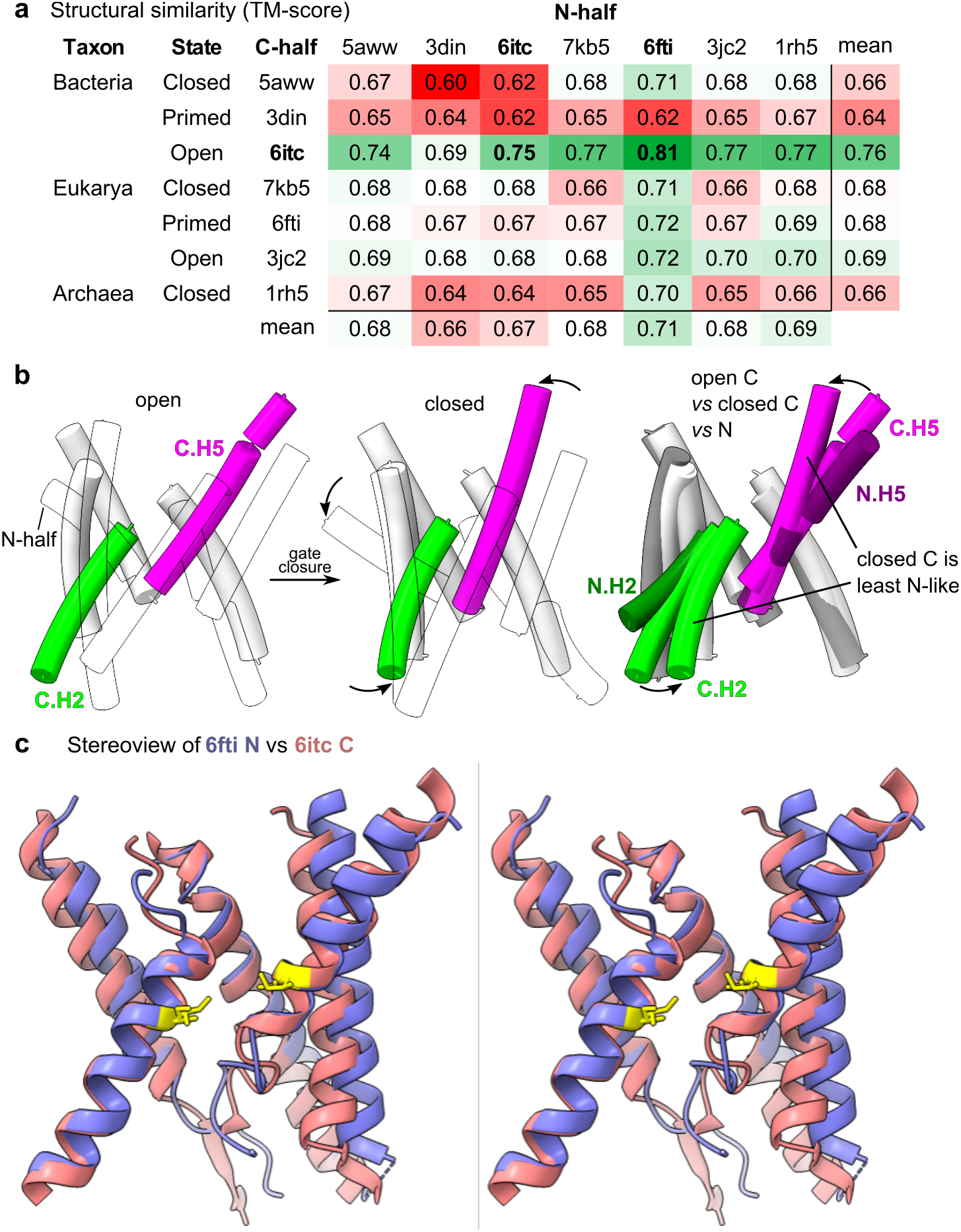
Structural similarity and symmetry breaking between SecY halves. **a** Pairwise comparisons of the SecY halves by TM-score. TM-scores can range from 0 to 1. Boldface scores indicate the most similar pair from the same structure (6itc) or from any structure (6itc C, 6fti N). **b** Symmetry-breaking tilts in C.H2 and C.H5 induced by channel closure. At left, the TMHs from open (5aww) and closed (6itc) models are shown, with the N-half rendered in transparent outline. At right is a superposition of closed (5aww) and open C-halves (6itc) and an N-half (6itc). All models were aligned by H1/4. **c** Walleye stereoview of the most similar pair of N- and C-halves. The pore ring residues are shown as sticks and highlighted yellow.

**Figure 3-Figure supplement 1.**
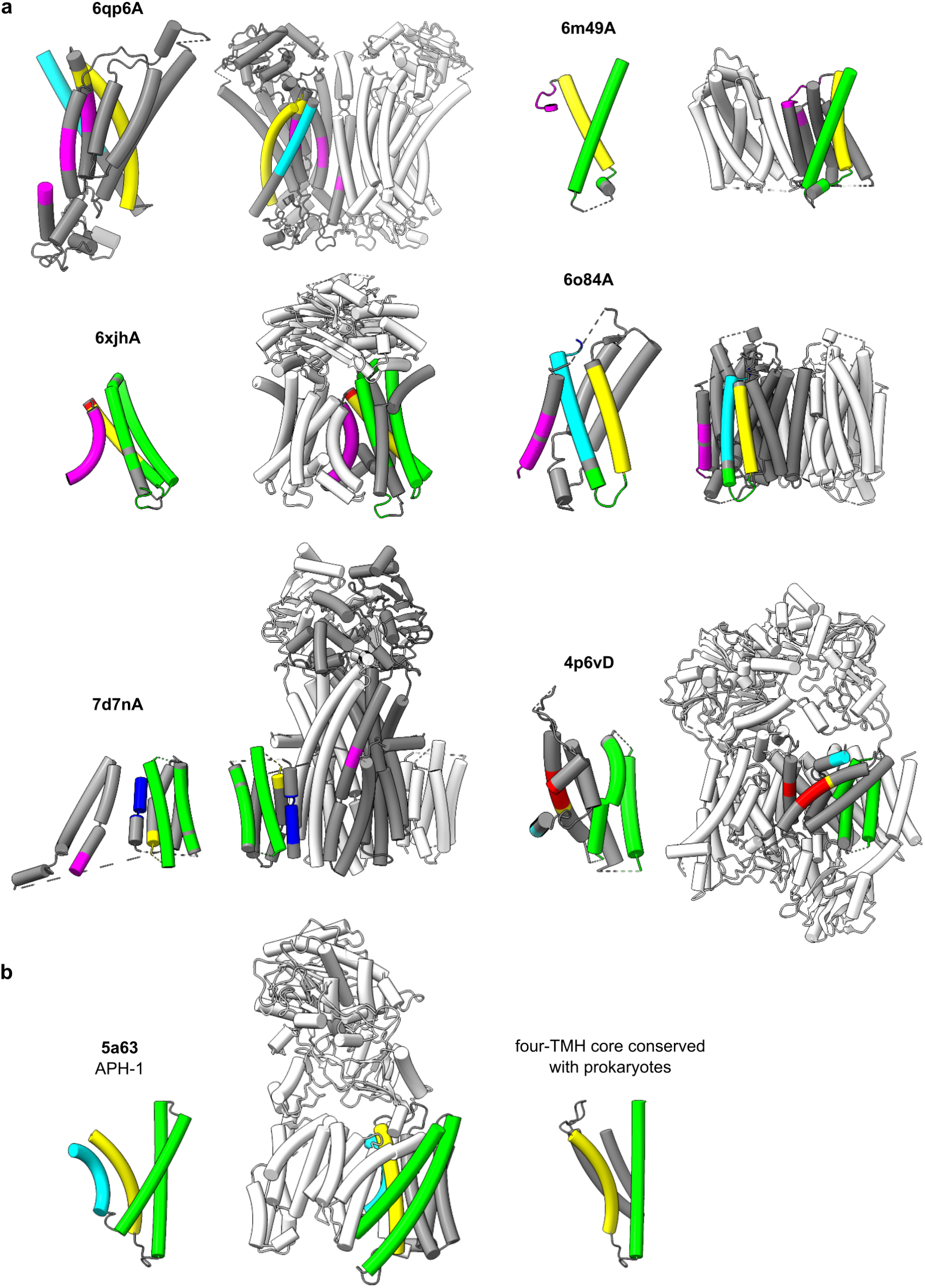
Structures of non-Oxa1 superfamily top Dali hits. **a** Isolated hits. The aligned chain is shown in grey, and any other chains in the model are shown in white. Regions of each hit are colour-coded according to which part of the query structure they aligned with, using the colour scheme from Figure 4. **b** The multiple hit APH-1. In addition to the representations shown in panel a, at right are shown the four TMHs conserved among APH-1’s prokaryotic homologs. As in a, the non-aligned TMHs are shown in grey, and the two aligned TMHs are coloured.

**Figure 4-Figure supplement 1.**
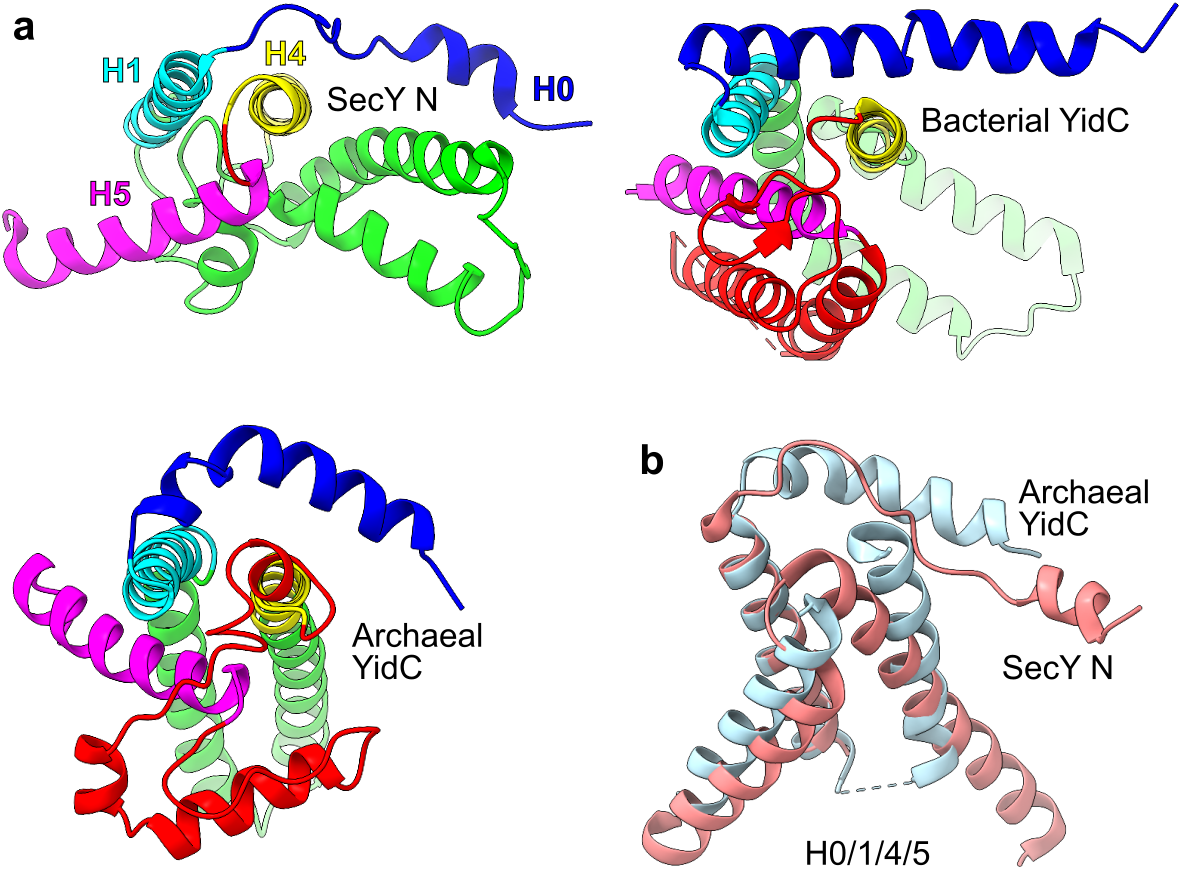
Structural comparison of H0 in SecY and the Oxa1 superfamily. **a** Matched views of the SecY N-half (*M. jannaschii*, 1rhz), bacterial YidC (*B. halodurans* YidC2, 3wo6), and archaeal YidC (*M. jannaschii* MJ0480, see Figure 4-Figure supplement 4). **b** Alignment of YidC and SecY N, both from *M. jannaschii*.

**Figure 4-Figure supplement 2.**
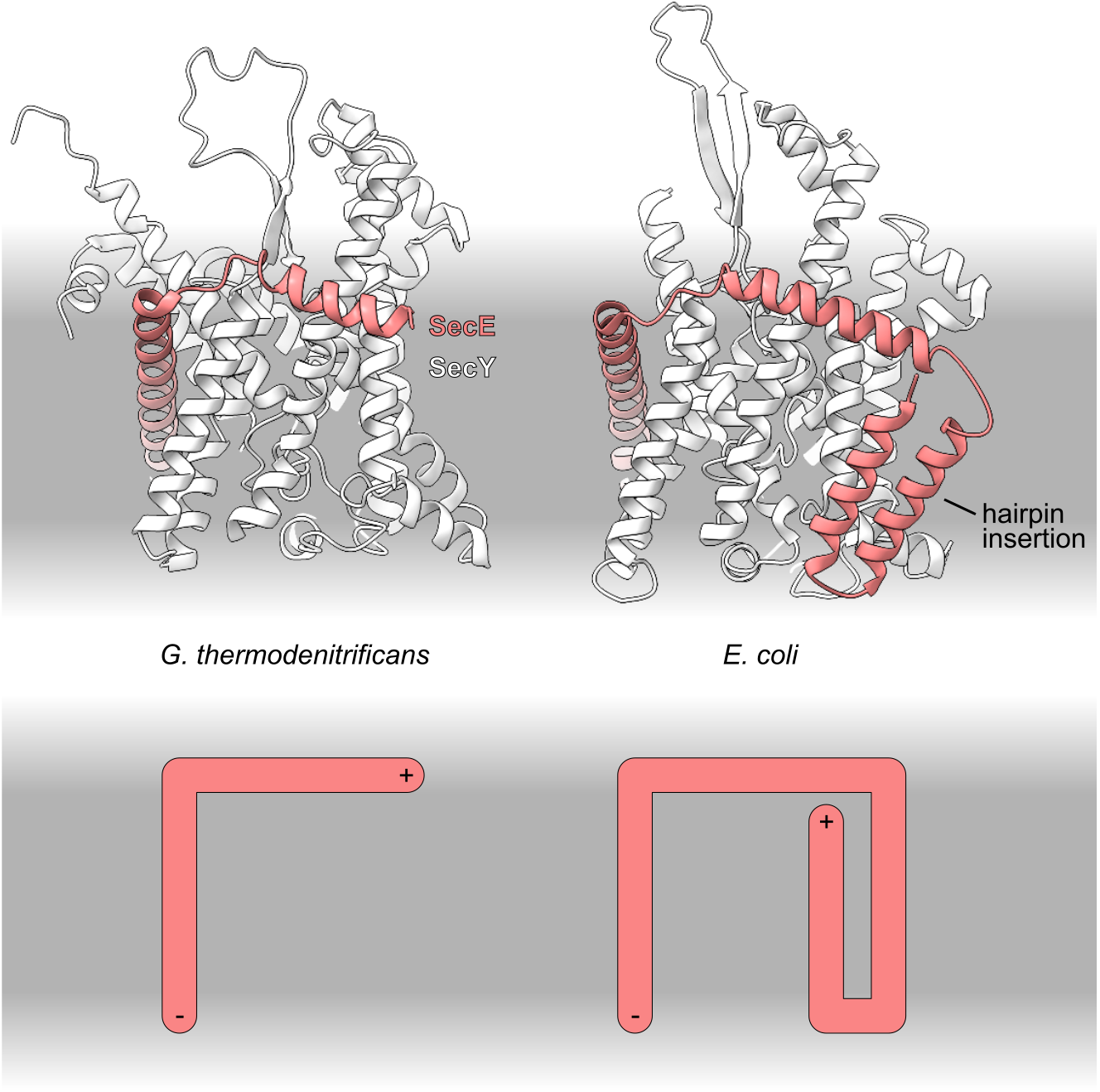
Structure of the acquired transmembrane hairpin in SecE. Left: *G. thermodenitrificans* SecYE (6itc). Right: *E. coli* SecYE (6r7l; Kater et al., 2019). Top: molecular models of SecY (white) and SecE (light coral). Bottom: diagram of SecE topology, in which plus and minus signs indicate the N- and C-termini.

**Figure 4-Figure supplement 3.**
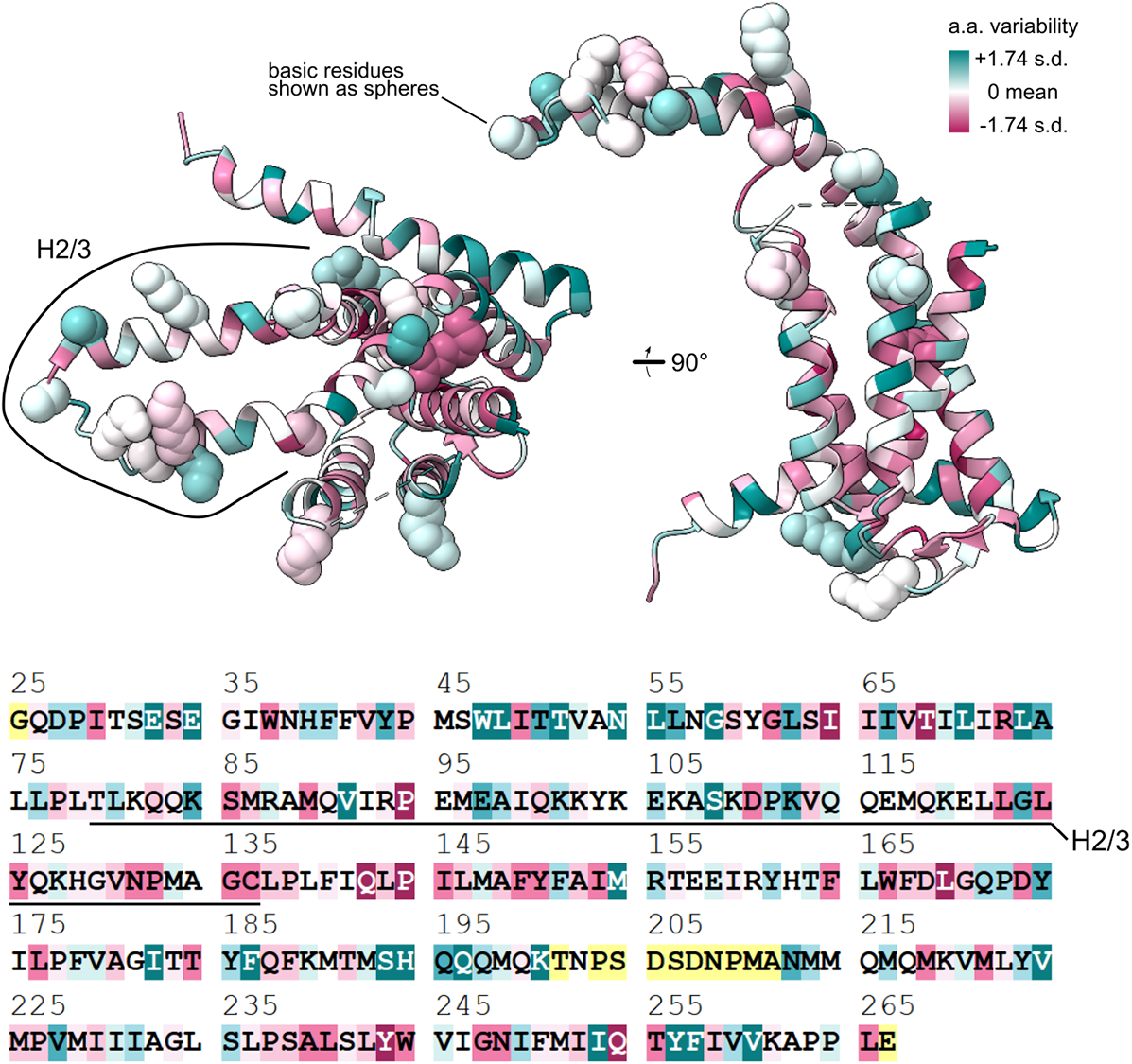
Amino acid conservation in bacterial YidC. The conservation scores and annotated sequence and PDB model shown (*B. halodurans* YidC2, 3wo7A) were retrieved from ConSurf-DB (Chorin et al., 2020). The structure is colour-coded using the continuous scale shown, whereas the sequence is colour-coded on a similar but quantized scale. The scale shown encompasses the minimum but not maximum score.

**Figure 4-Figure supplement 4.**
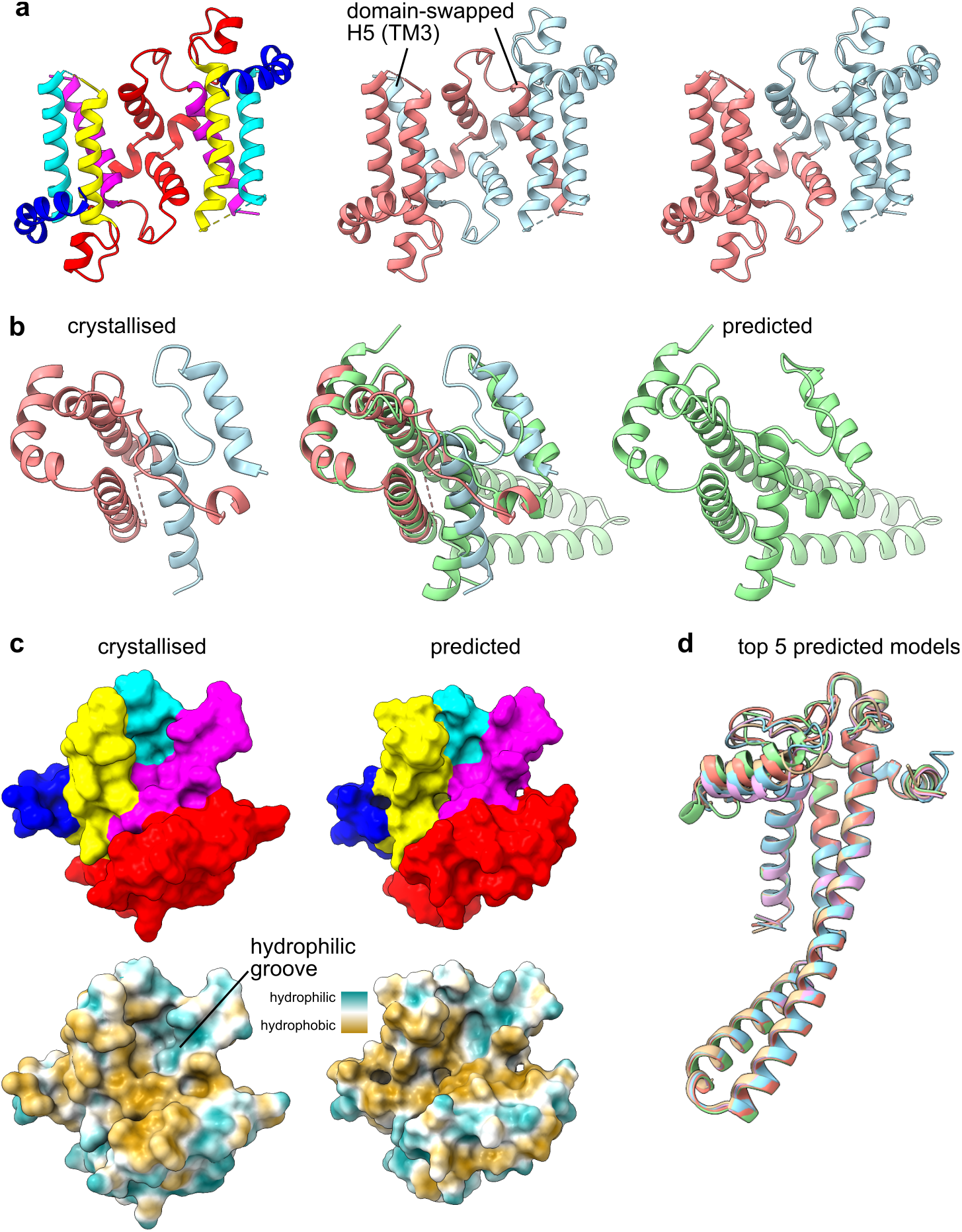
Crystallised and predicted structures of archaeal YidC. **a** Domain swapping in crystallised archaeal YidC (*M. jannaschii* MJ0480 a.k.a. Ylp1, 5c8j). Left: colour-coded by consensus element as in Figure 4. Centre: colour-coded by chain. Right: colour-coded by chain, except the domain-swapped parts of H4/5 have been recoloured to match the chain against which it packs. **b** Alignment of the crystal structure (left, red and blue chains) with the trRosetta-predicted structure (right, green). **c** Comparison of the solvent-excluded surfaces of the crystallised and predicted structures, colour-coded by consensus element (top) or by hydropathy (bottom). **d** Alignment of the 5 highest-probability models built by trRosetta. The models were aligned by their first 40 amino acids.

**Figure 8-Figure supplement 1.**
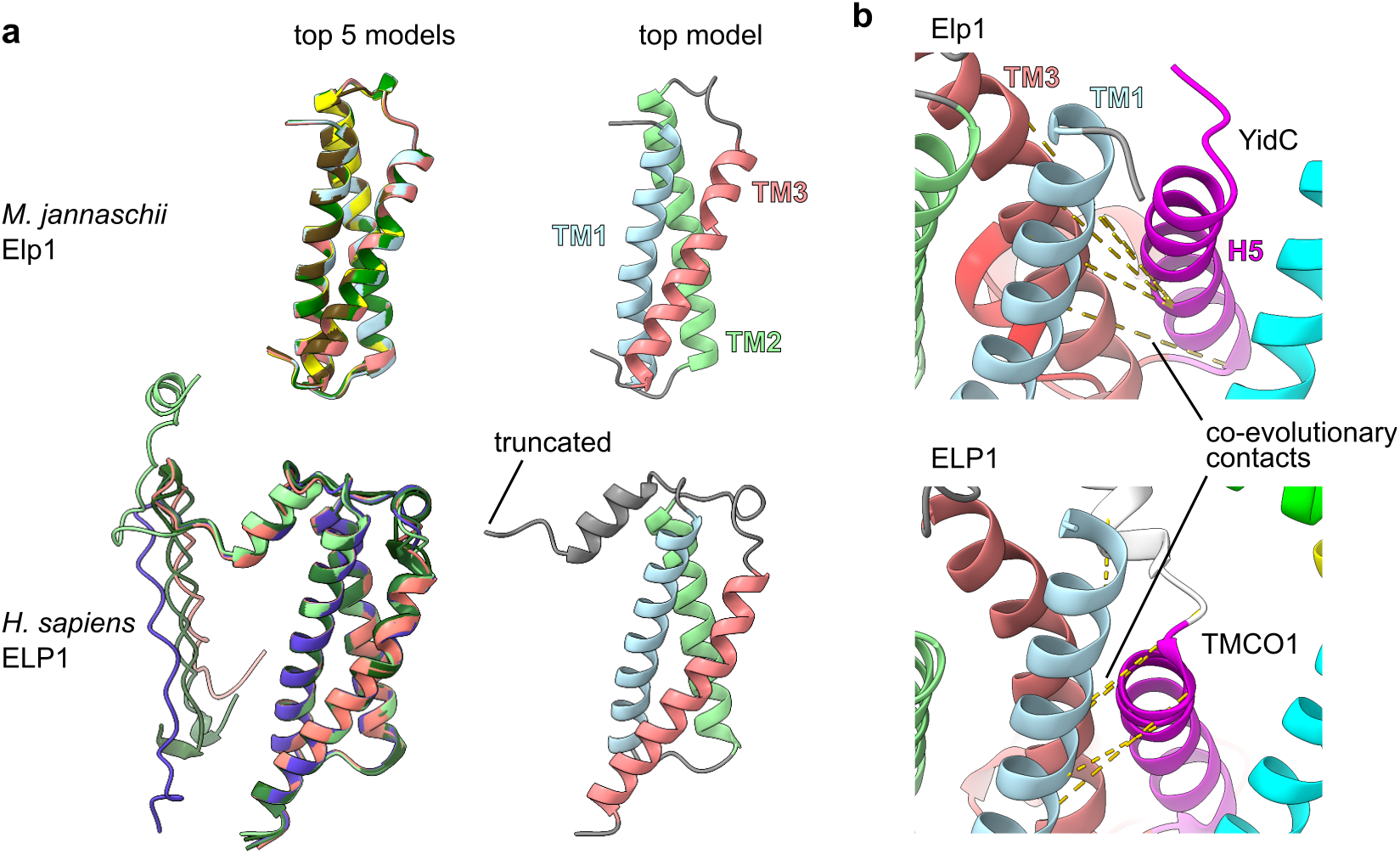
Structure and contact prediction for archaeal and human EMC6-like proteins. **a** Alignment of the five highest-probability models determined by trRosetta (left) and an annotated view of the single best model (right), with disordered residues omitted. Models were predicted from the protein sequence of *M. jannaschii* Elp1 (MJ0606) and *H. sapiens* ELP1 (C20orf24). **b** Close-up view of the five highest-probability contacts between YidC/Elp1 and TMCO1/ELP1, represented by dashed gold lines.

**Figure 8-Figure supplement 2.**
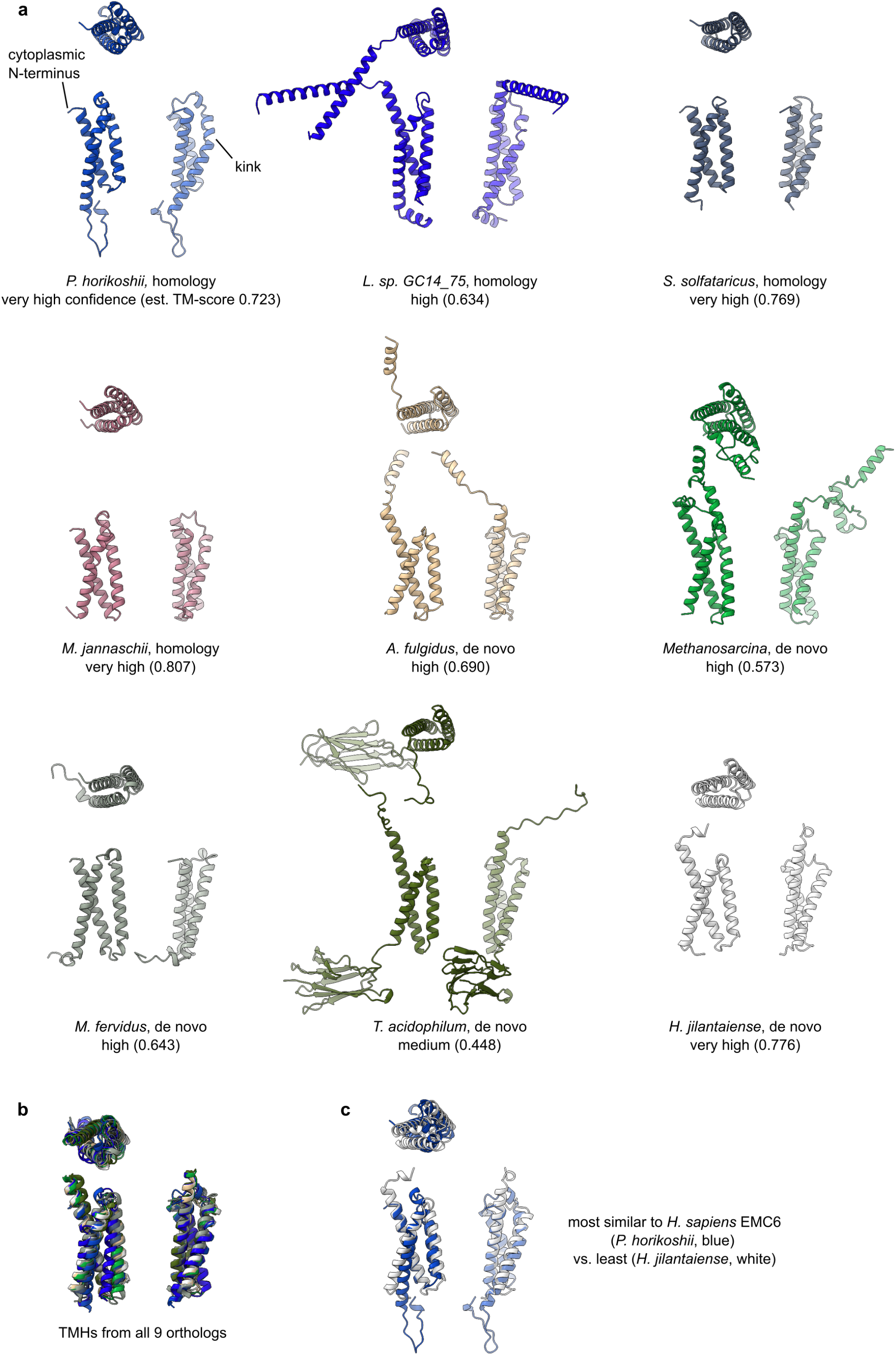
Structural models for nine diverse archaeal EMC6-like proteins. **a** Front, top, and side views of each model, separated by 90° rotations. A kink in TM3 which is present in the homology models but absent from the *de novo* models is indicated. **b,c** Alignments of the indicated structural models.

**Figure 11-Figure supplement 1.**
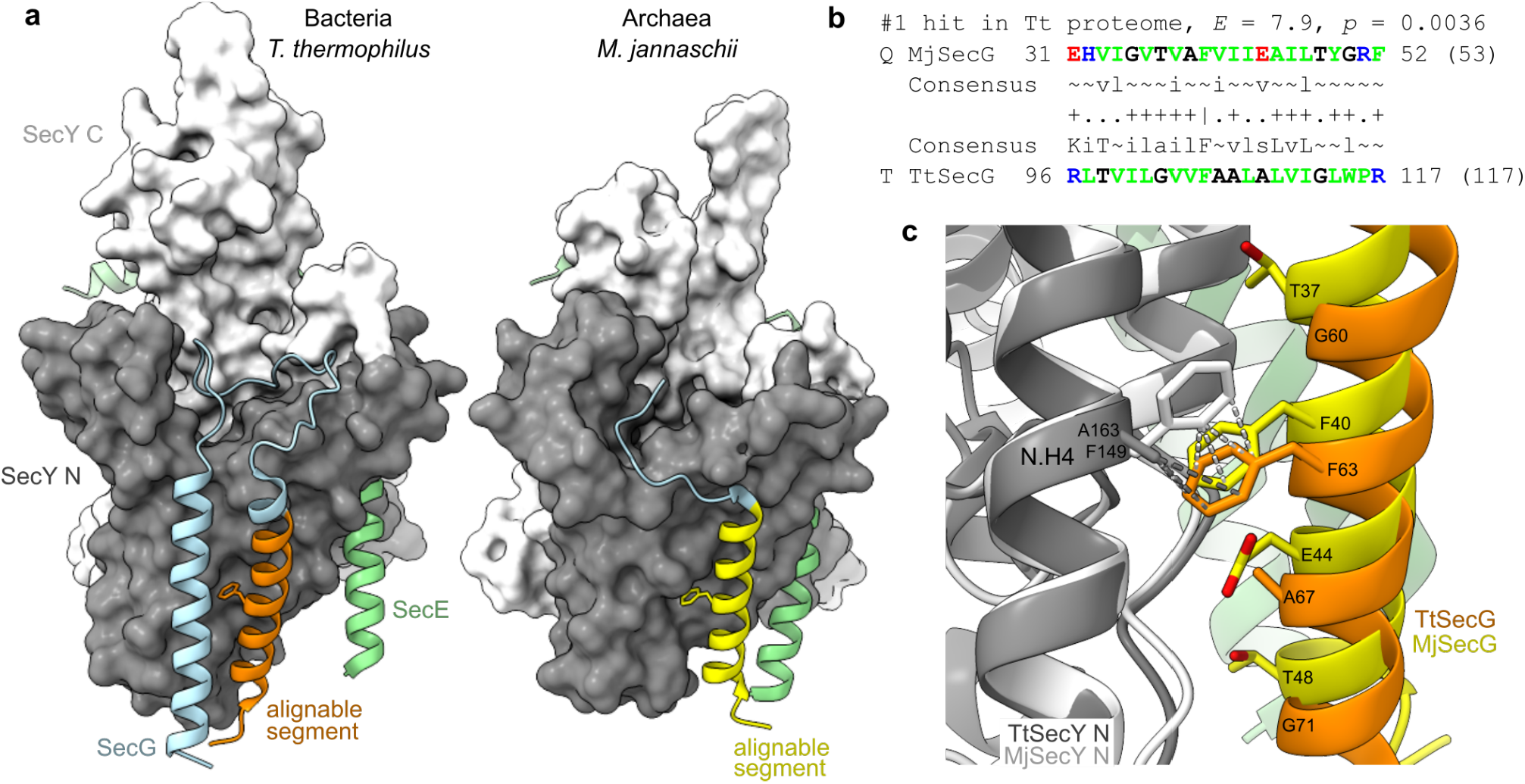
Similarity between archaeal and bacterial SecG. **a** *T. thermophilus* and *M. jannaschii* SecYEG, shown separately but oriented by aligning the two SecY TMs against which SecG packs (N.H1 and N.H4). The SecG subsequences identified as homologous by HHpred (panel **b**) are highlighted in orange and yellow. **b** The top hit from an HHpred query of the *T. thermophilus* proteome with *M. jannaschii* SecG. **c** The hydrophilic face of SecG and a conserved phenylalanine which contacts a specific residue in SecY N.H4.

