## Supplementary material for "A unified evolutionary origin for the ubiquitous protein transporters SecY and YidC": Figure 8-Source data 1. Structure and contact prediction for archaeal and human EMC6-like proteins: 0README_total

#----------------------------------------------------------------------
# Explanation of files in {seqid}.all_in_one.zip,
# where seqid is a number assigned by IntConPred to the query sequences
#----------------------------------------------------------------------

0README_total         : This README file.

#--- input sequences ---#
{seqid}A.fasta        : The first user input sequence in FASTA format.

{seqid}B.fasta        : The second user input sequence in FASTA format.


#--- single Multiple Sequence Alignment ---#
{seqid}A.a3m          : Generated Multiple Sequence Alignment (MSA) in A3M format for the first input sequence.

{seqid}B.a3m          : Generated Multiple Sequence Alignment (MSA) in A3M format for the second input sequence.


#--- merged Multiple Sequence Alignment ---#
{seqid}A_{seqid}B.a3m_genomic     : Merged MSA in A3M format using genomic approach.

{seqid}A_{seqid}B.a3m_phylogenic  : Merged MSA in A3M format using phylogenic approach.


#--- predicted interfacial contact ---#
{seqid}A_{seqid}B.gcnn_genomic    : Predicted interfacial contact using genomic approach.

{seqid}A_{seqid}B.gcnn_phylogenic : Predicted interfacial contact using phylogenic approach.


#--- merged results ---#
{seqid}A_{seqid}B.gcnn_intra_A    : Intra-molecular predicted contact for the first sequence.

{seqid}A_{seqid}B.gcnn_intra_B    : Intra-molecular predicted contact for the second sequence.

{seqid}A_{seqid}B.gcnn_inter      : Inter-molecular predicted contact.

{seqid}A_{seqid}B.gcnn_inter_list : Sorted list of the inter-molecular predicted contact.

{seqid}A_{seqid}B.gcnn_inter.png  : Heatmap of the inter-molecular predicted contact.


[note]:
In this package, except the image file, all the other files are saved in plain text format.
