## Supplementary figures and images for "A unified evolutionary origin for the ubiquitous protein transporters SecY and YidC"

### 600283A_600283B.gcnn_inter.png

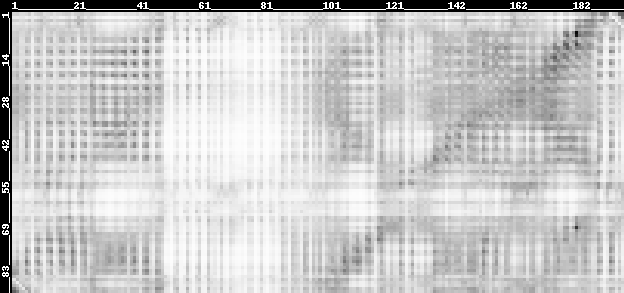

### 600407A_600407B.gcnn_inter.png

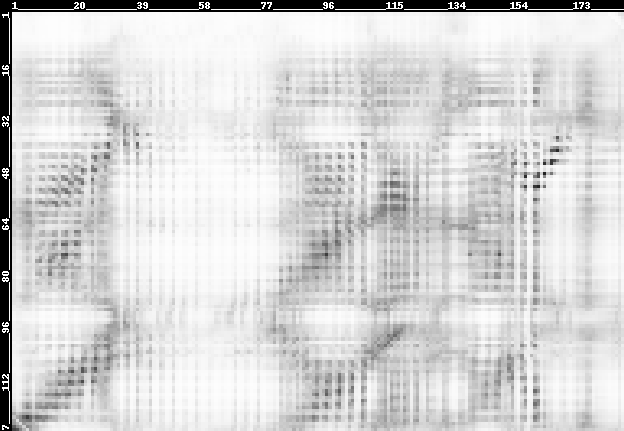

### seq.cont.png

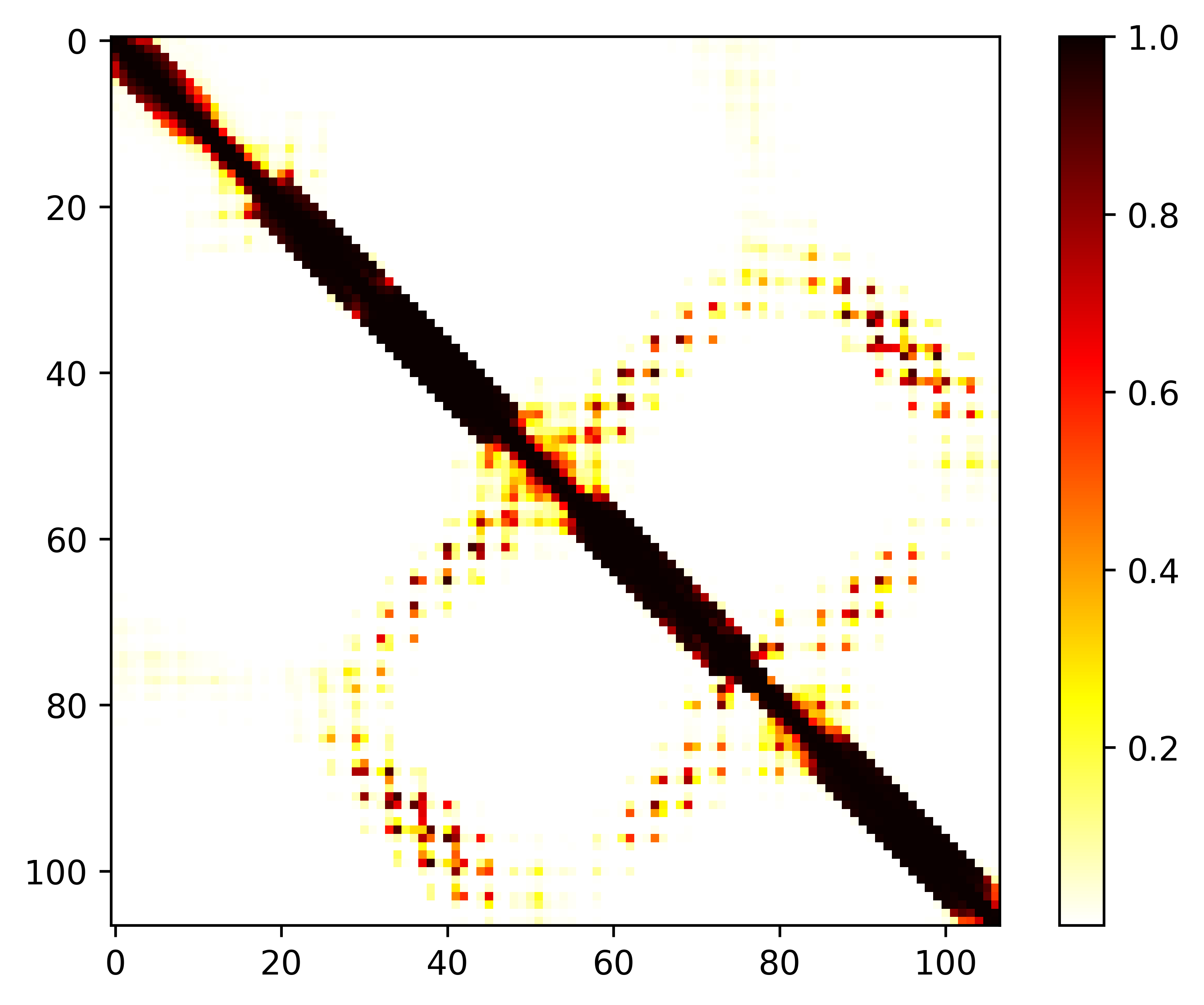

### seq.cont.png

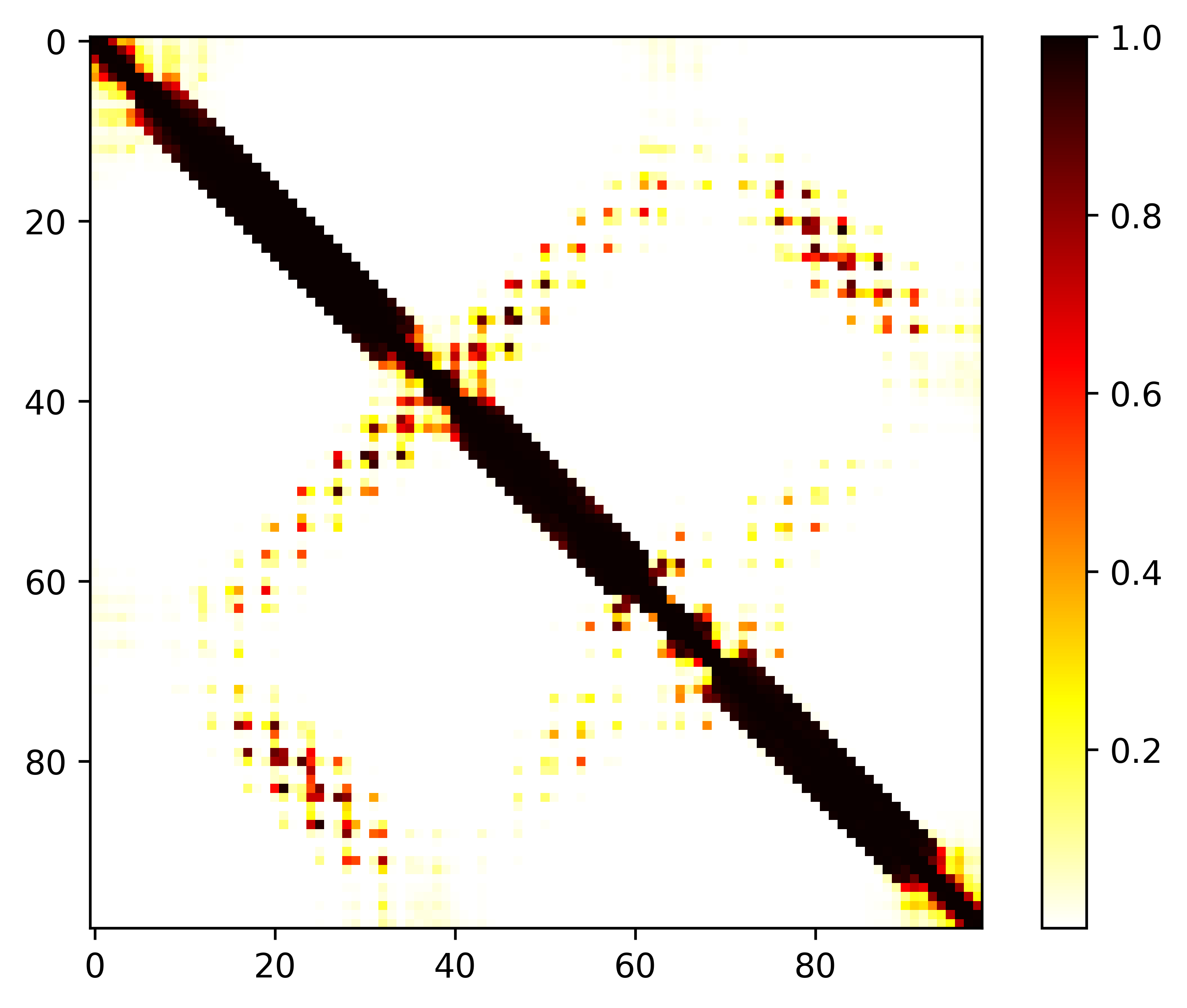

### seq.cont.png

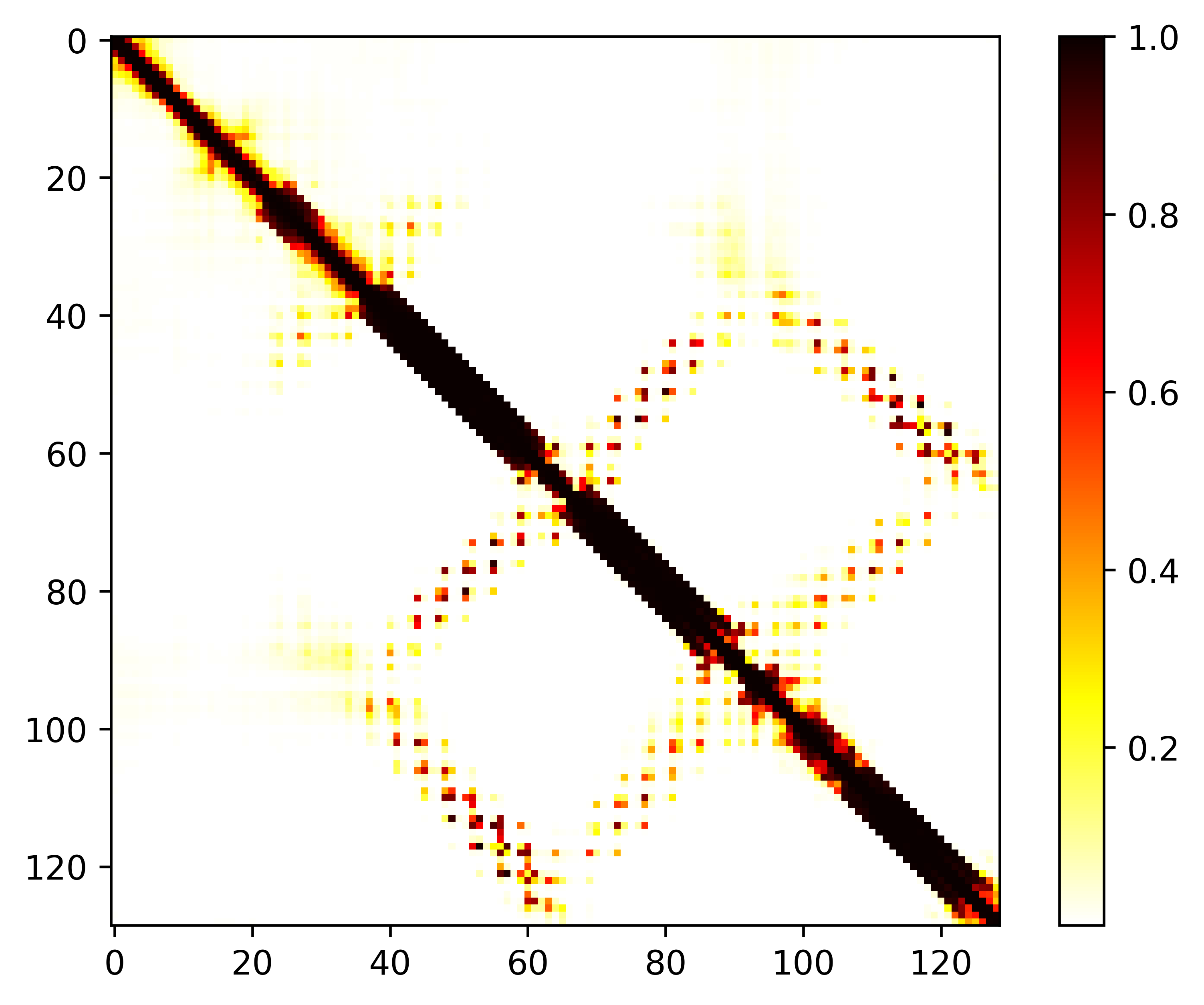

### seq.cont.png

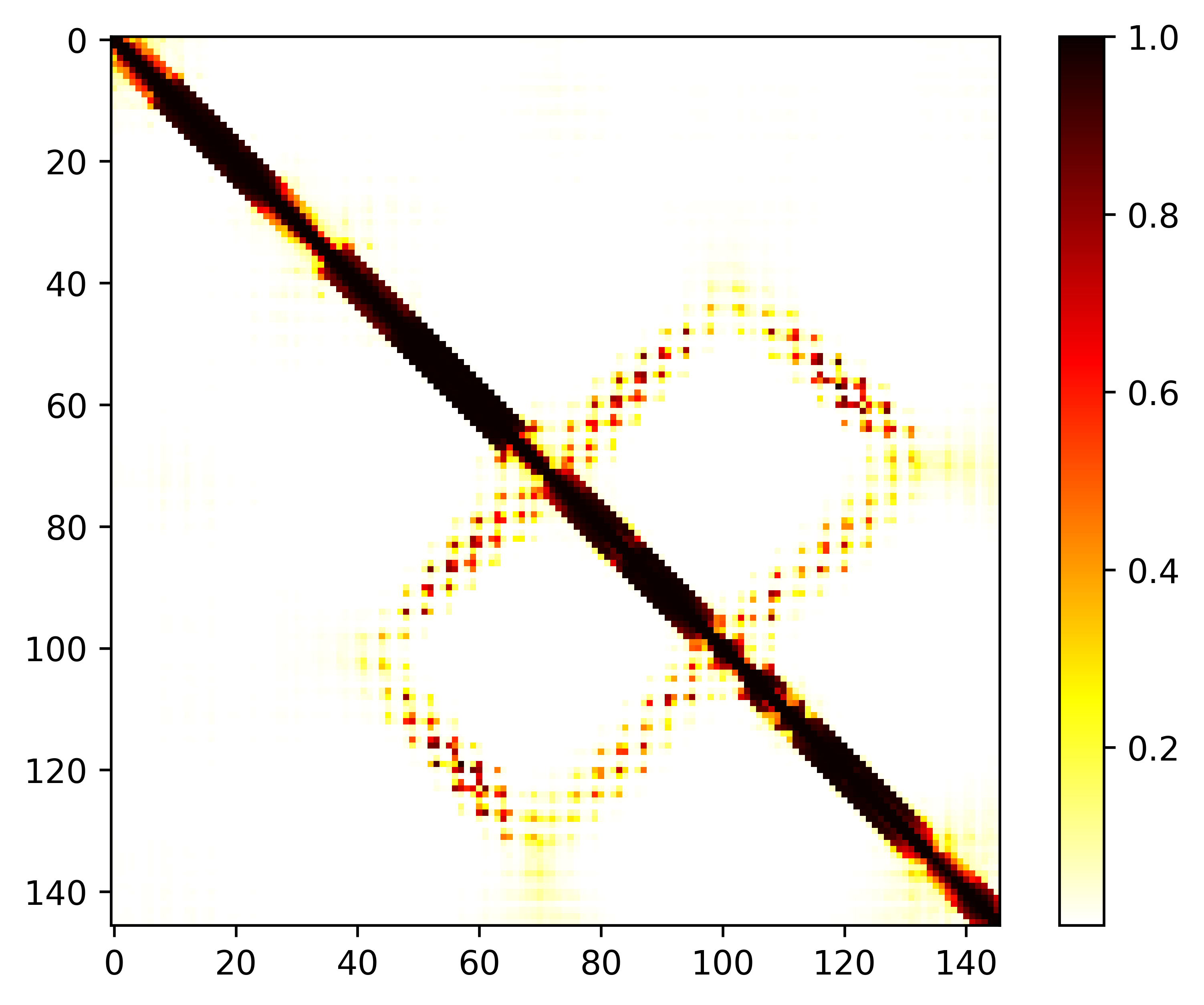

### seq.cont.png

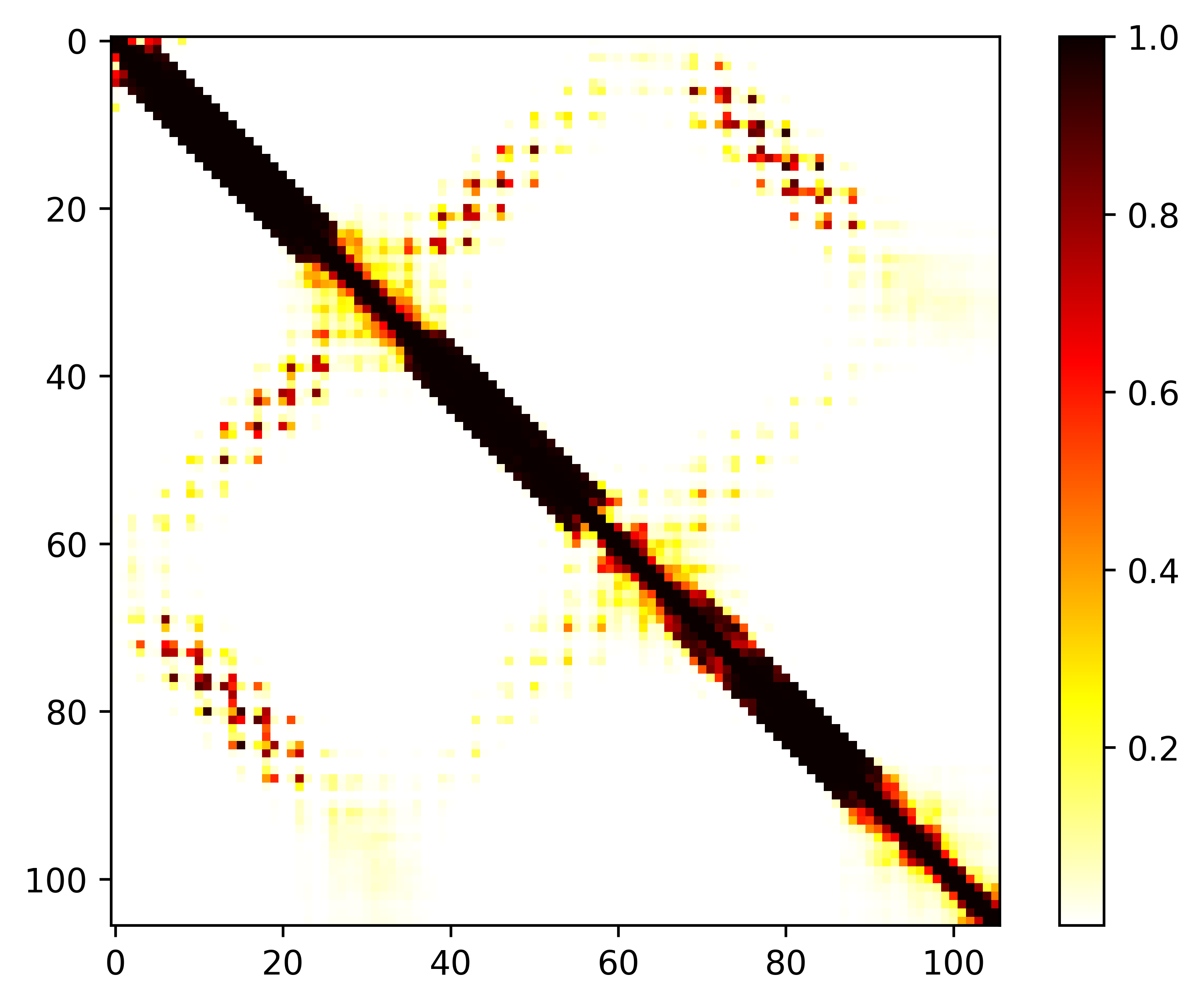

### seq.cont.png

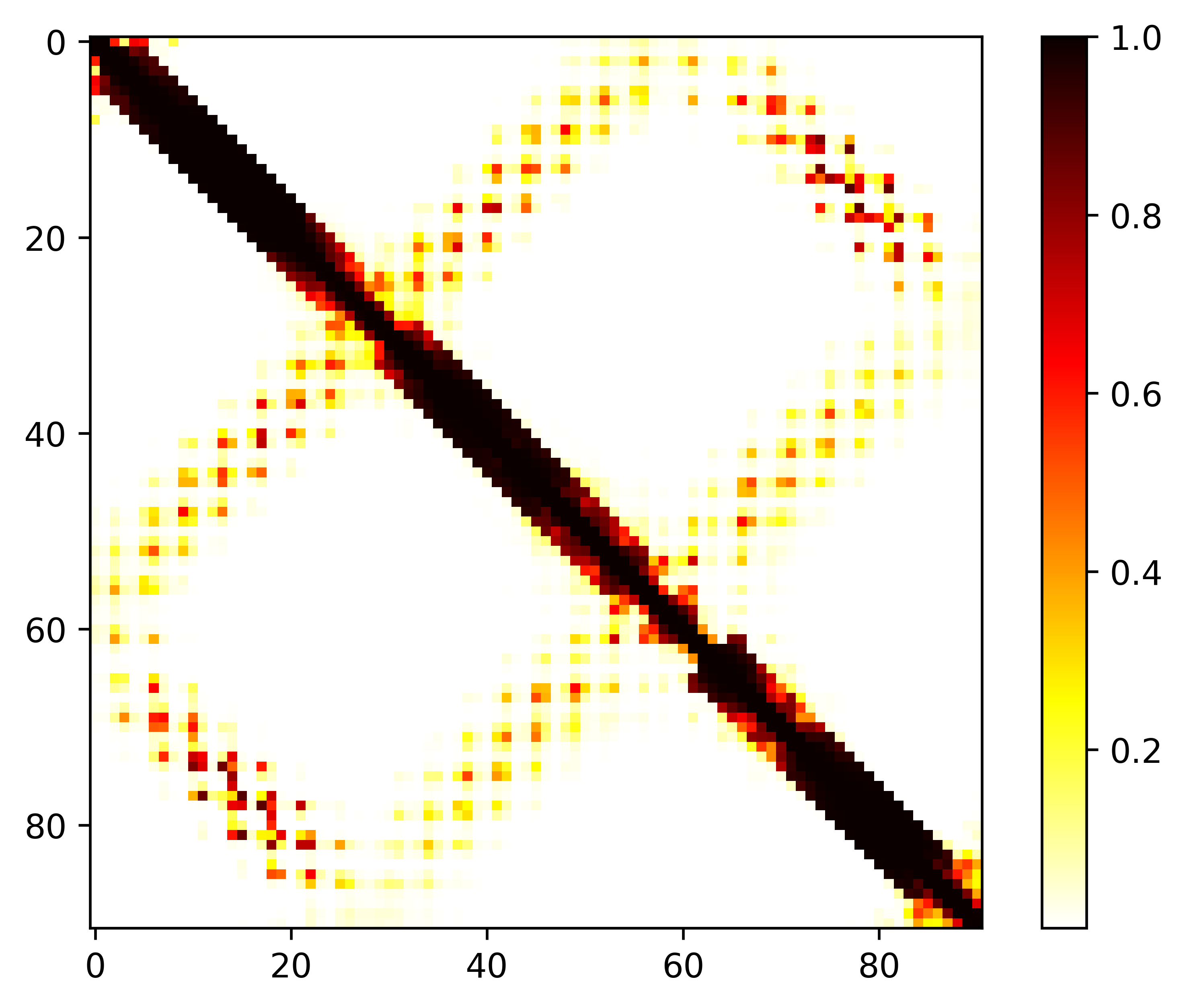

### seq.cont.png

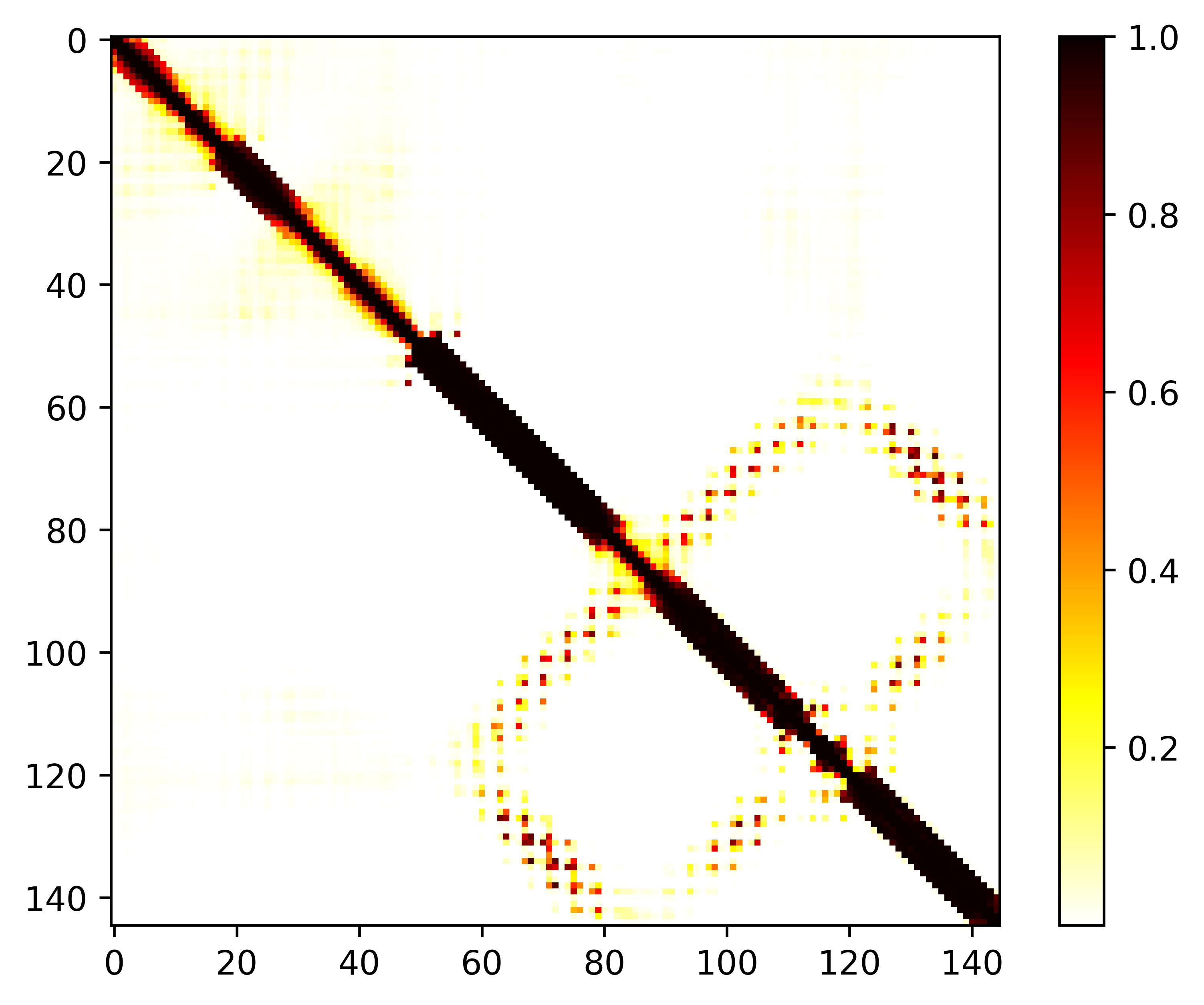

### seq.cont.png

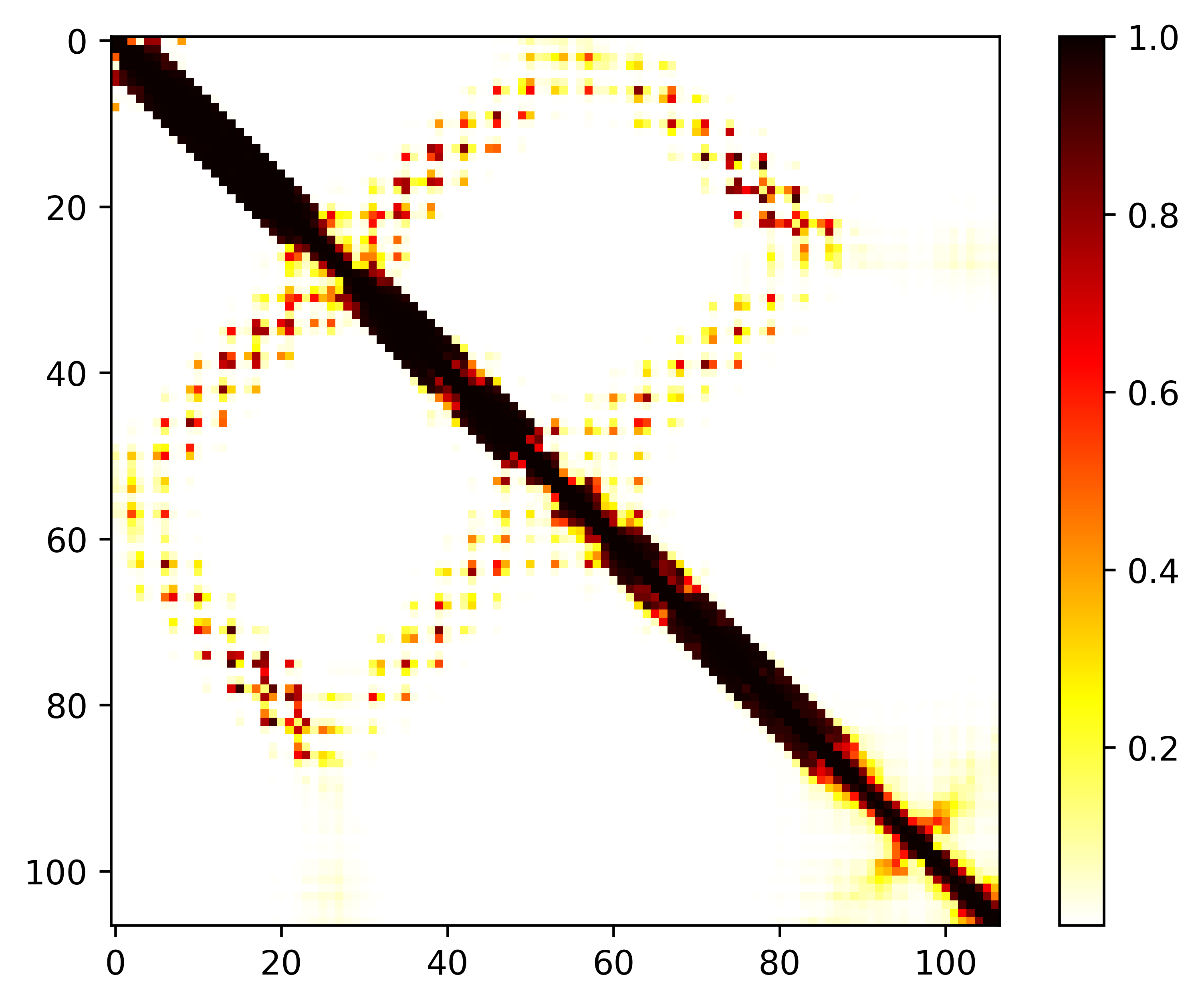

### seq.cont.png

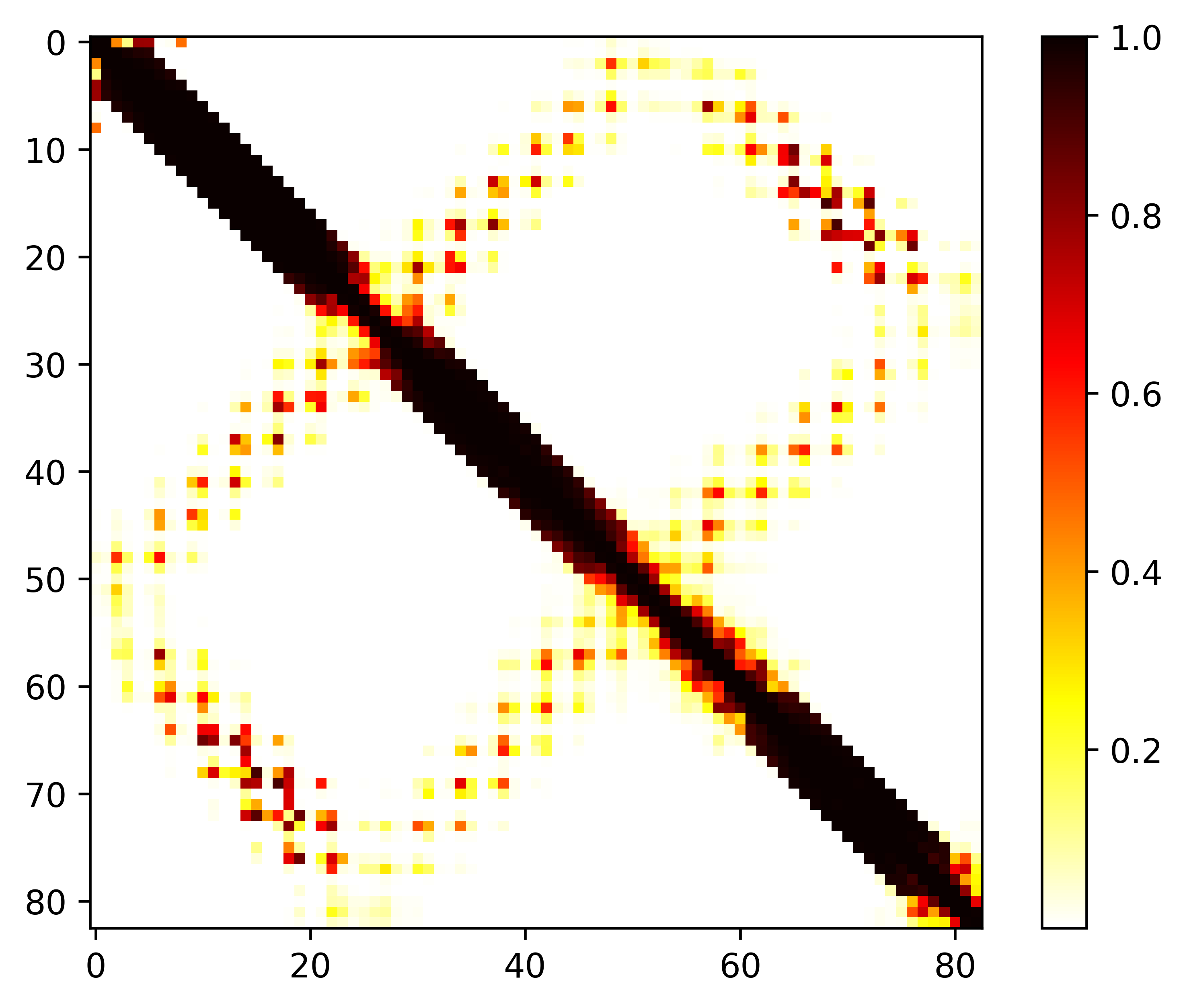

### seq.cont.png

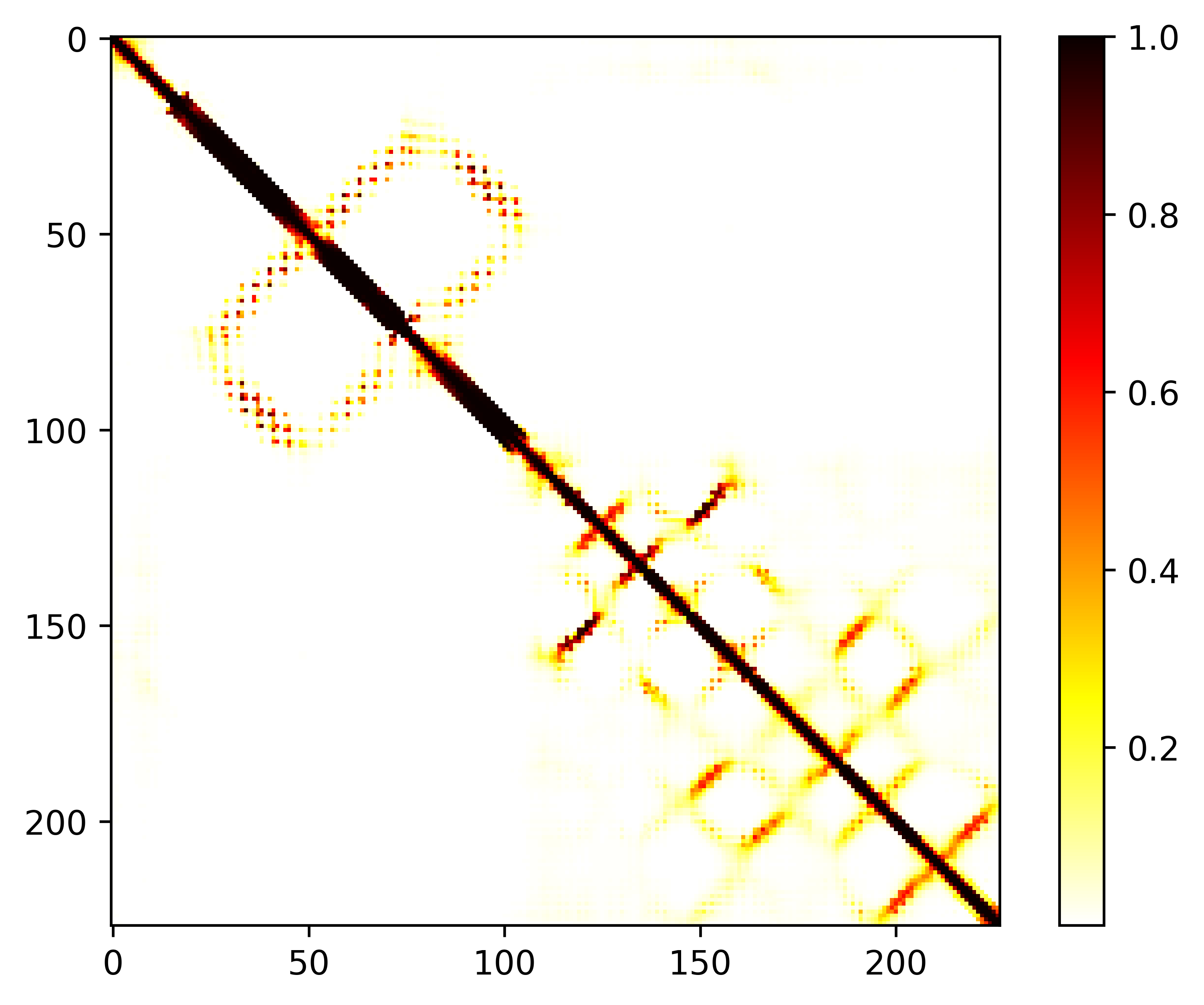

### seq.dist.png

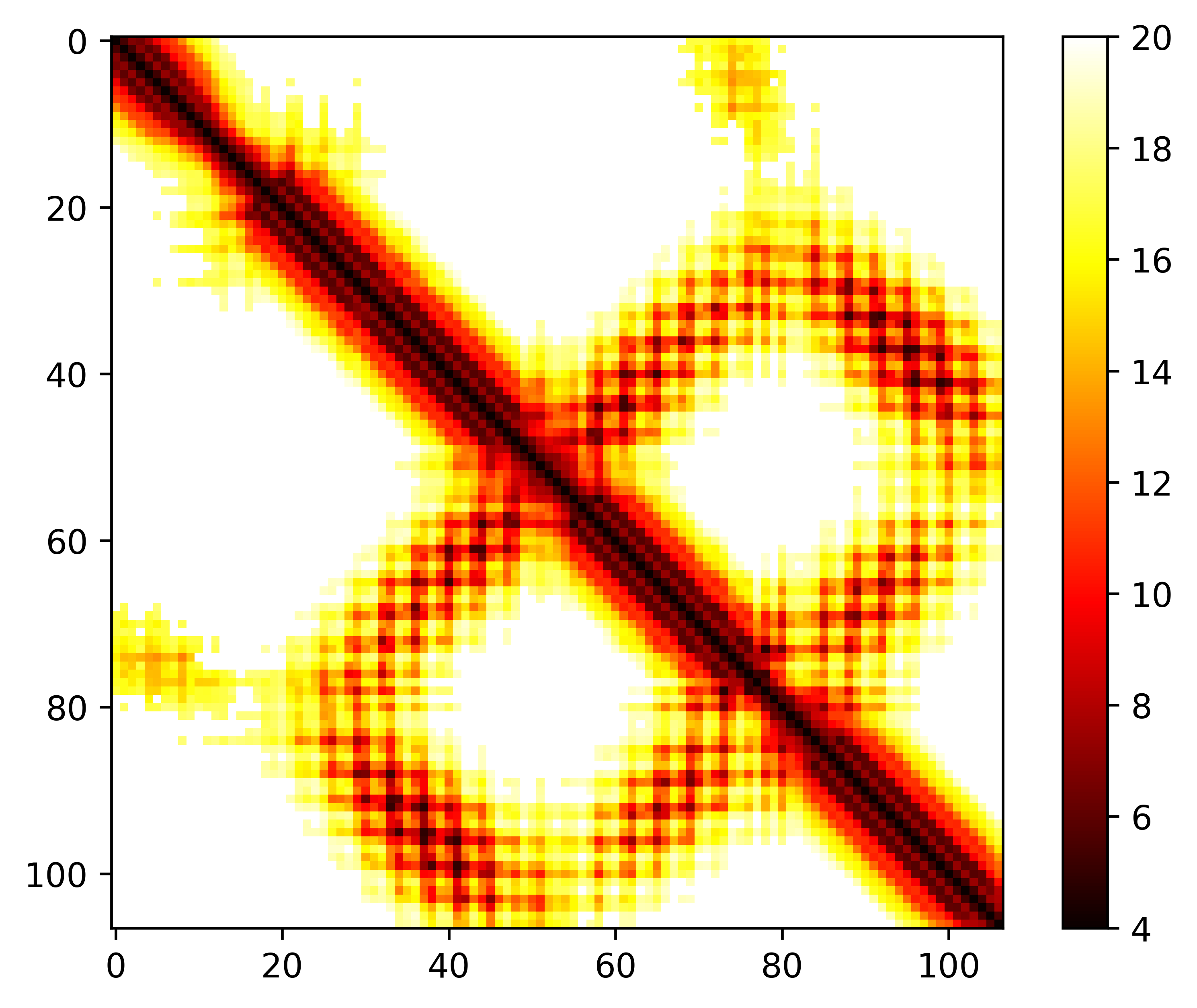

### seq.dist.png

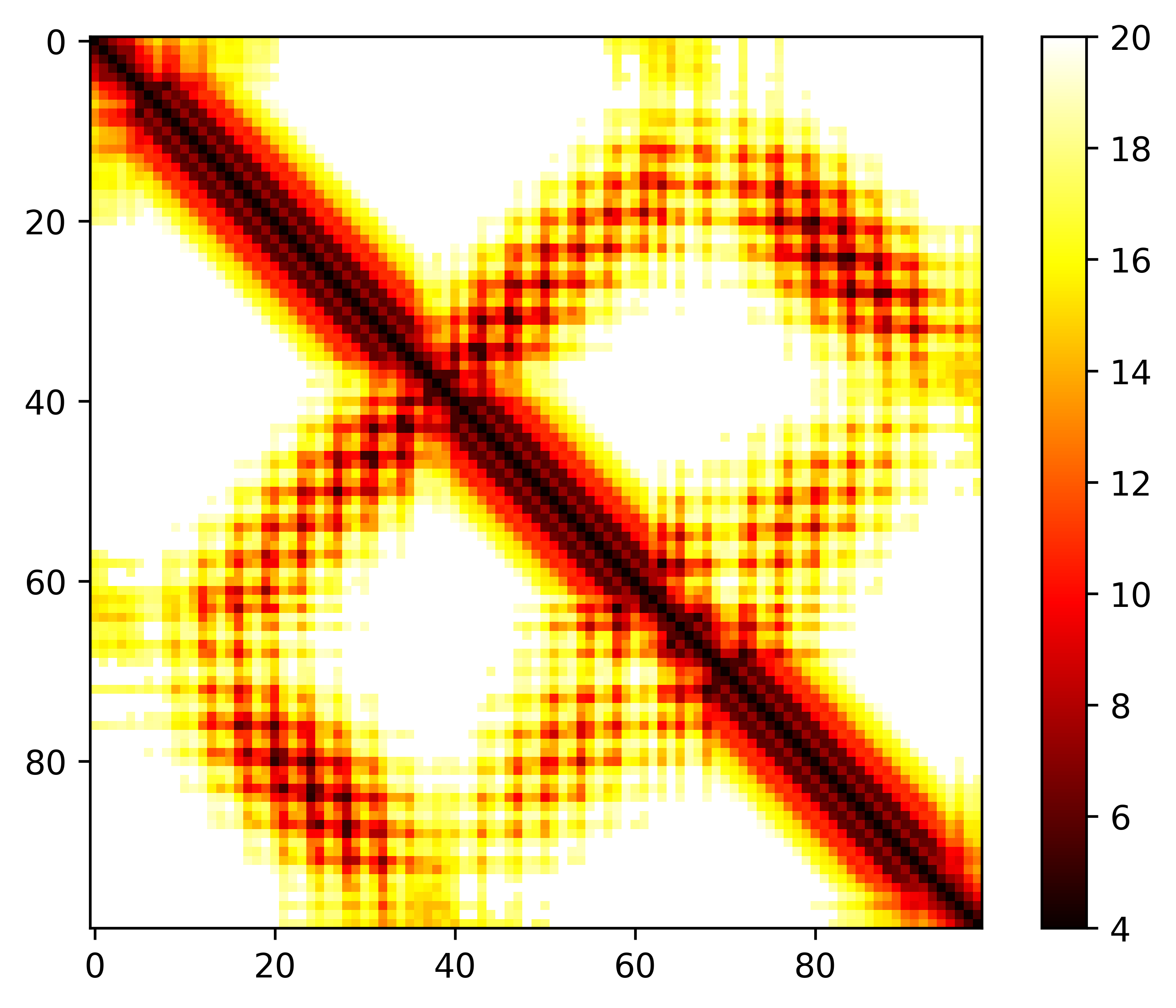

### seq.dist.png

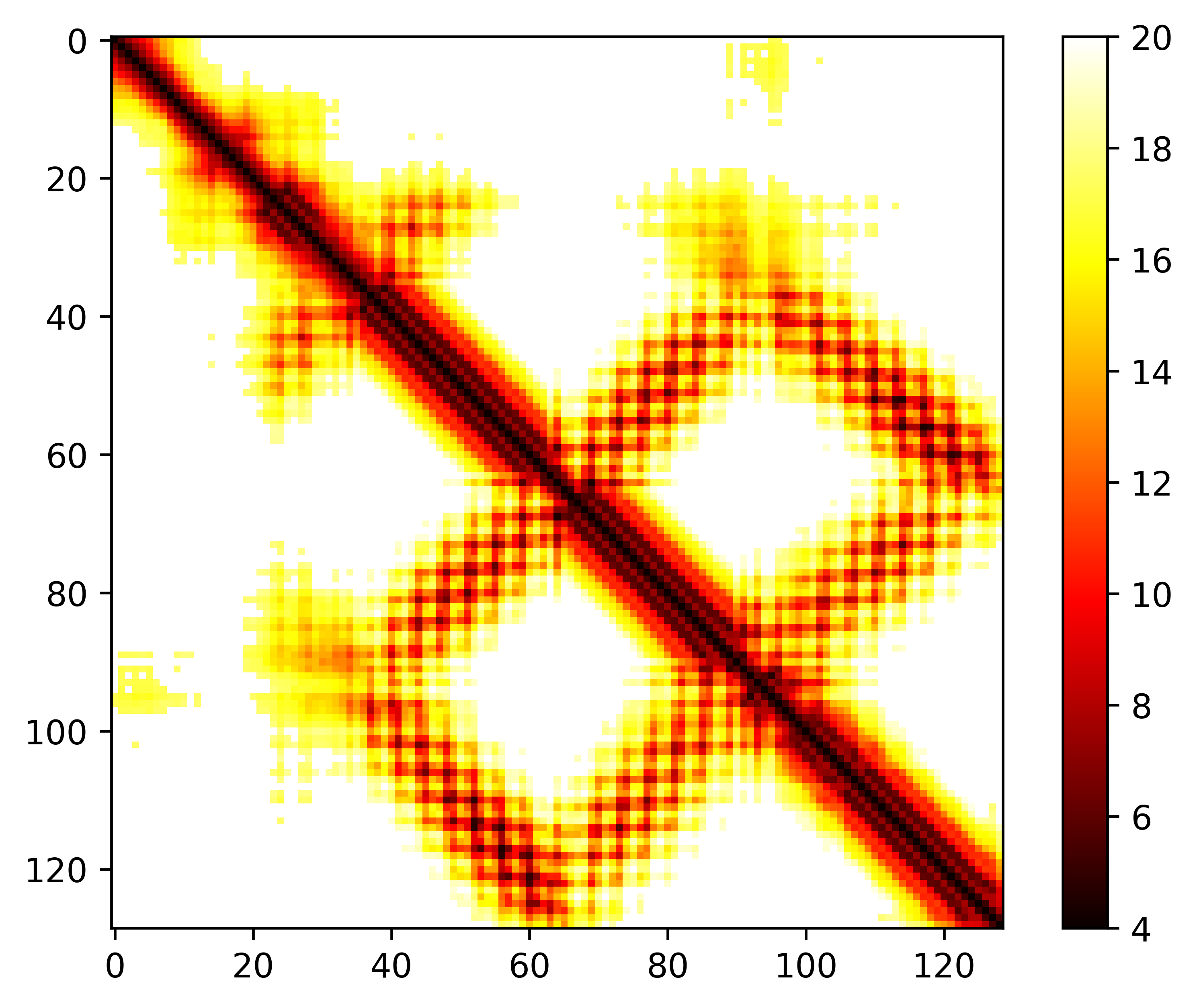

### seq.dist.png

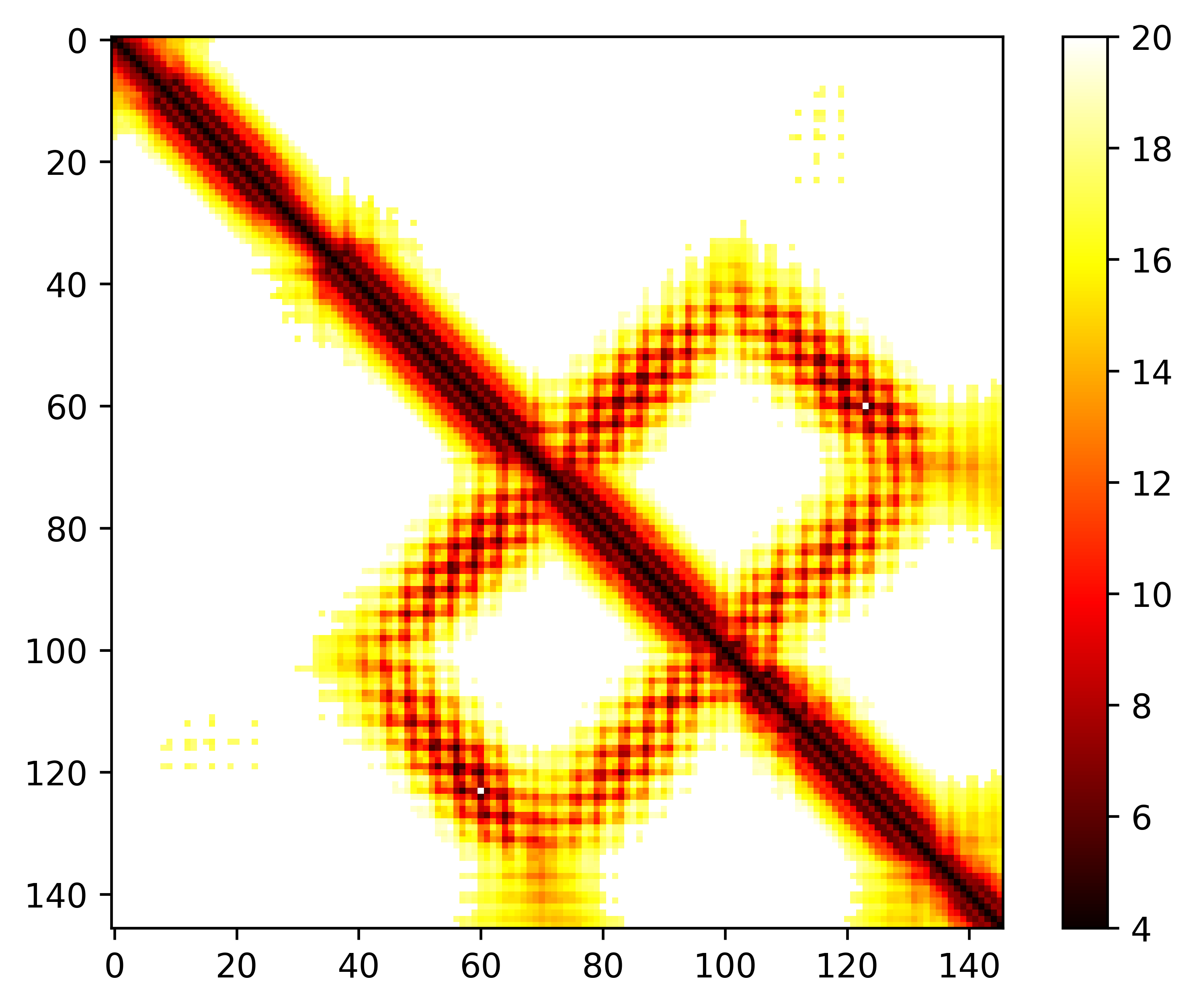

### seq.dist.png

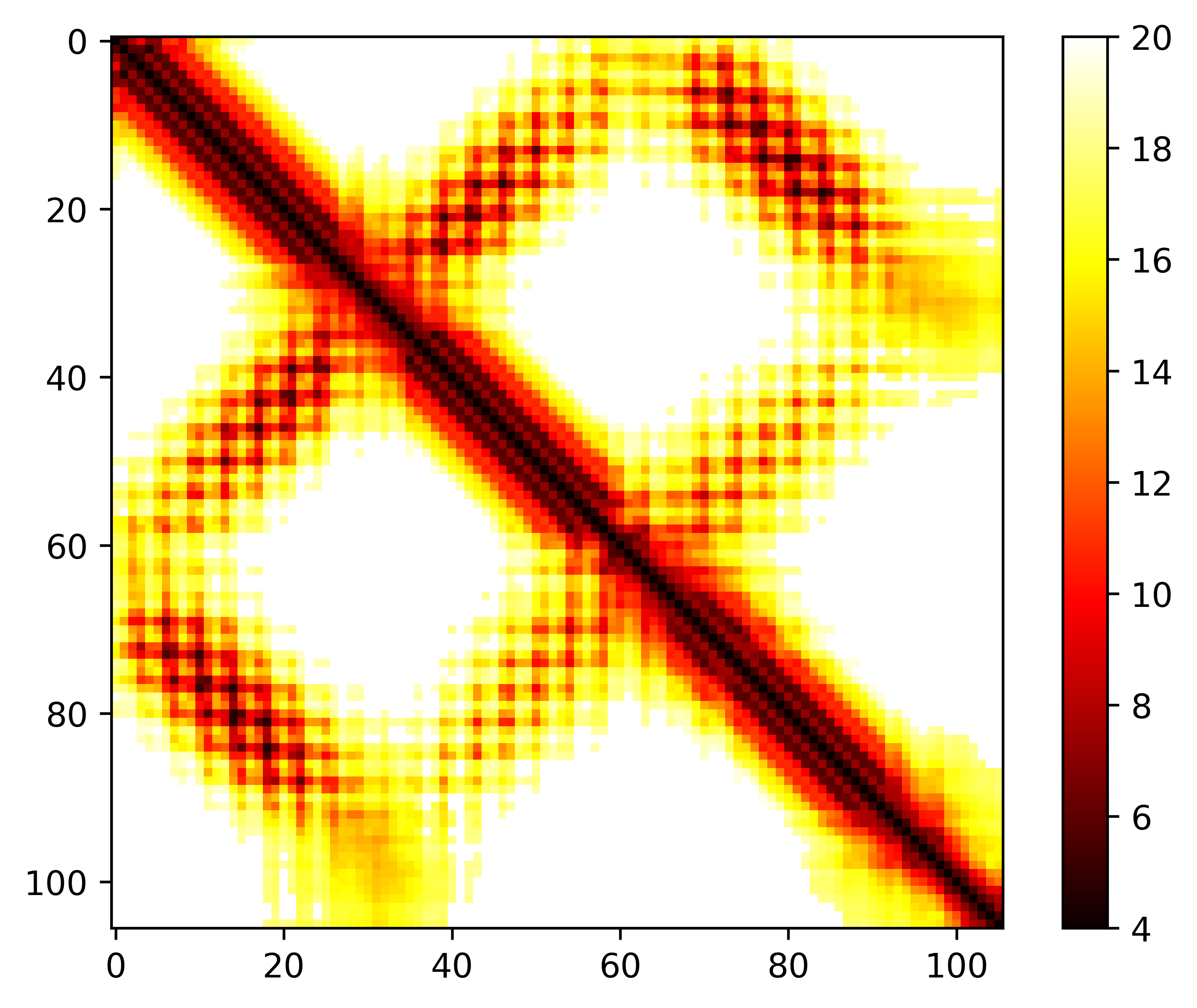

### seq.dist.png

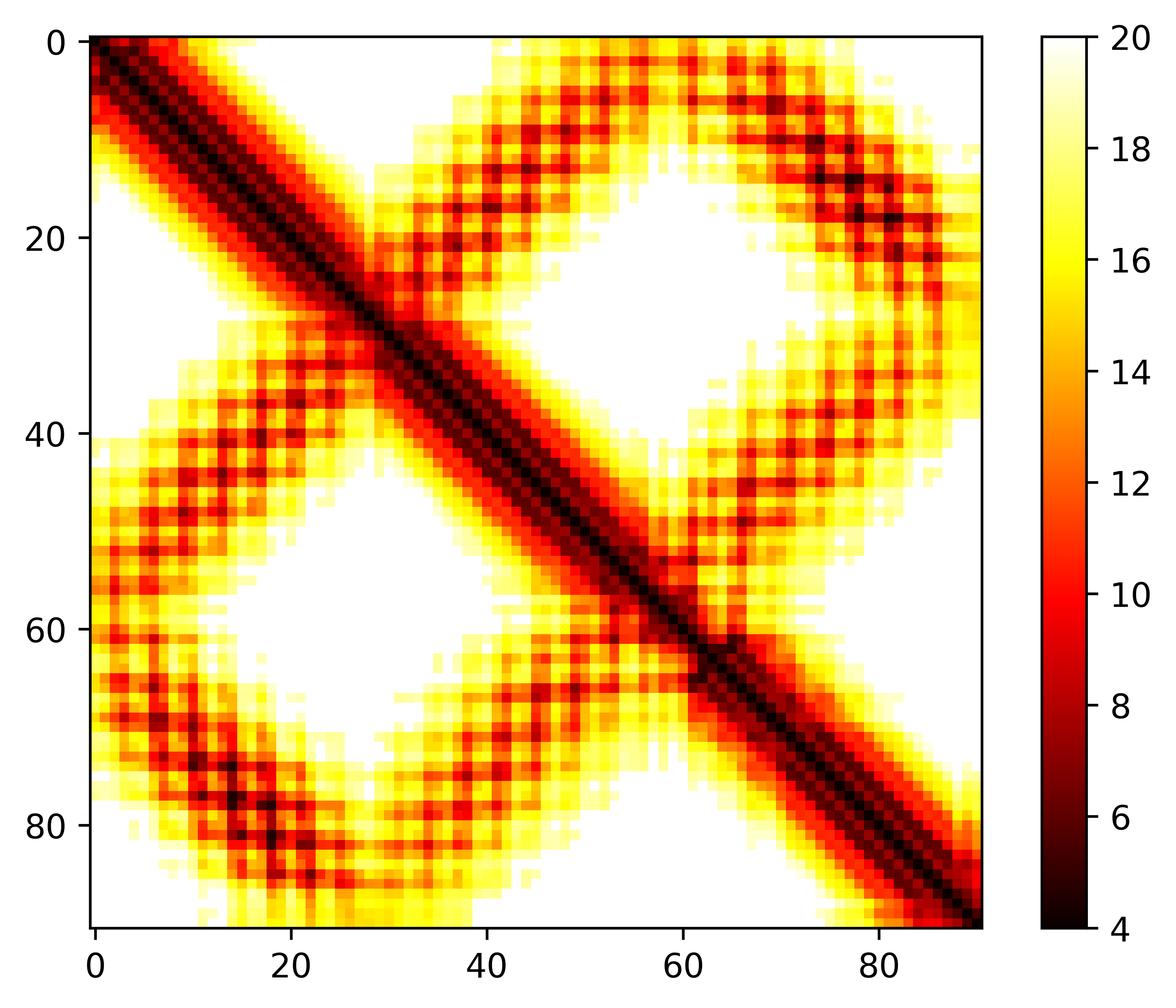

### seq.dist.png

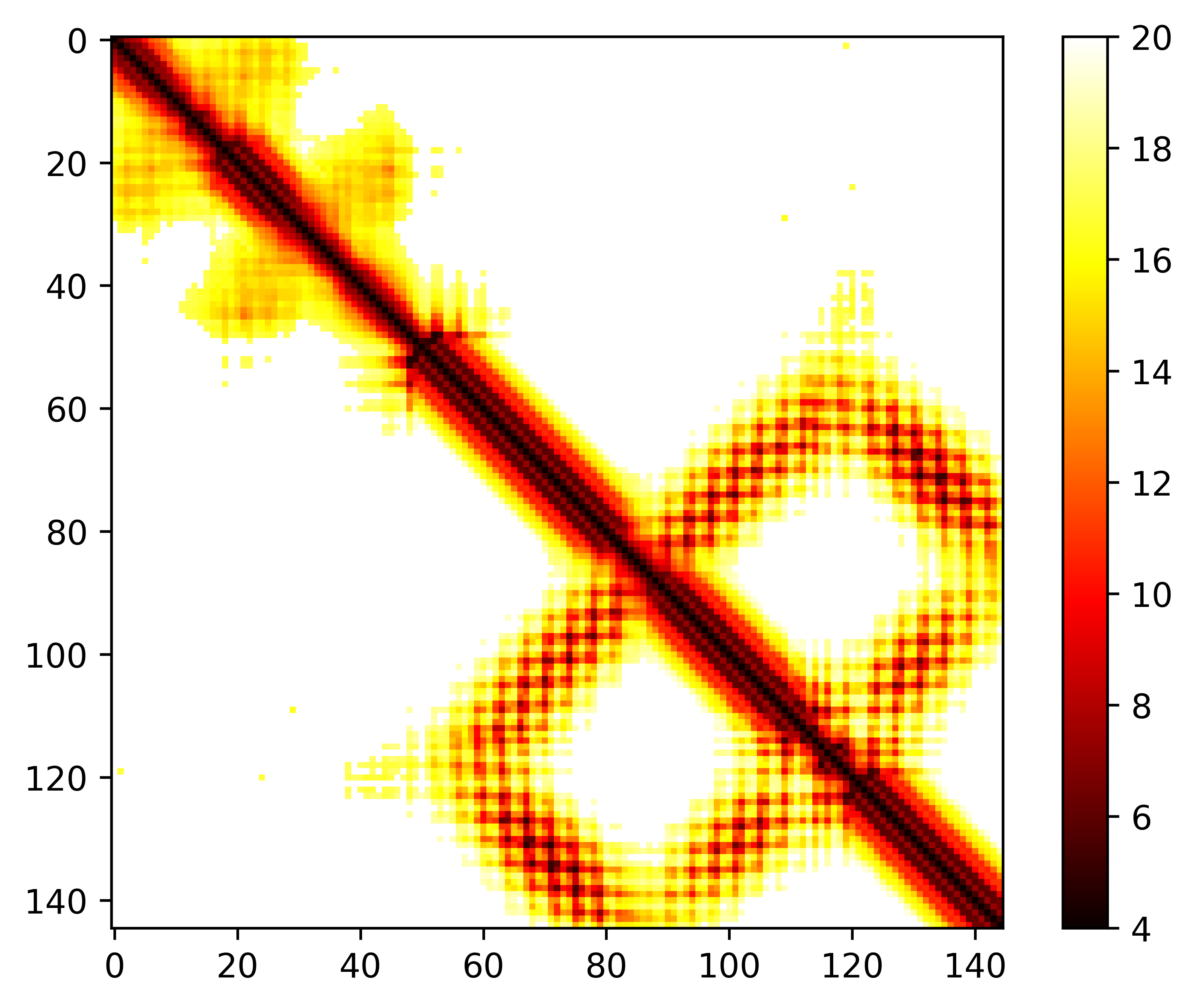

### seq.dist.png

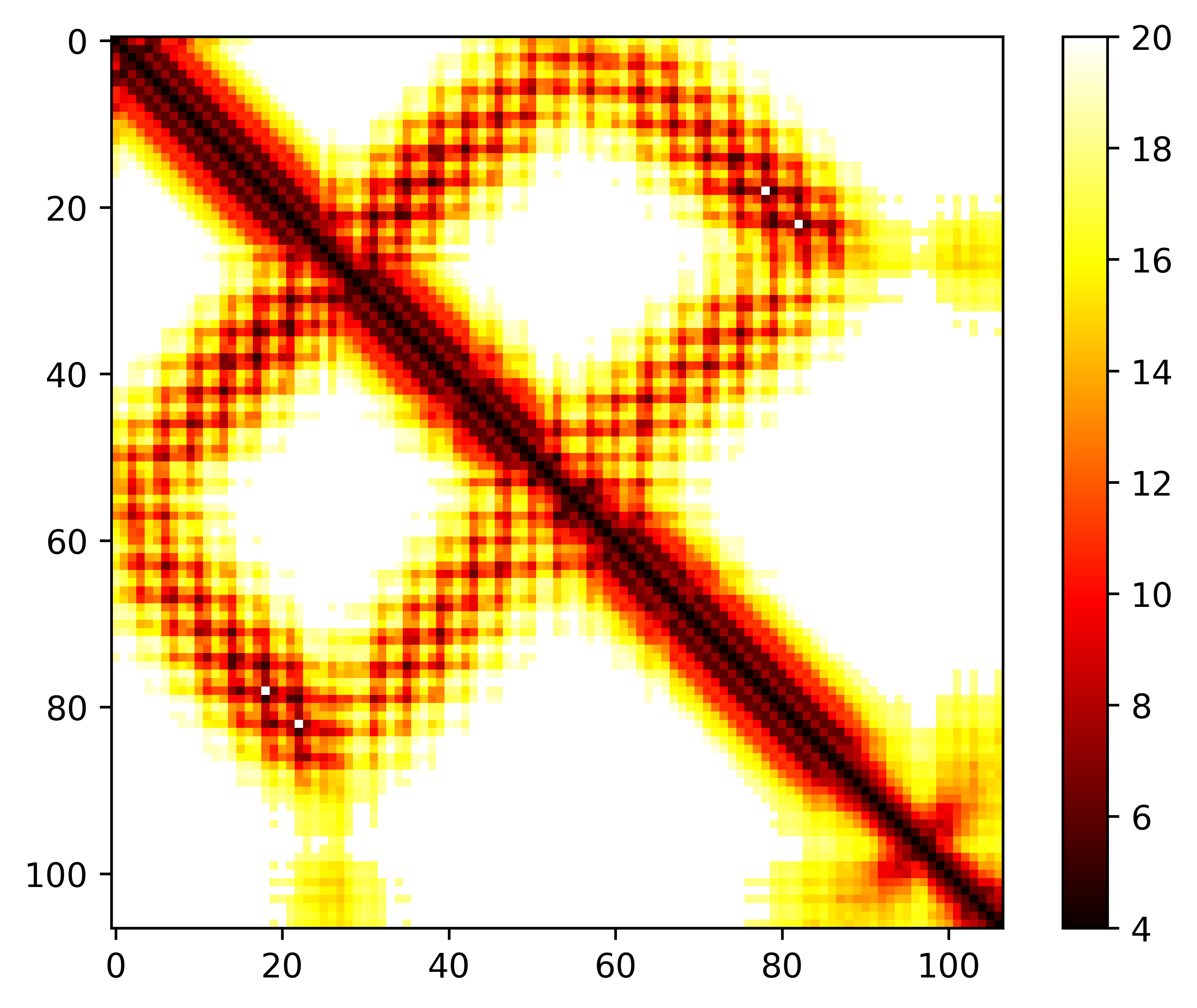

### seq.dist.png

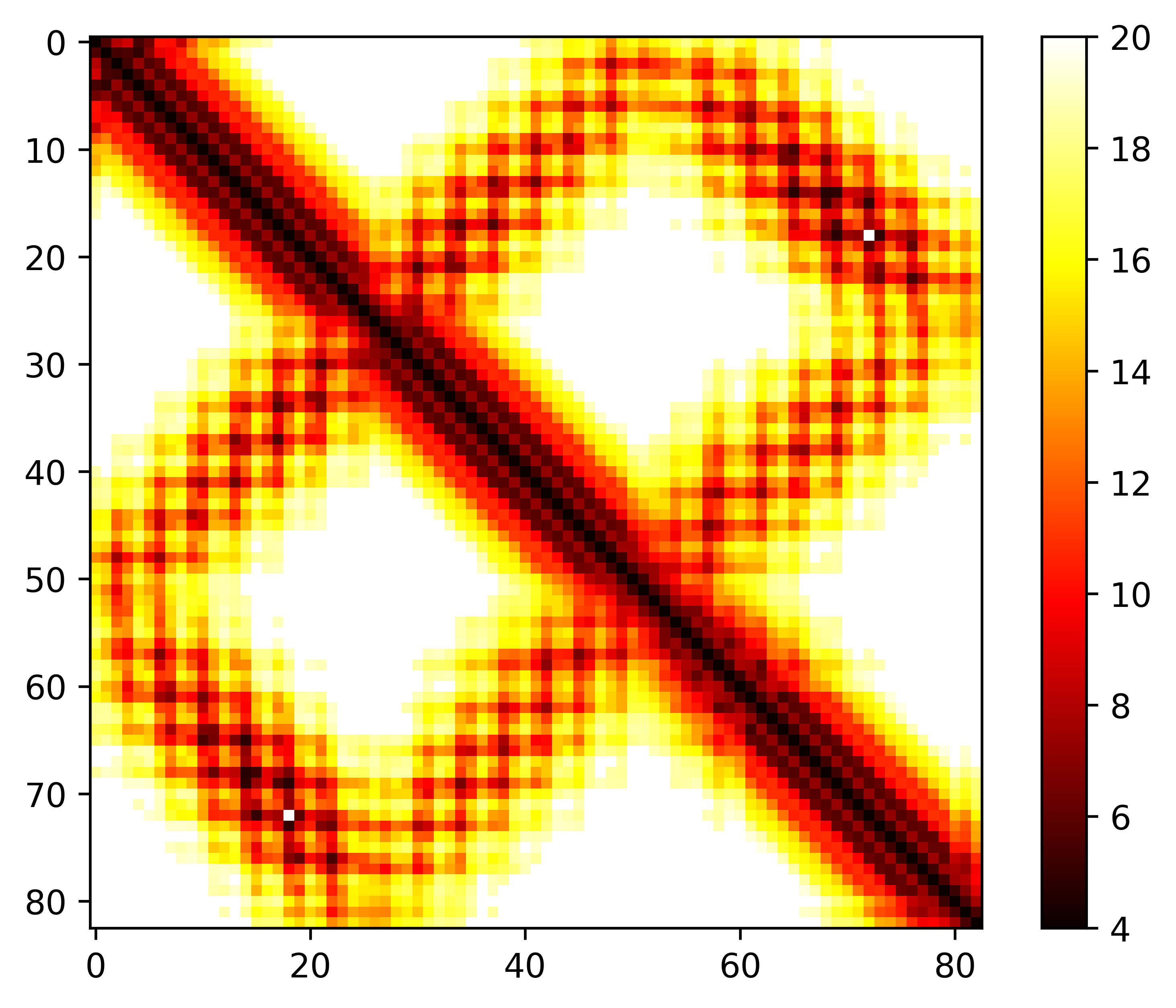

### seq.dist.png

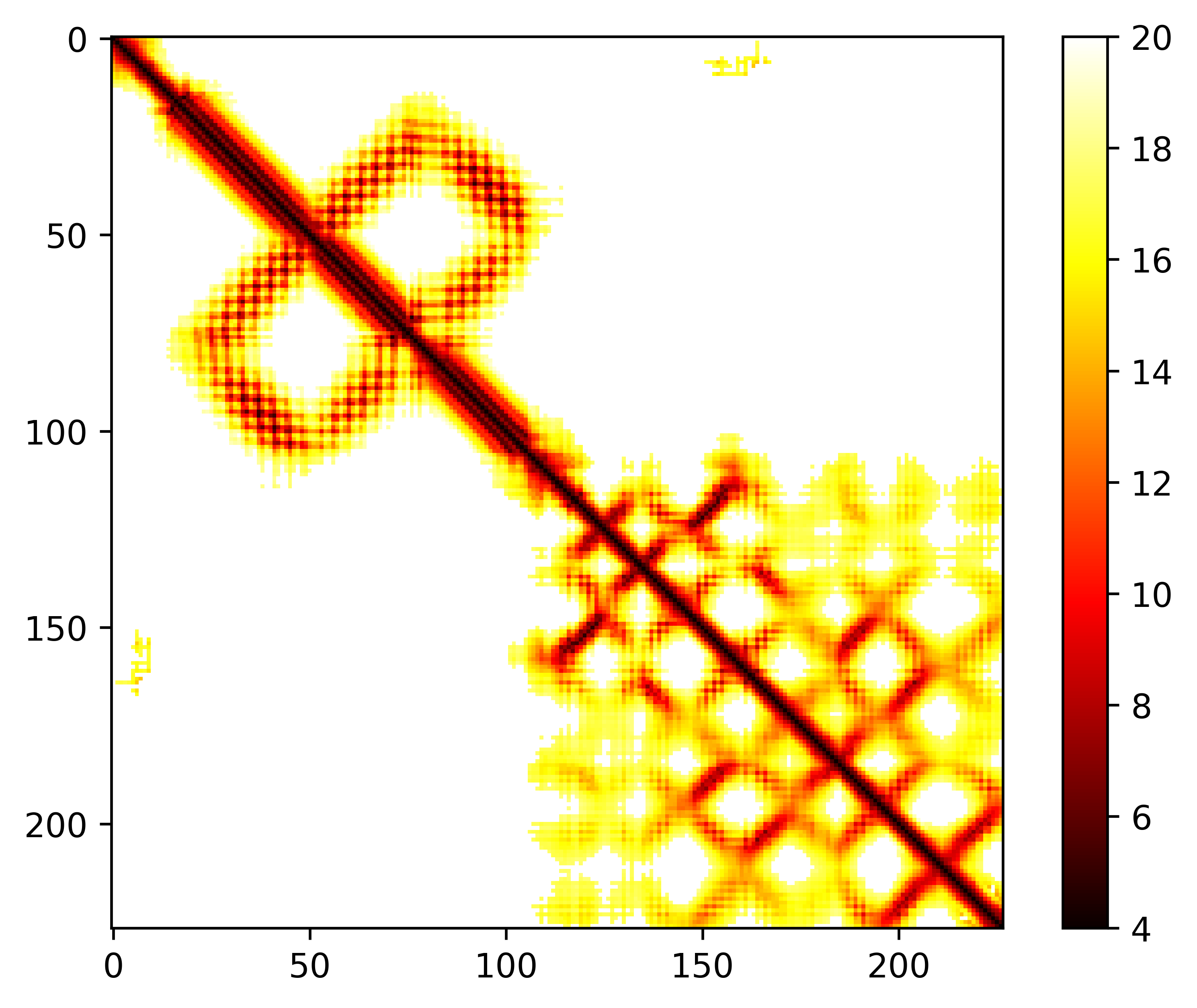

### seq.omega.png

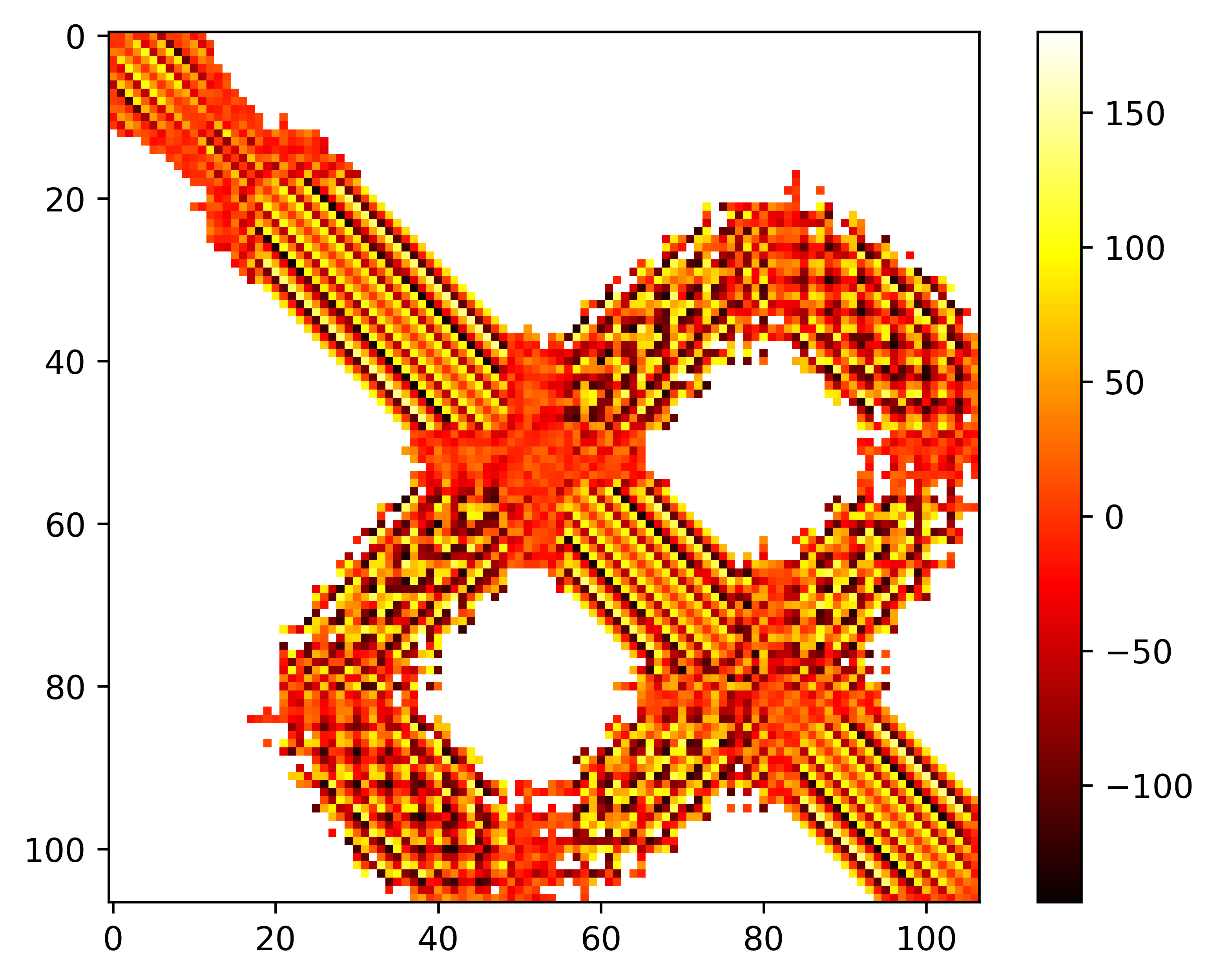

### seq.omega.png

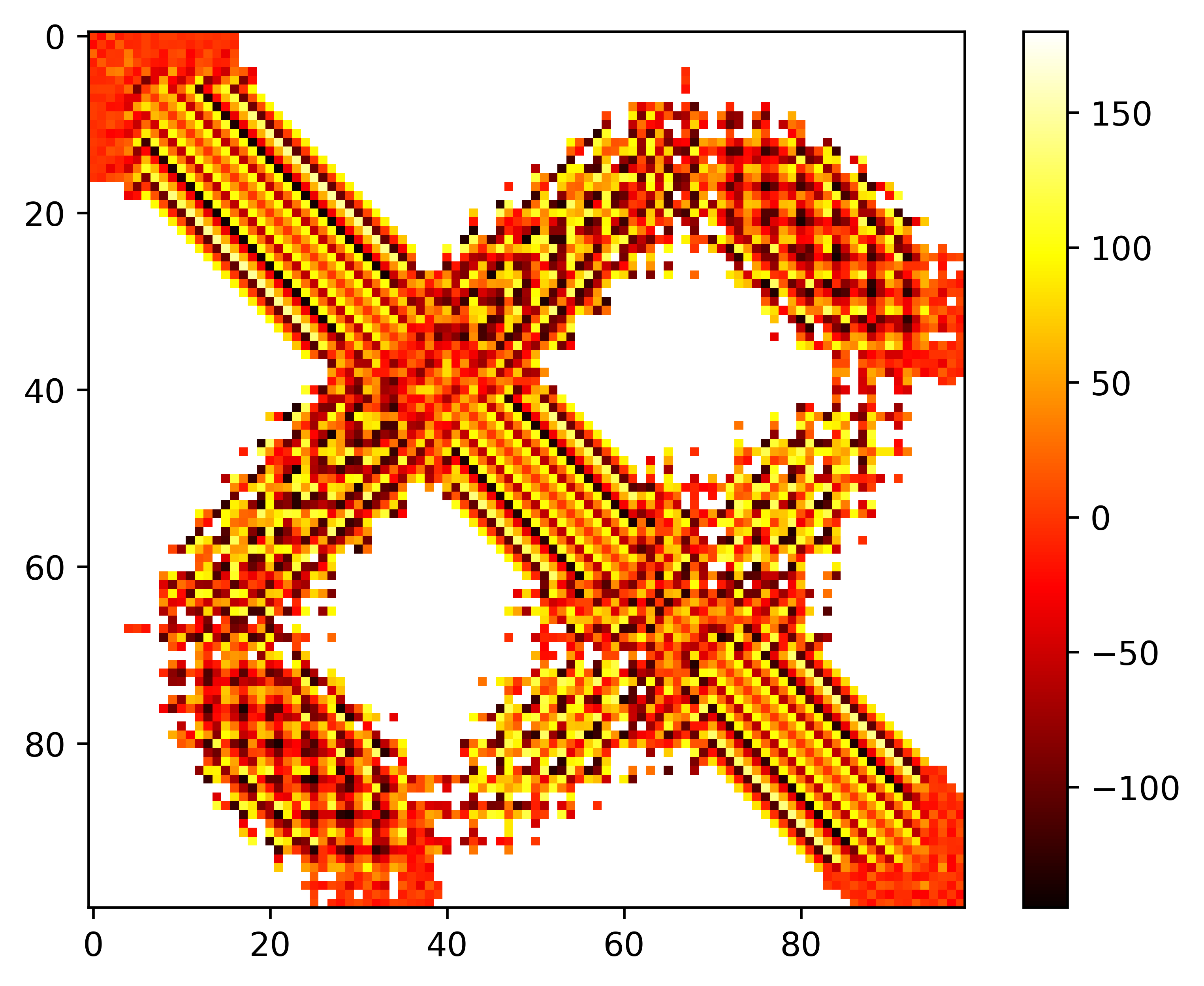

### seq.omega.png

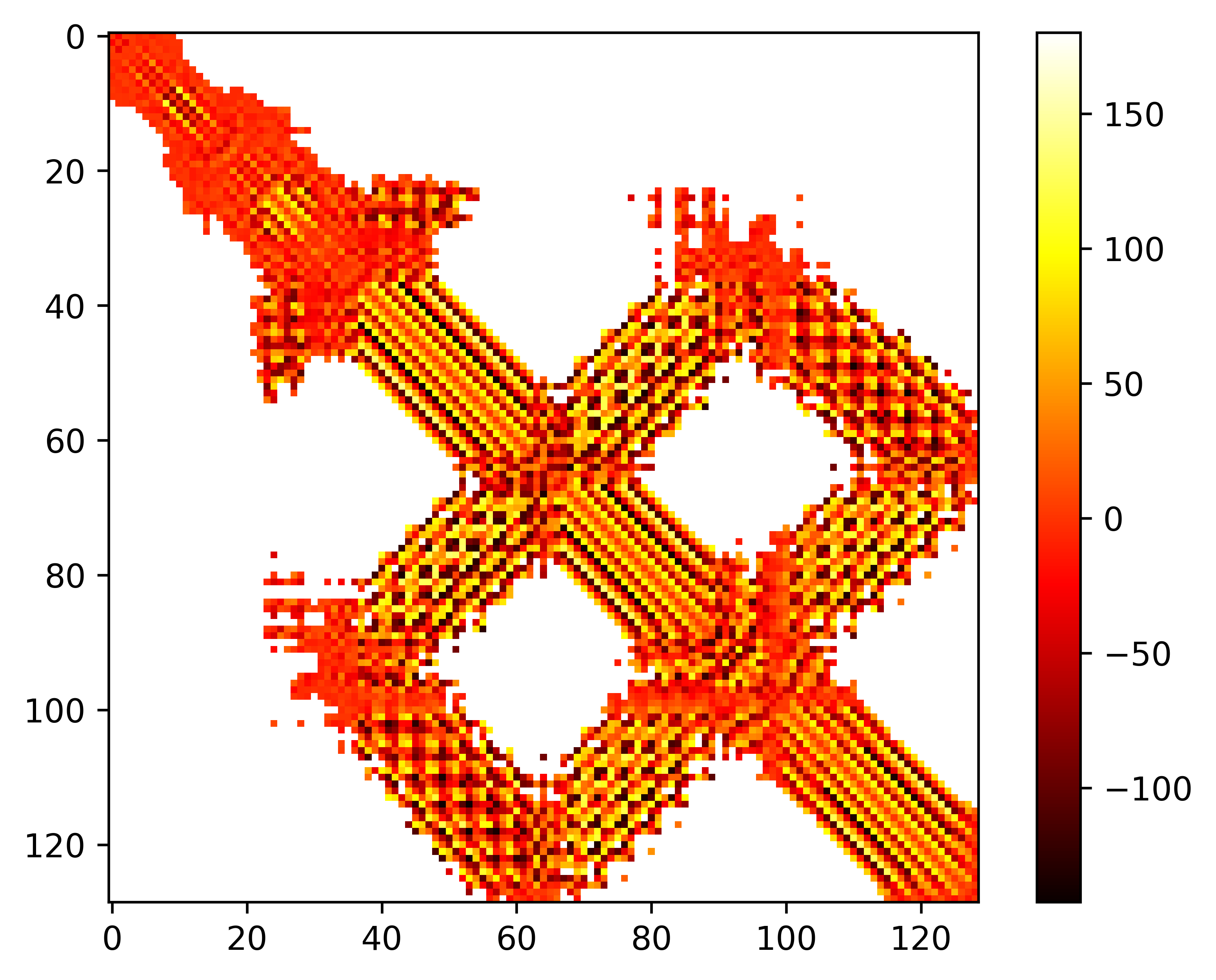

### seq.omega.png

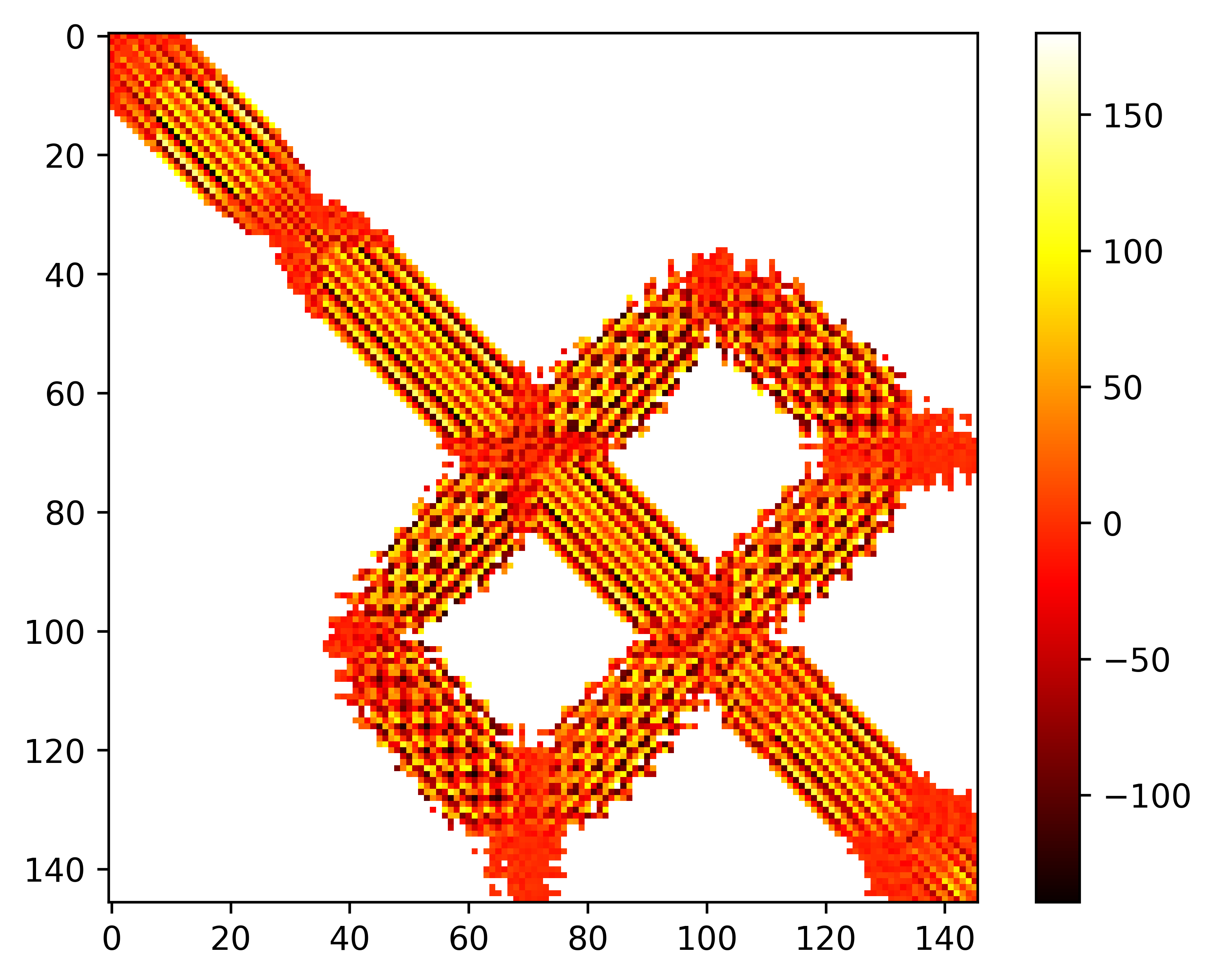

### seq.omega.png

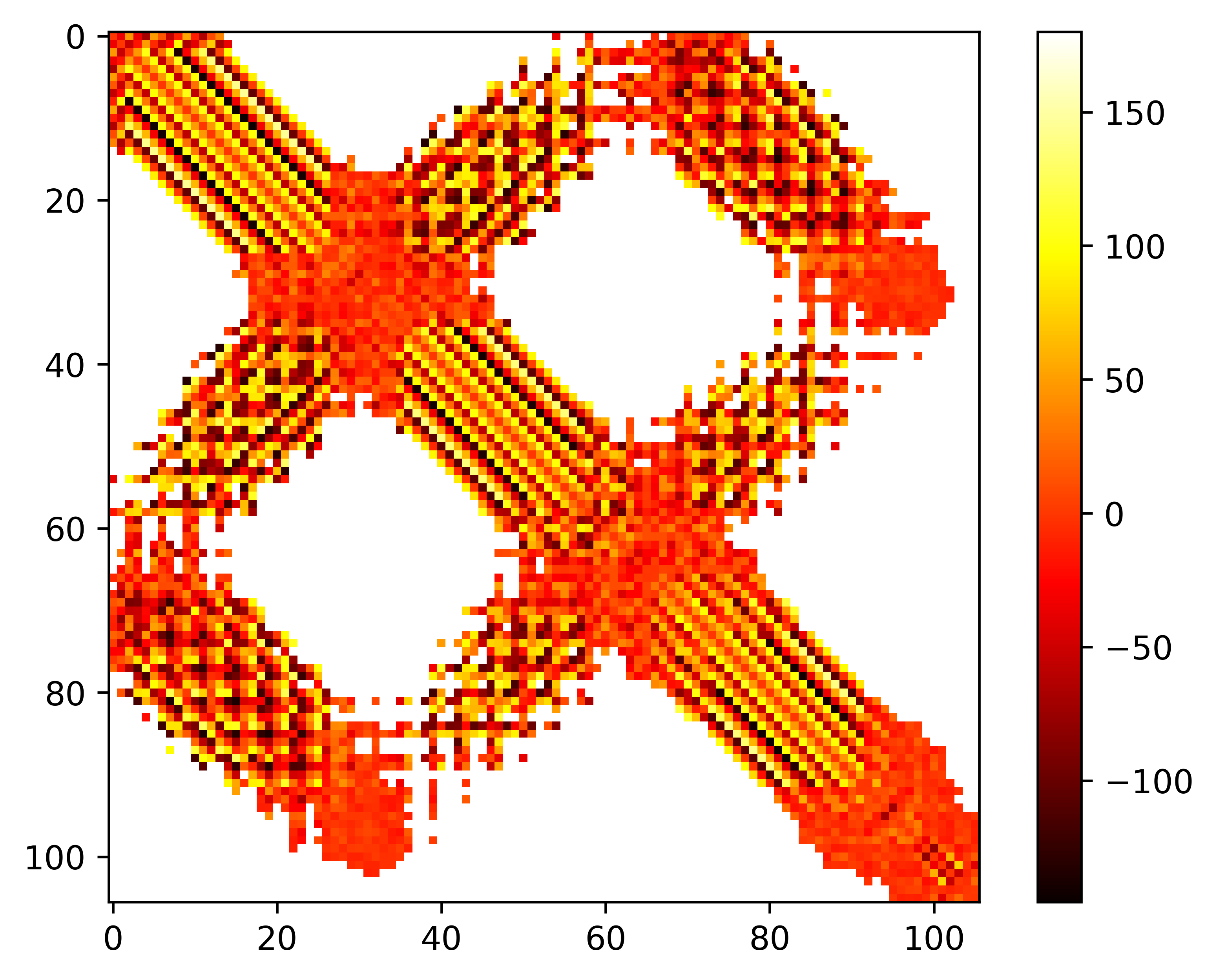

### seq.omega.png

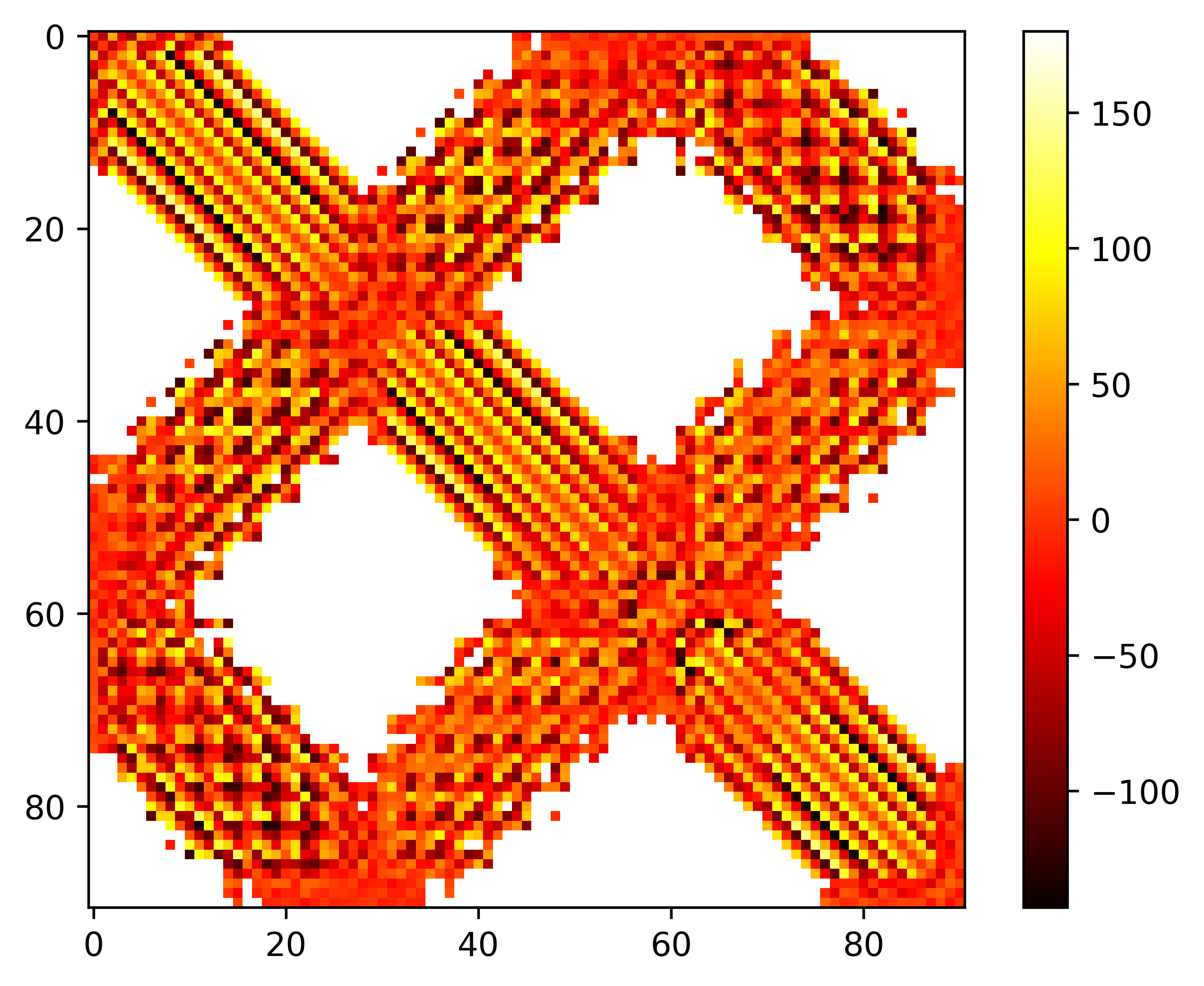

### seq.omega.png

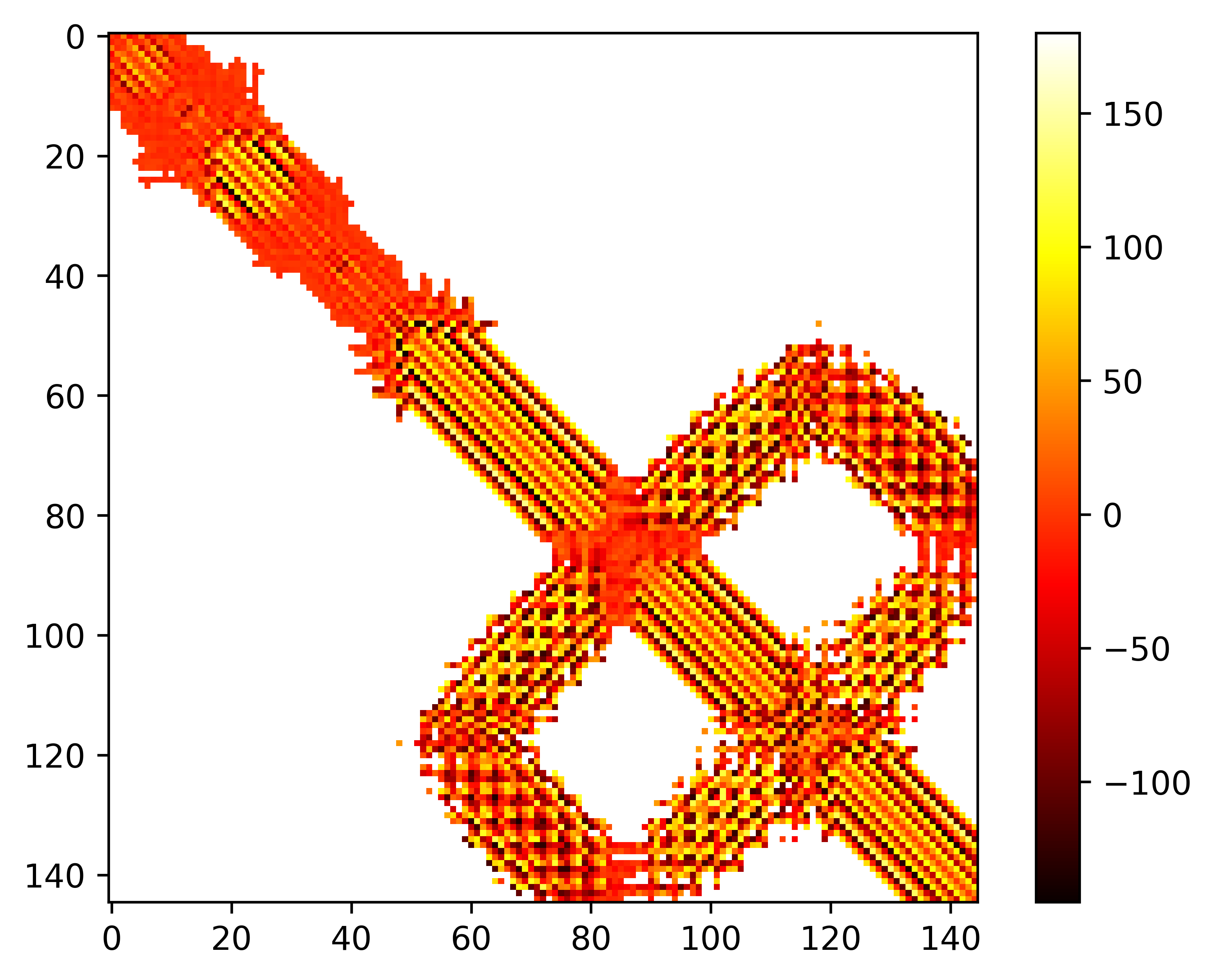

### seq.omega.png

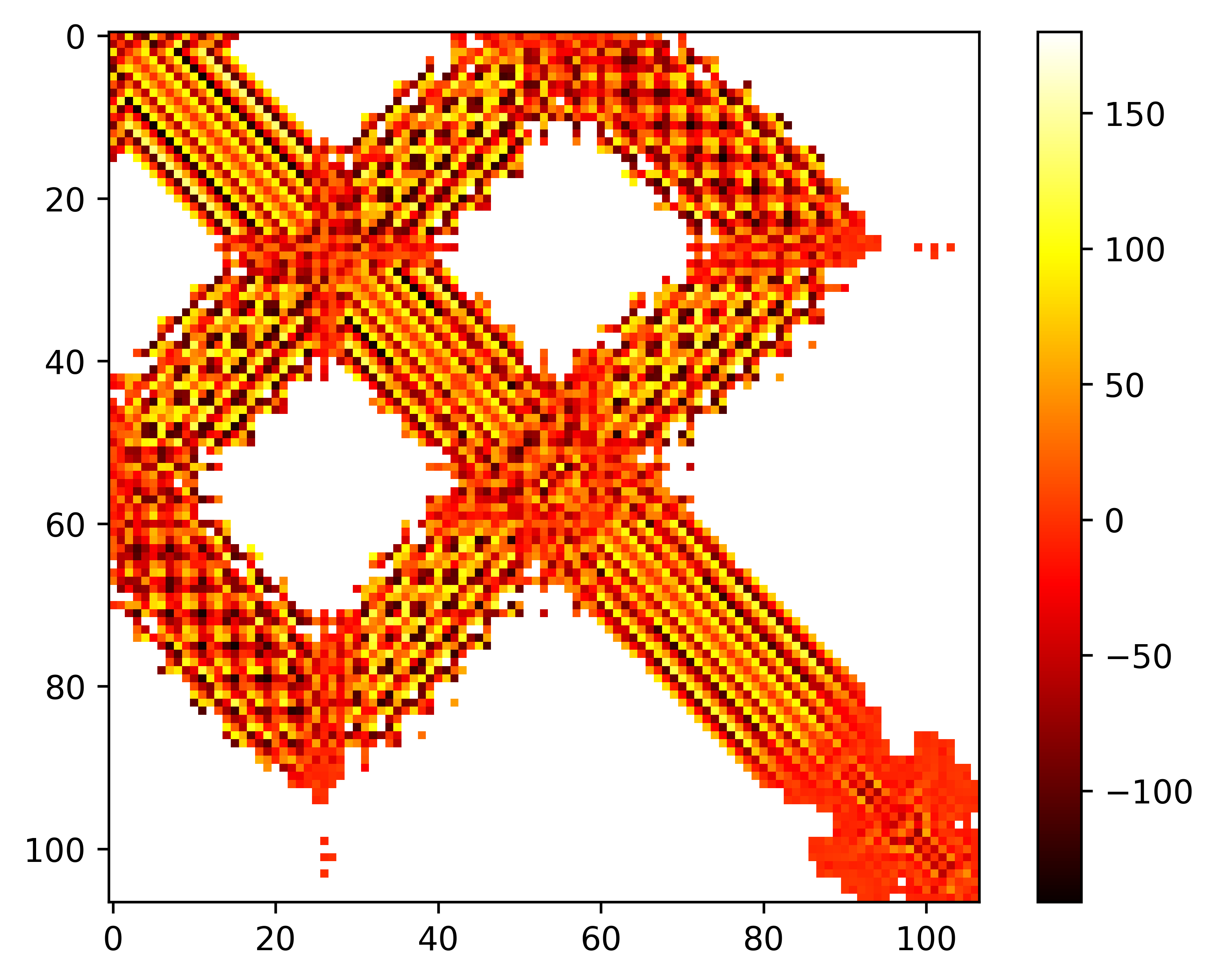

### seq.omega.png

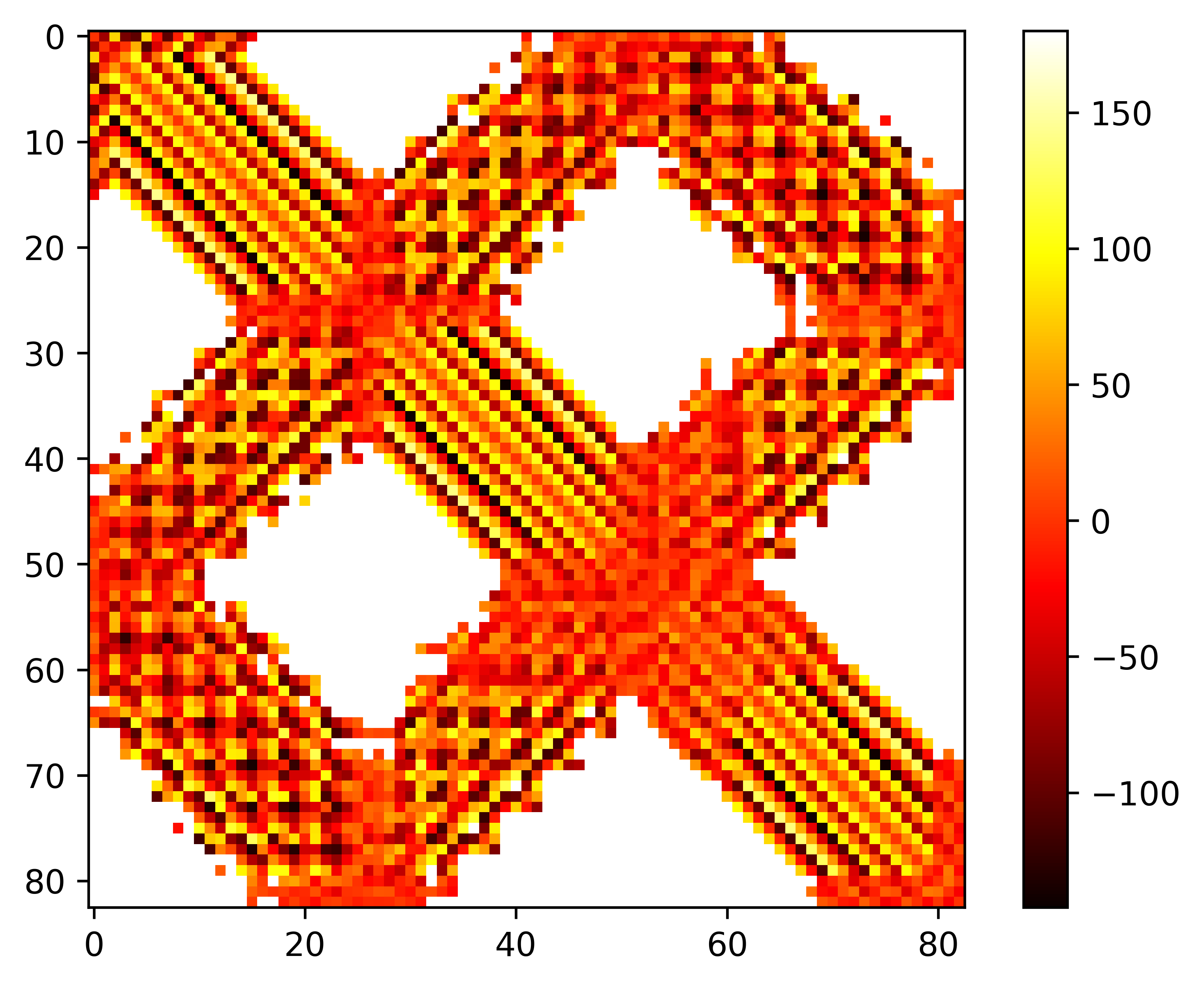

### seq.omega.png

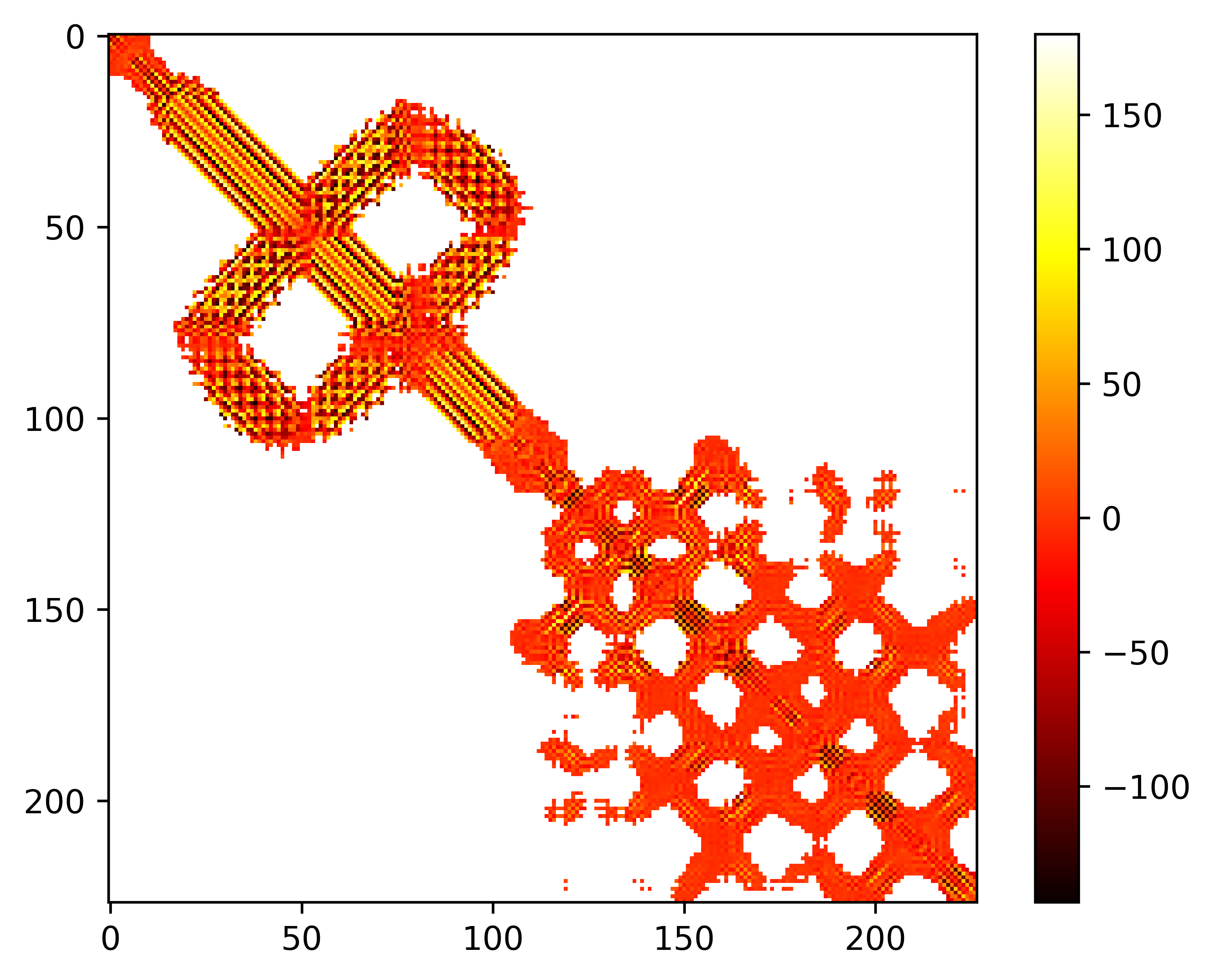

### seq.phi.png

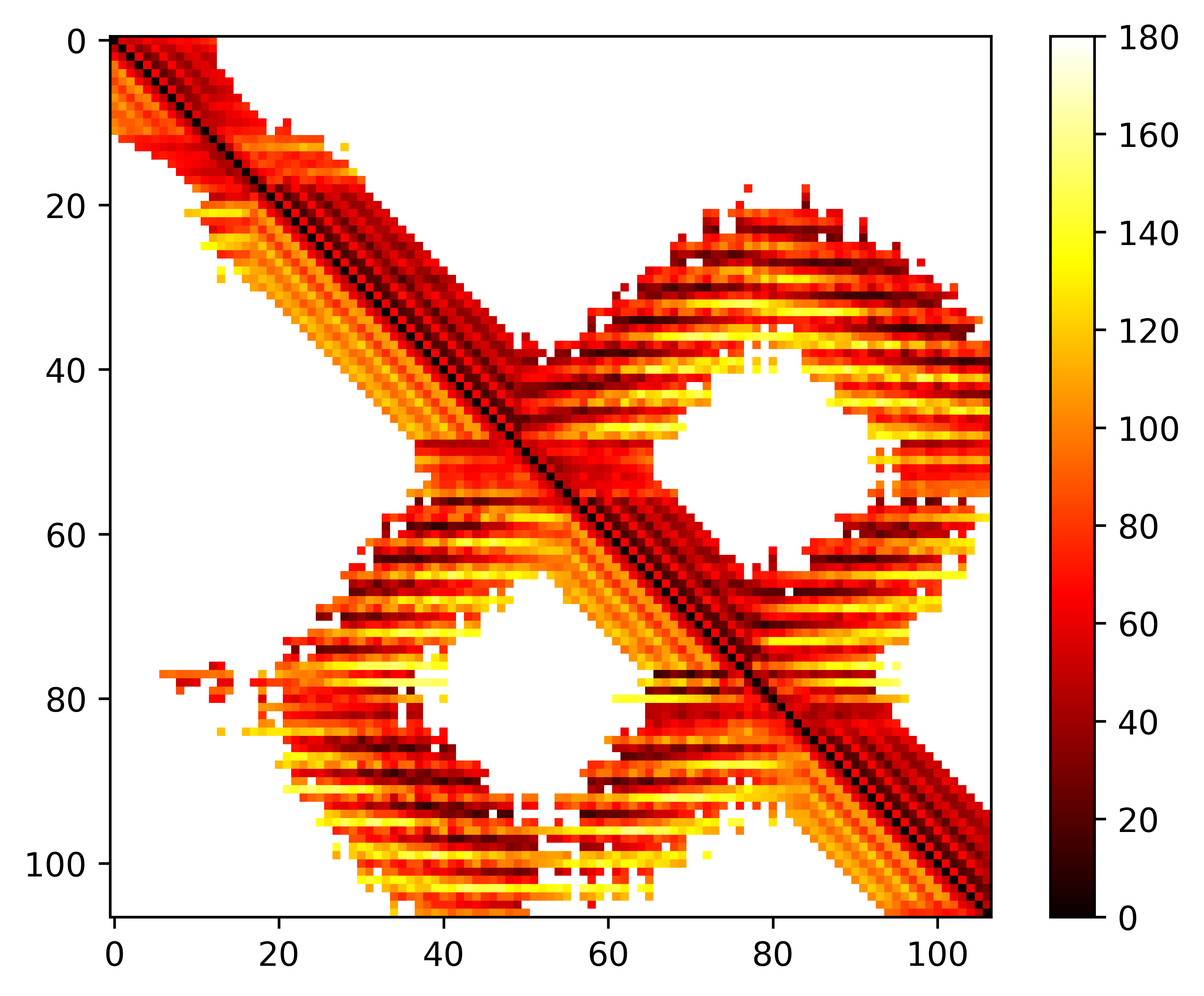

### seq.phi.png

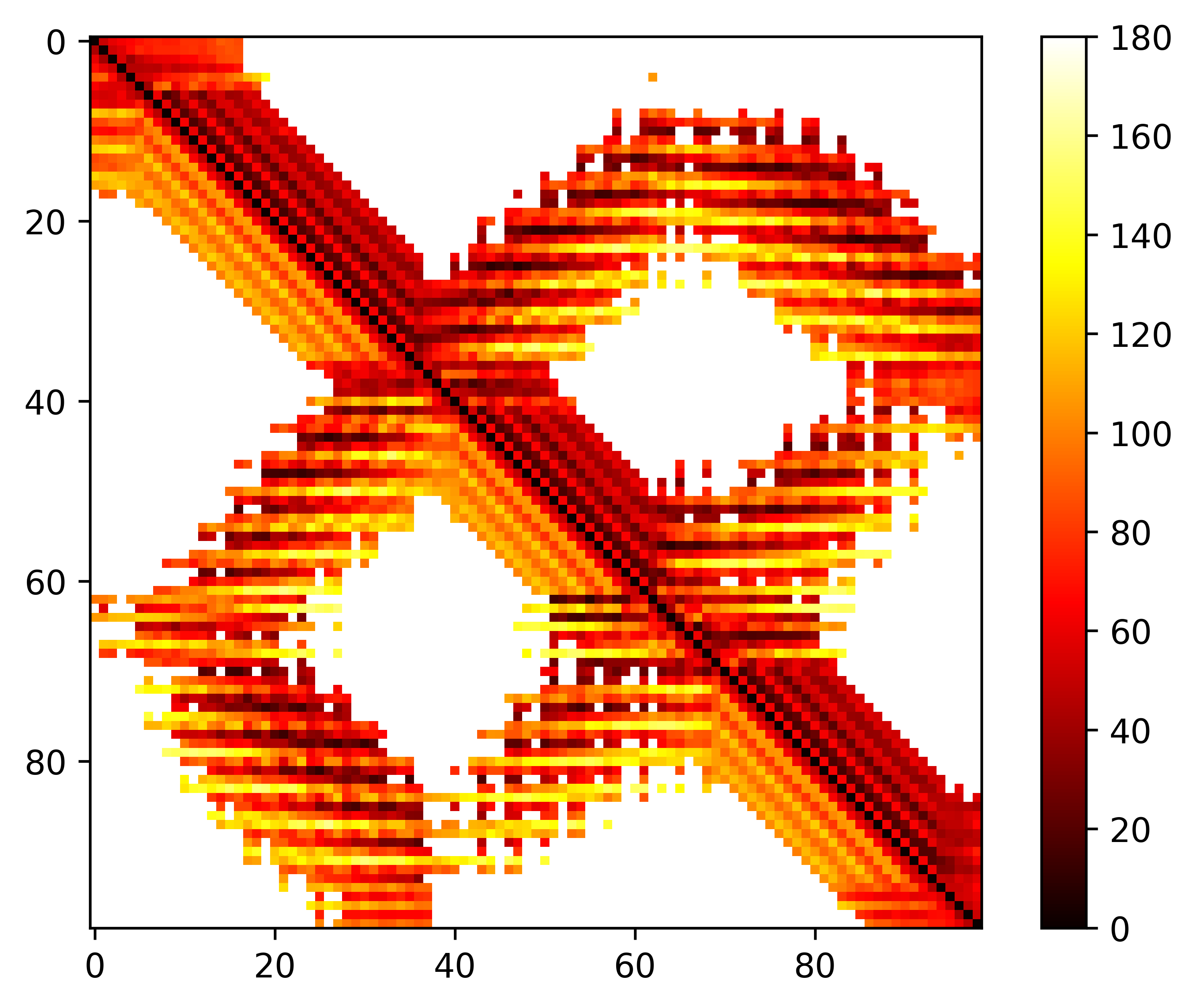

### seq.phi.png

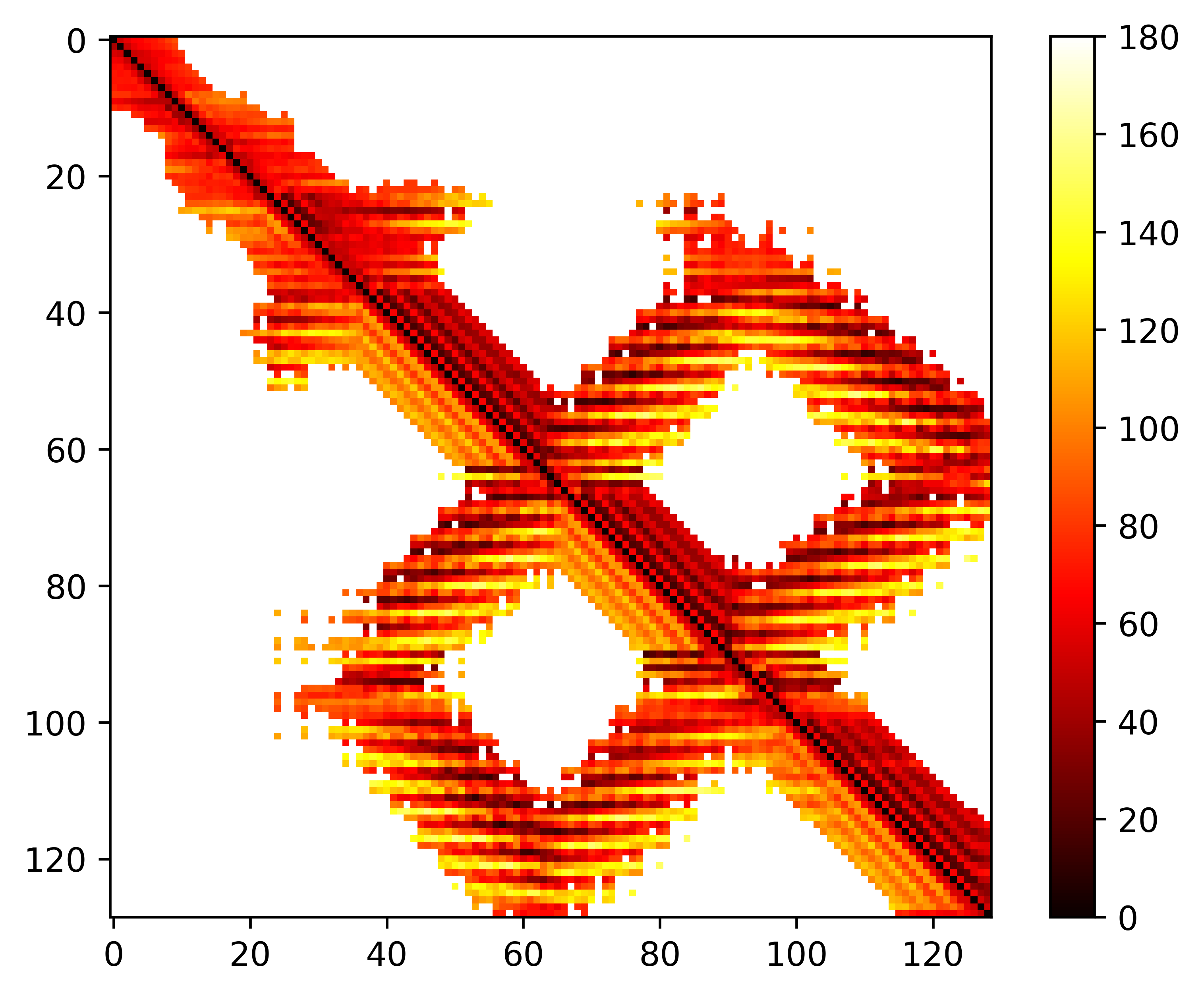

### seq.phi.png

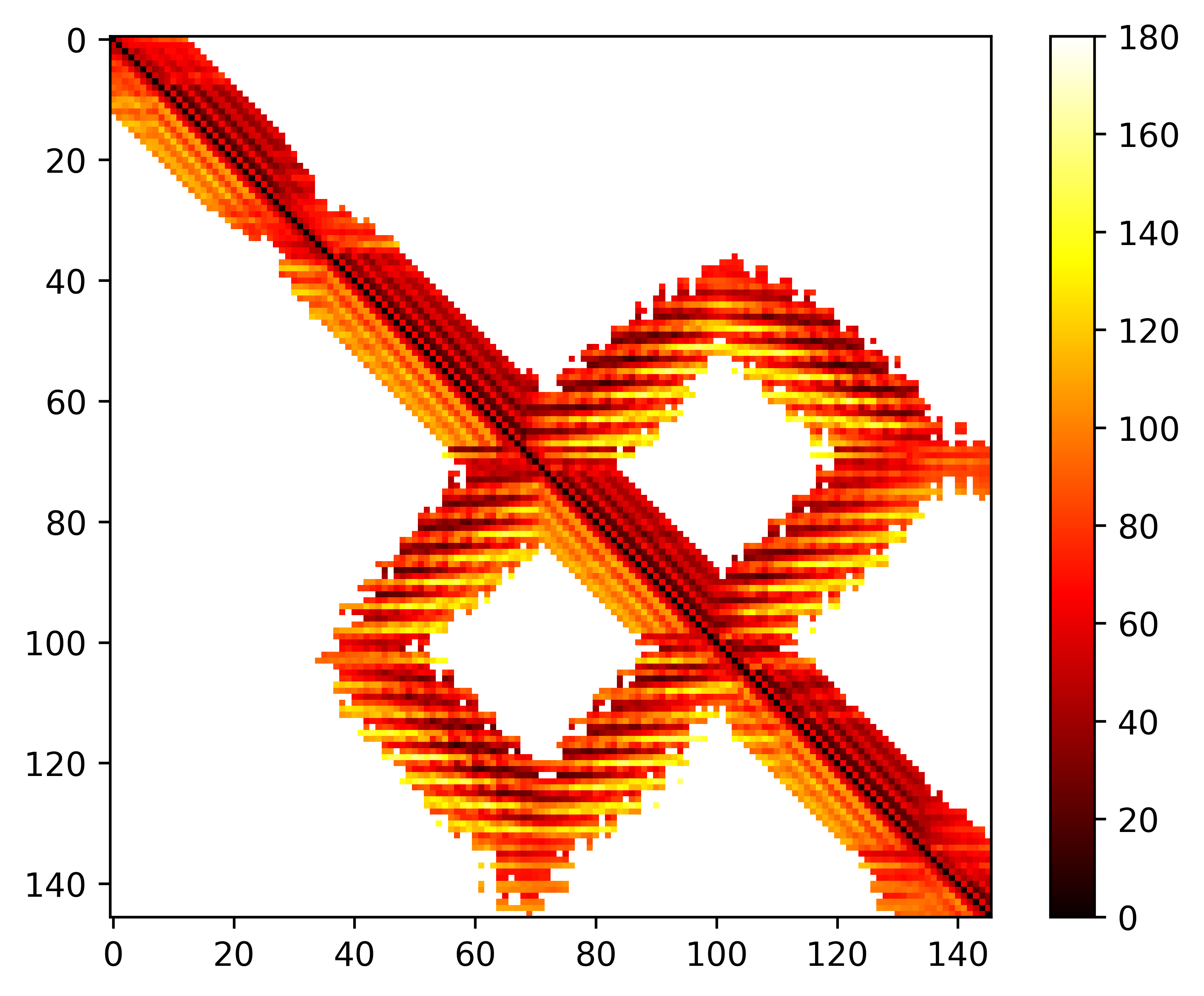

### seq.phi.png

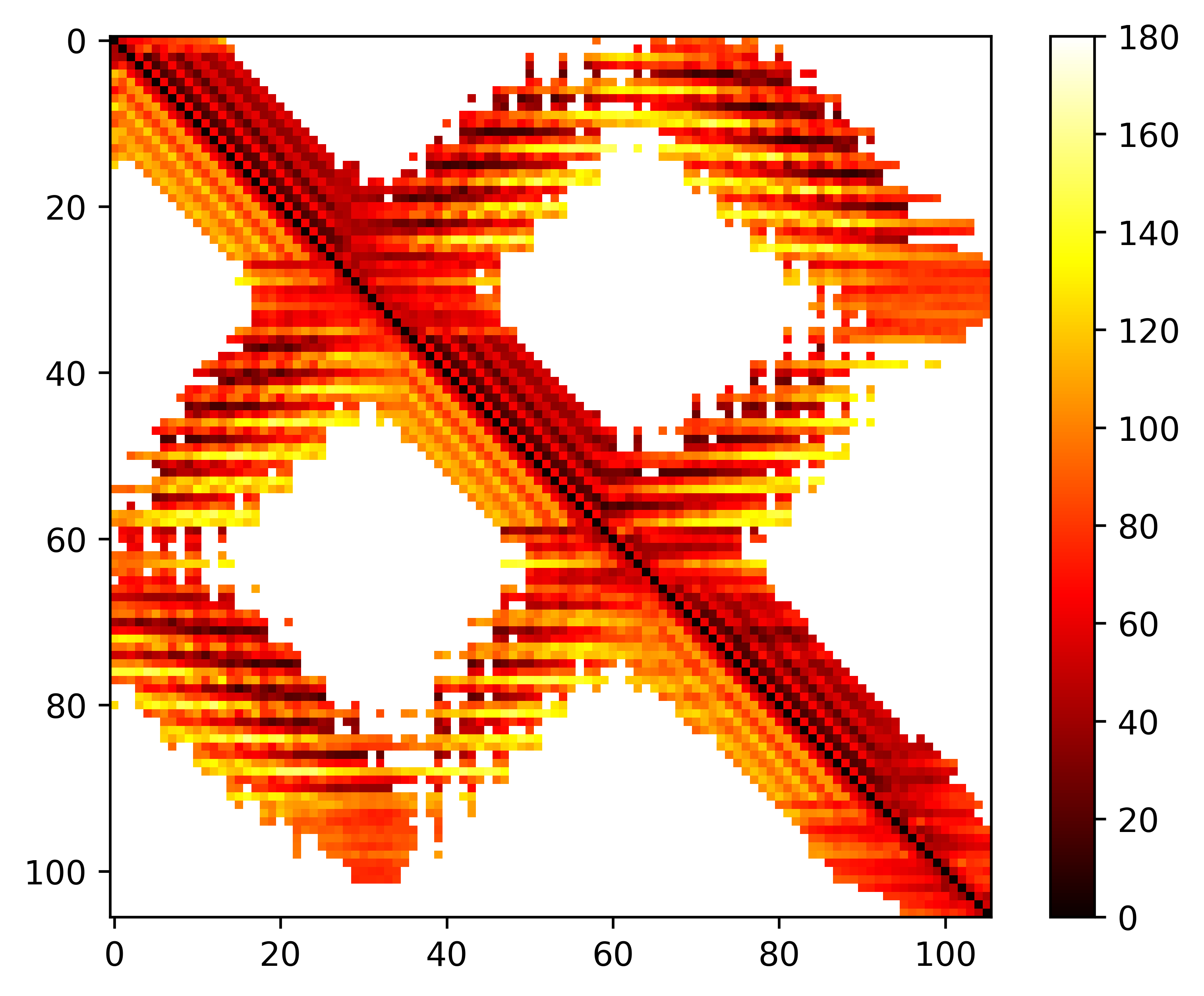

### seq.phi.png

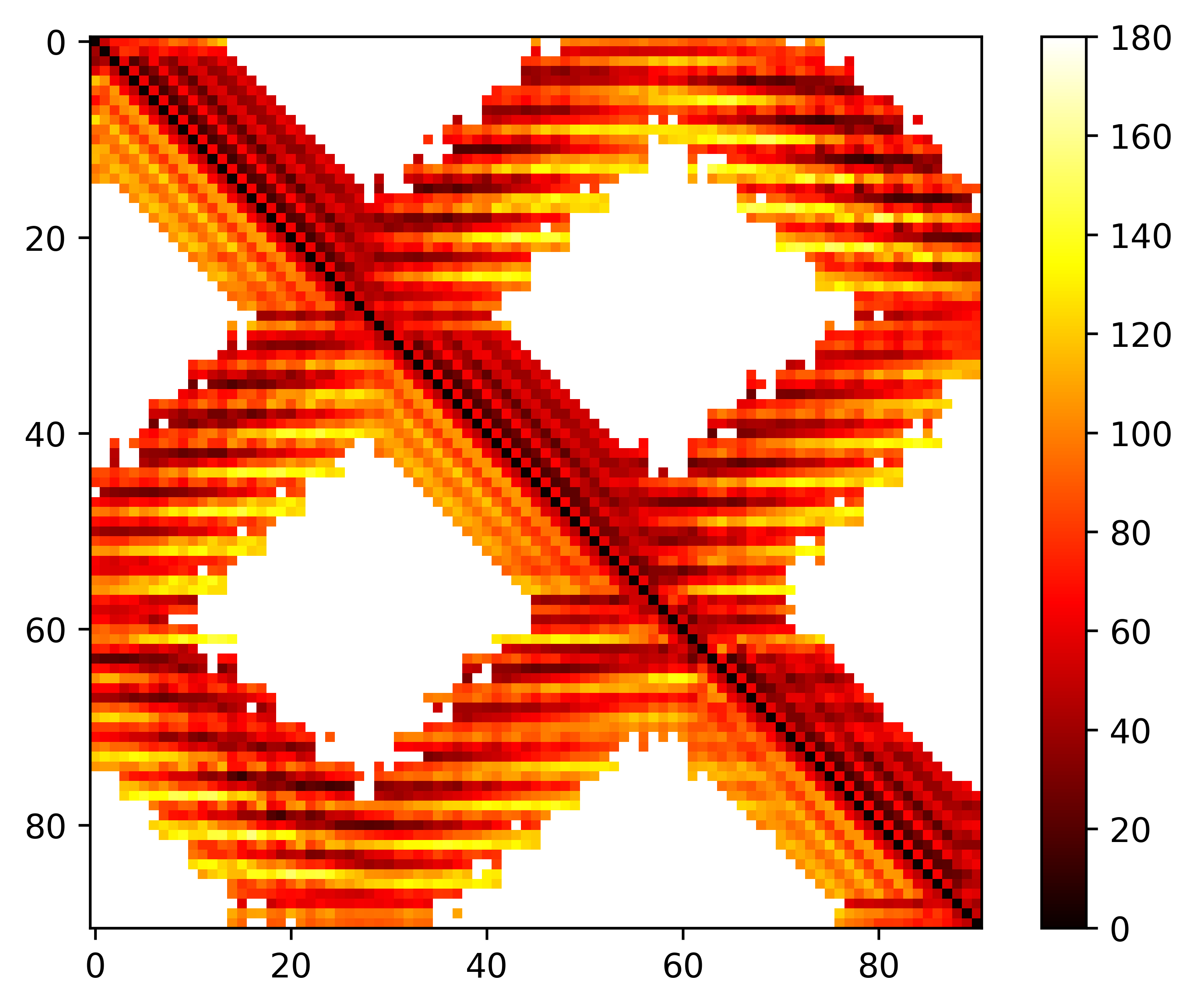

### seq.phi.png

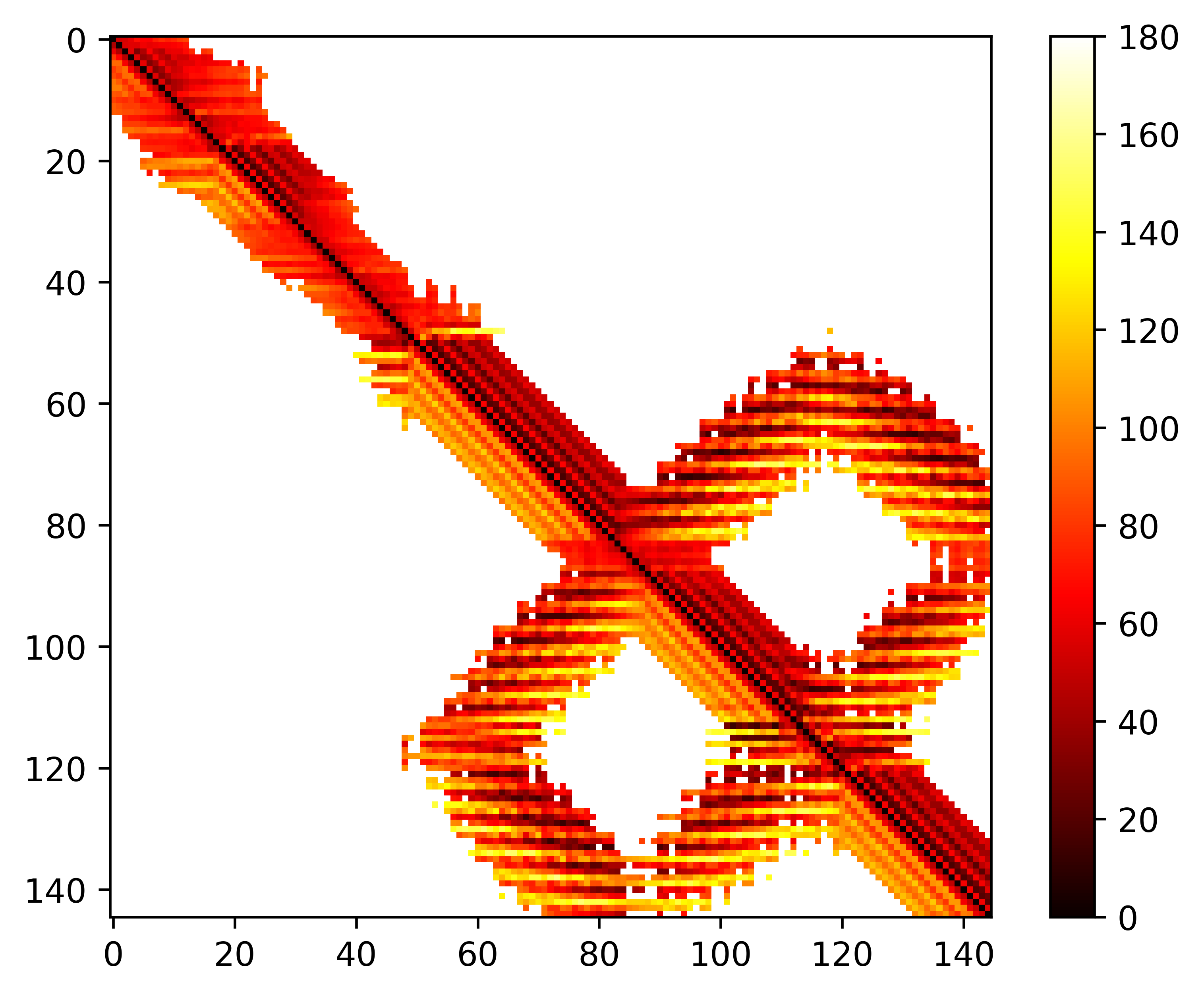

### seq.phi.png

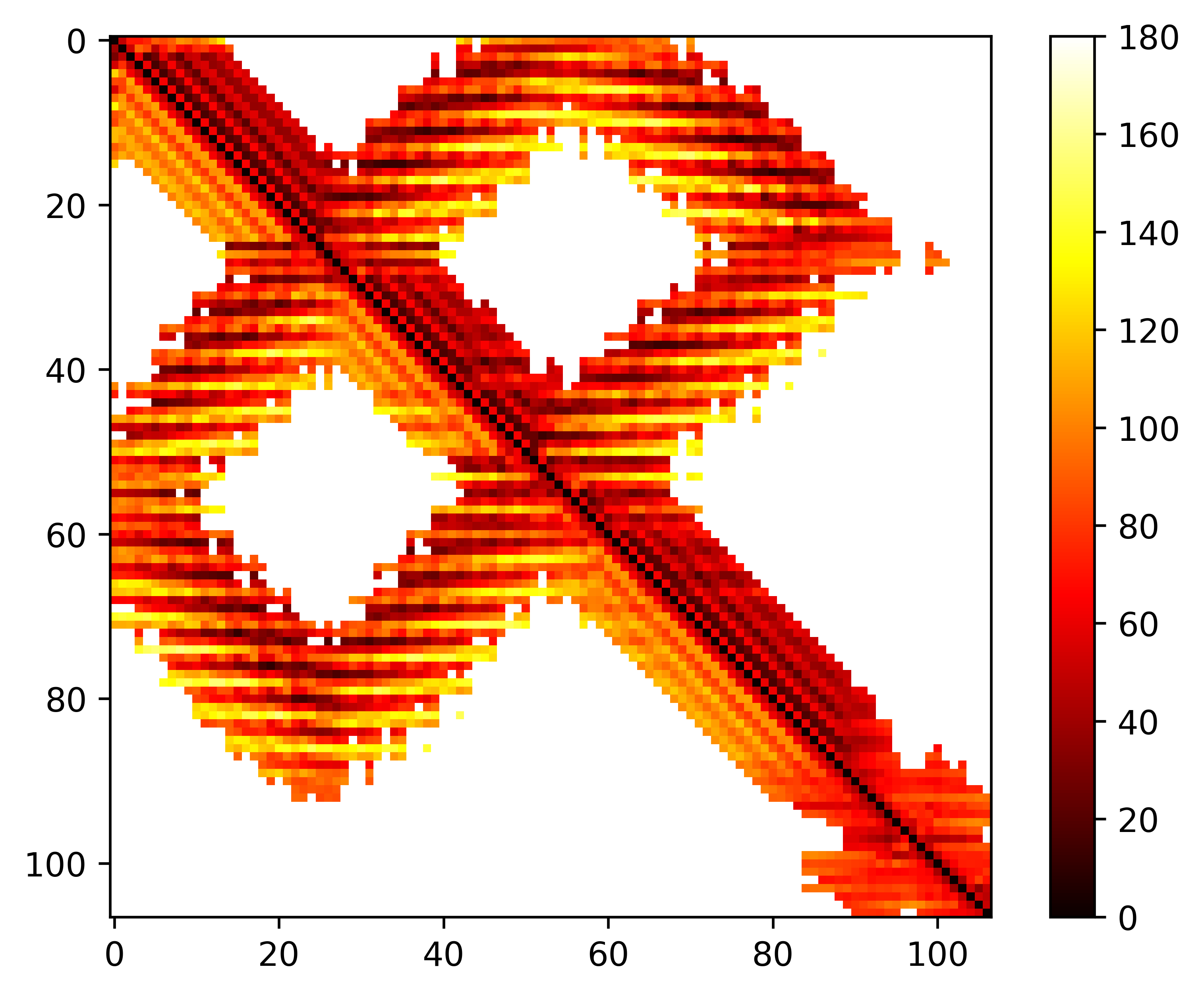

### seq.phi.png

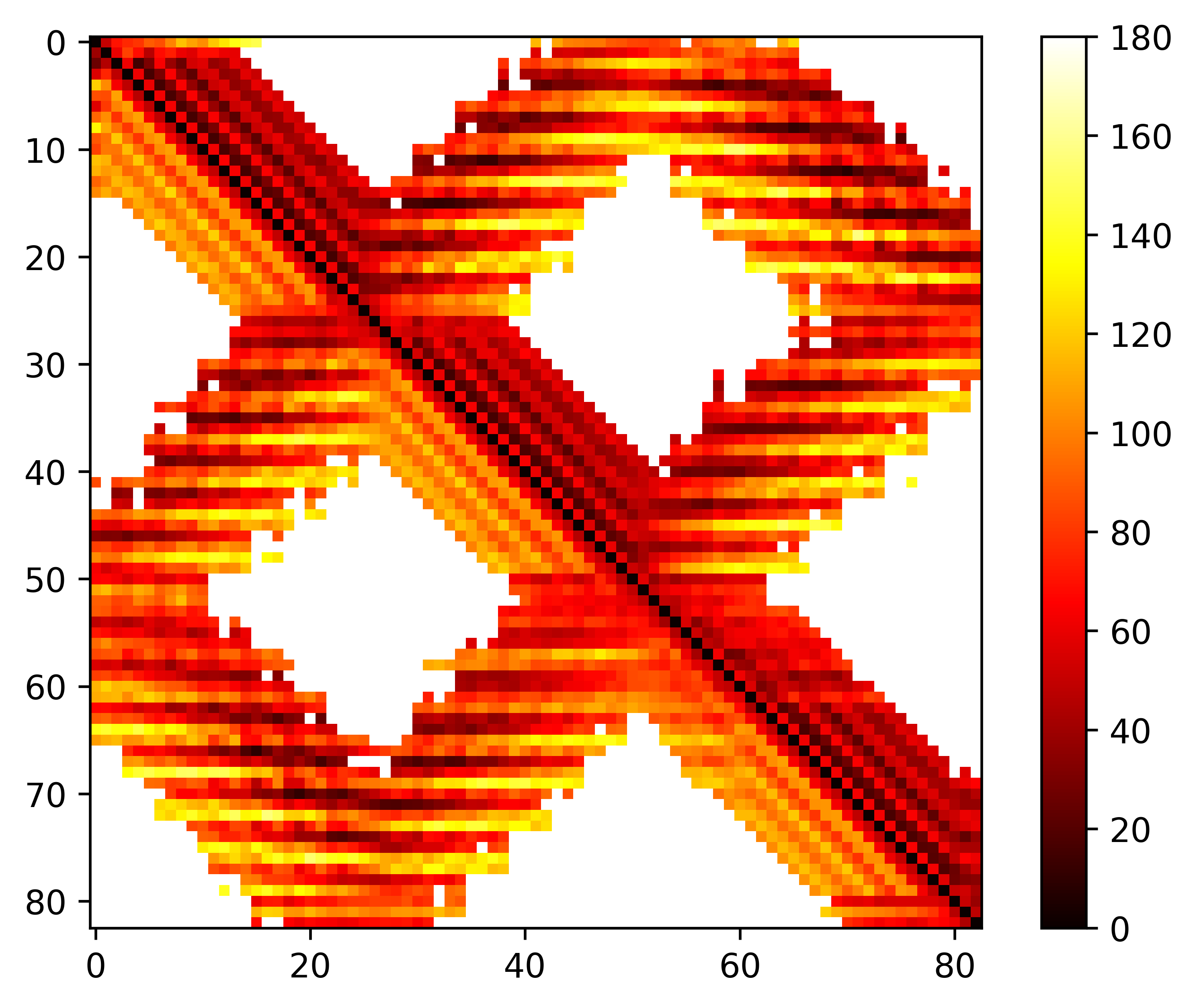

### seq.phi.png

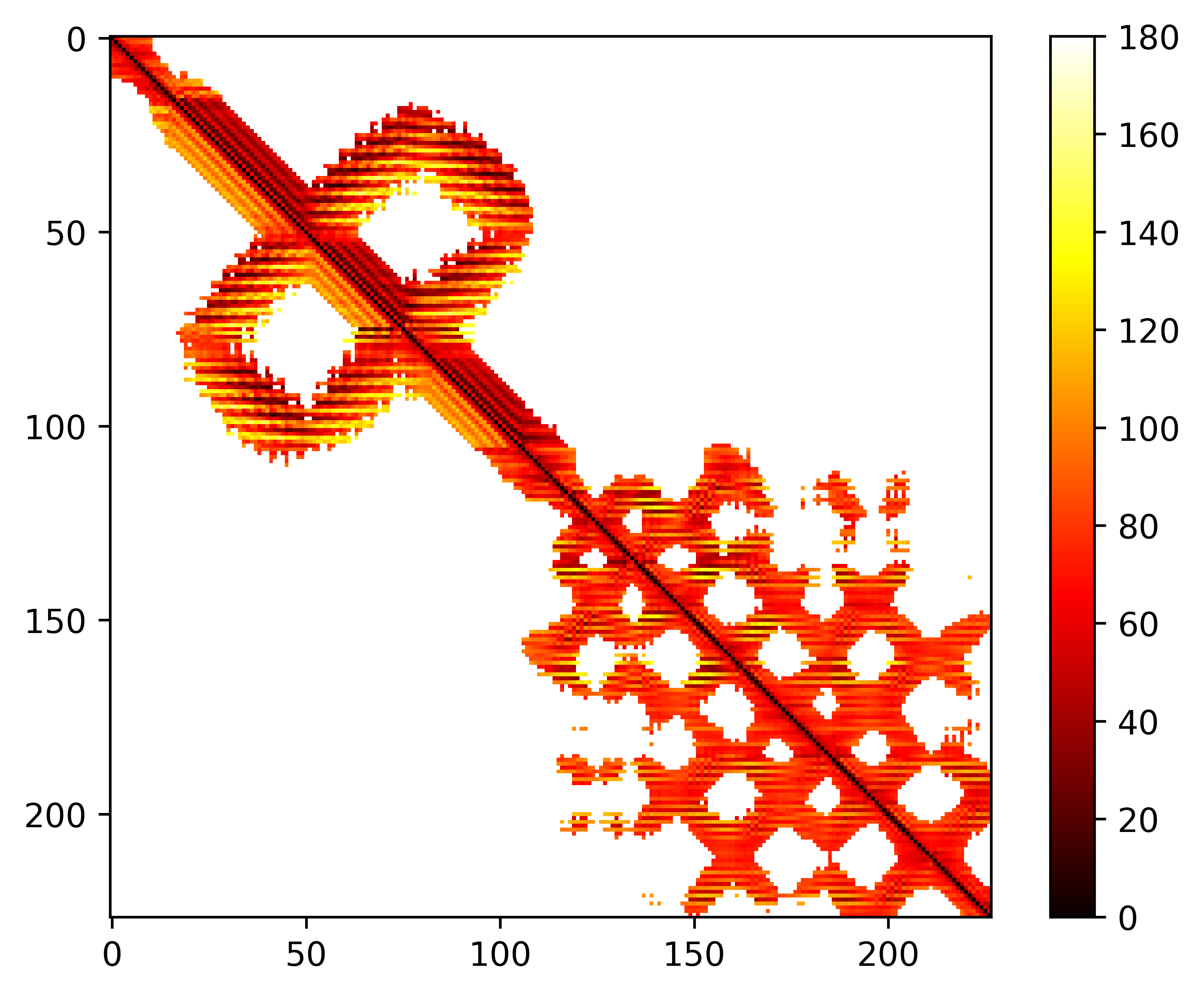

### seq.theta.png

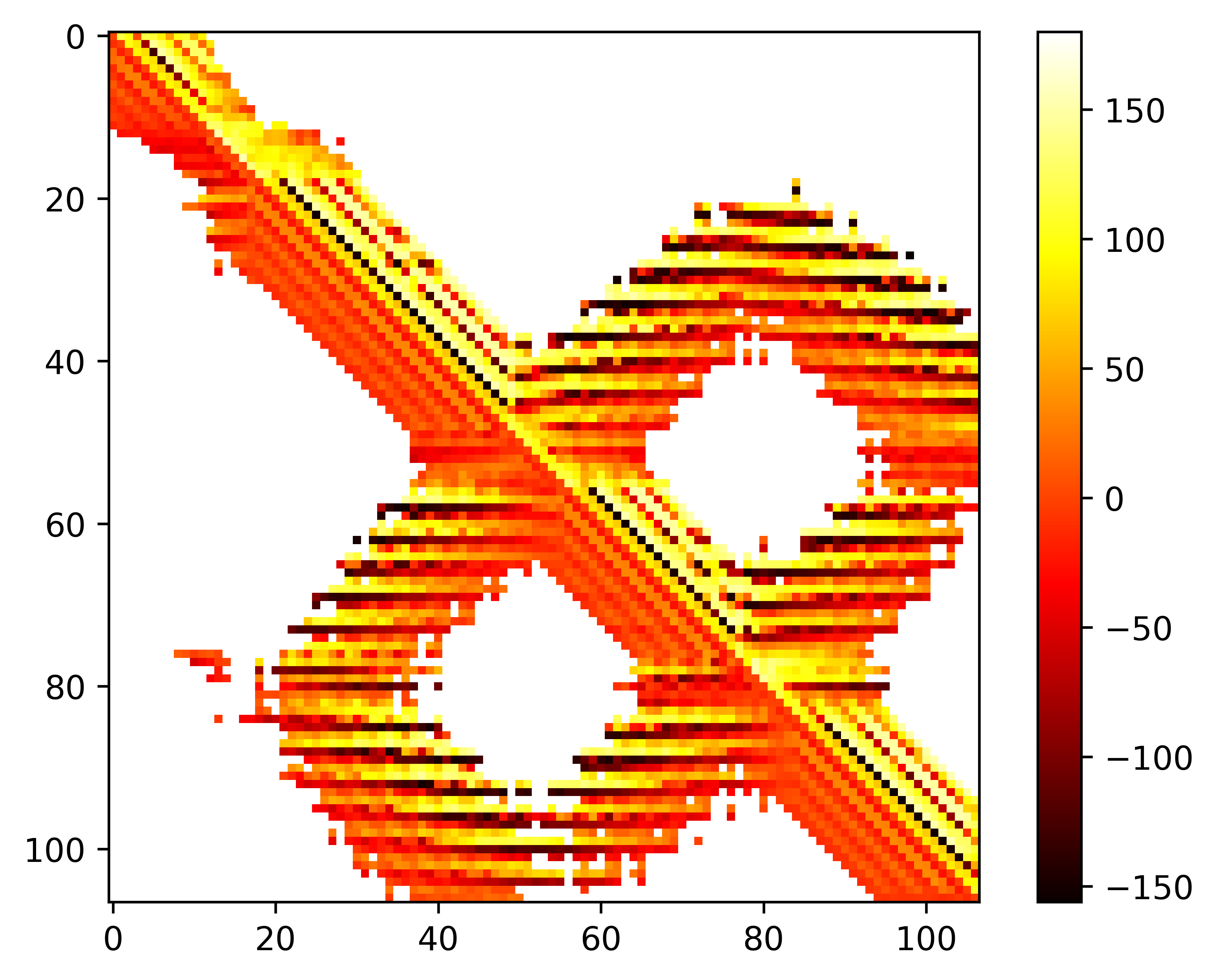

### seq.theta.png

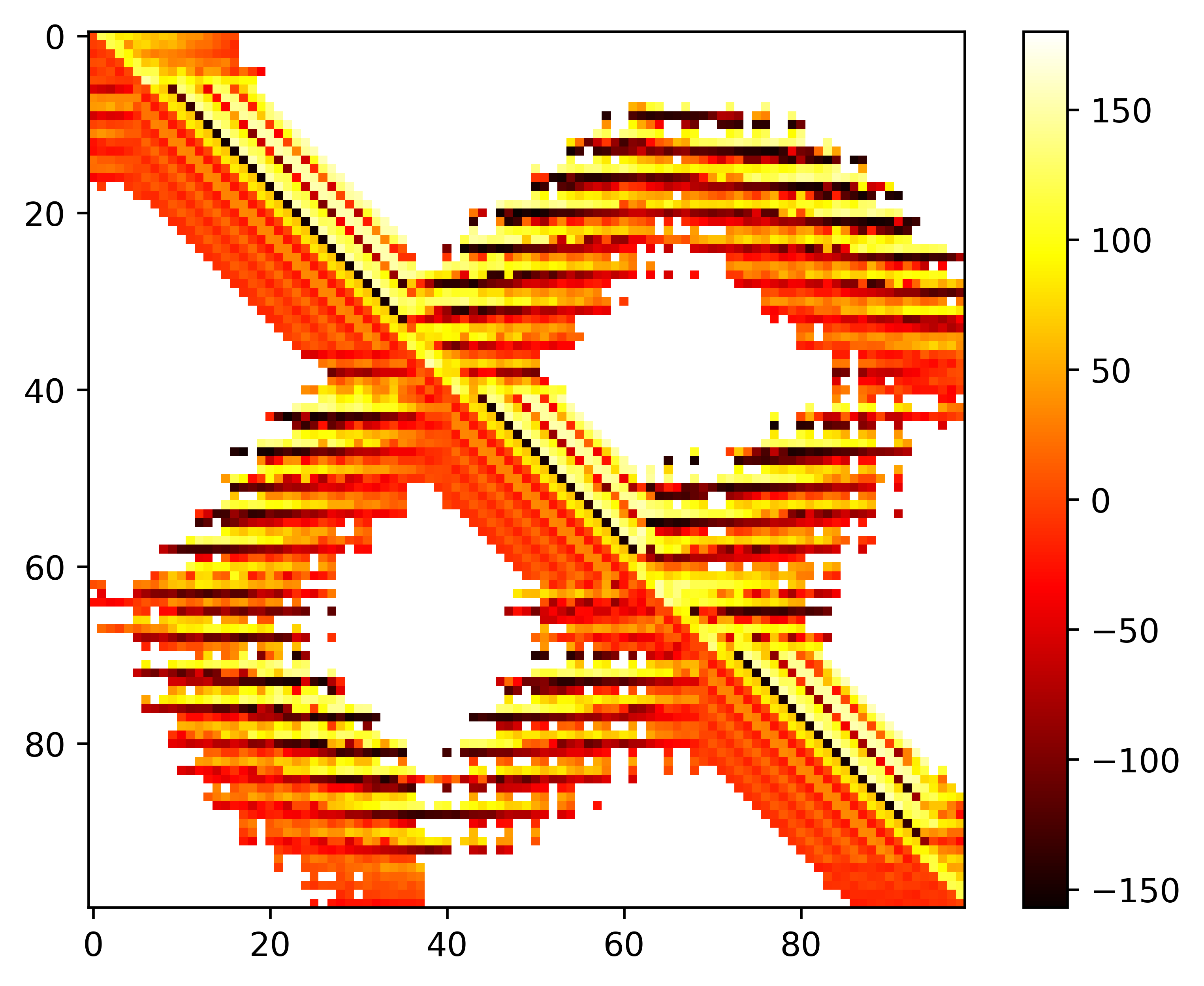

### seq.theta.png

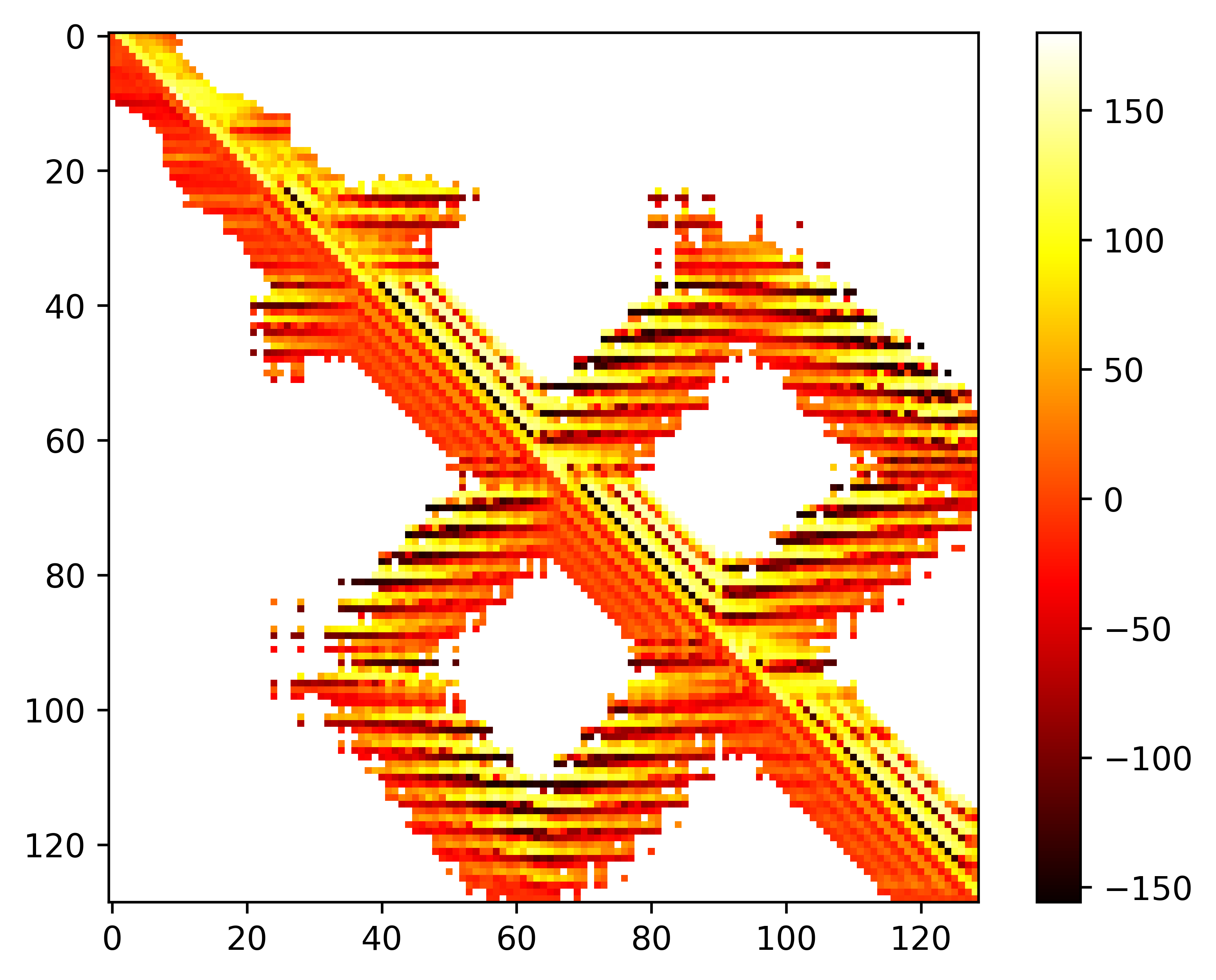

### seq.theta.png

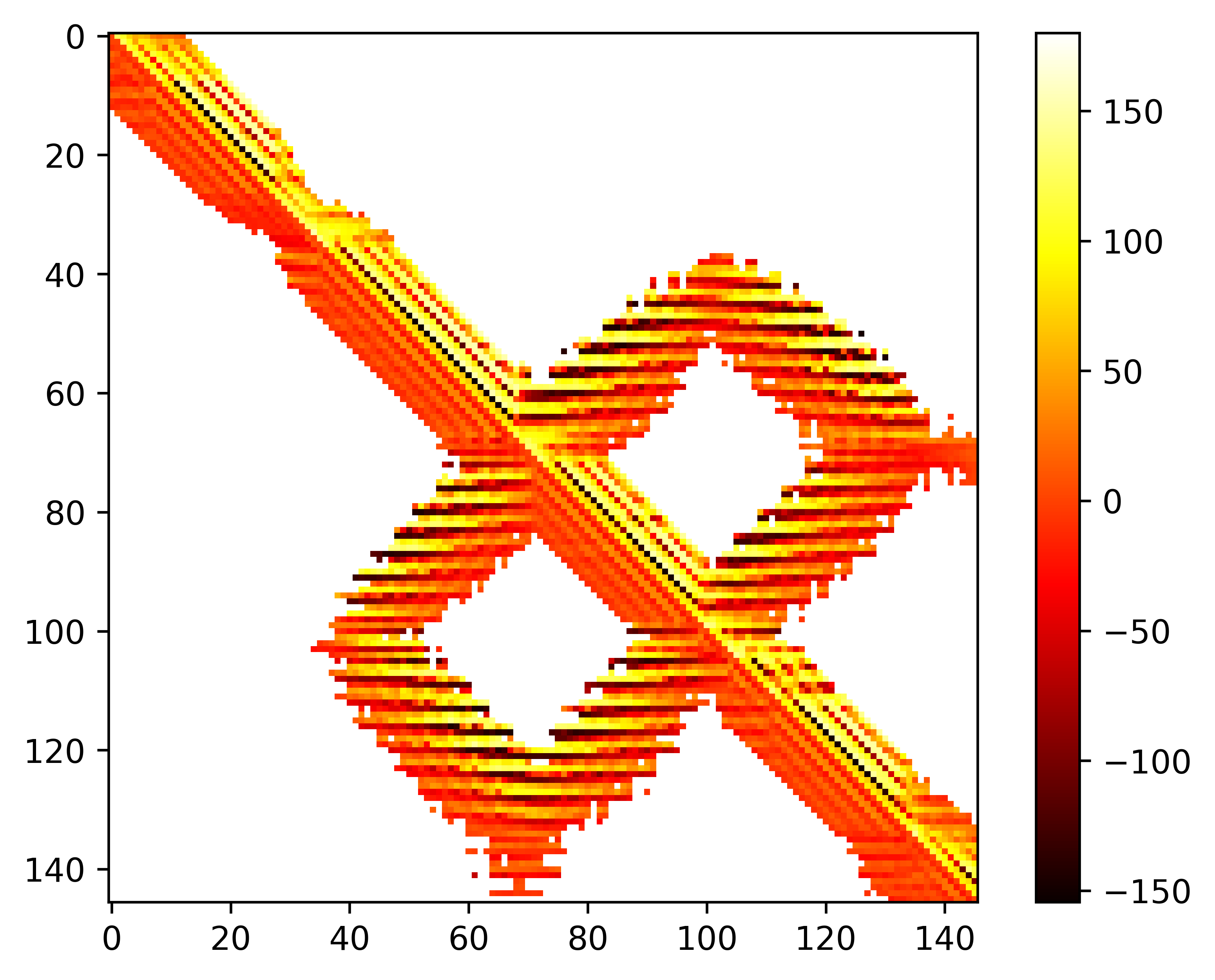

### seq.theta.png

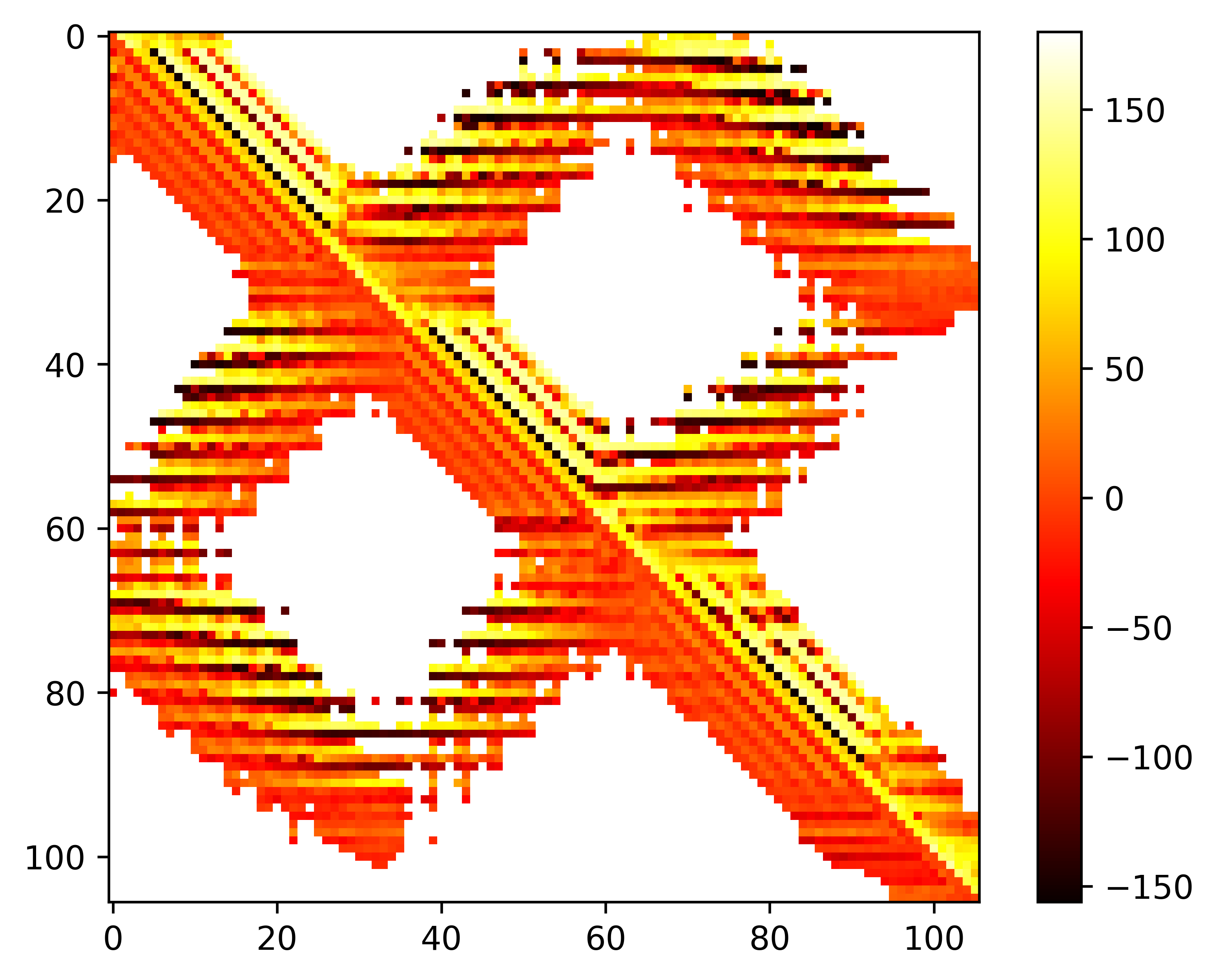

### seq.theta.png

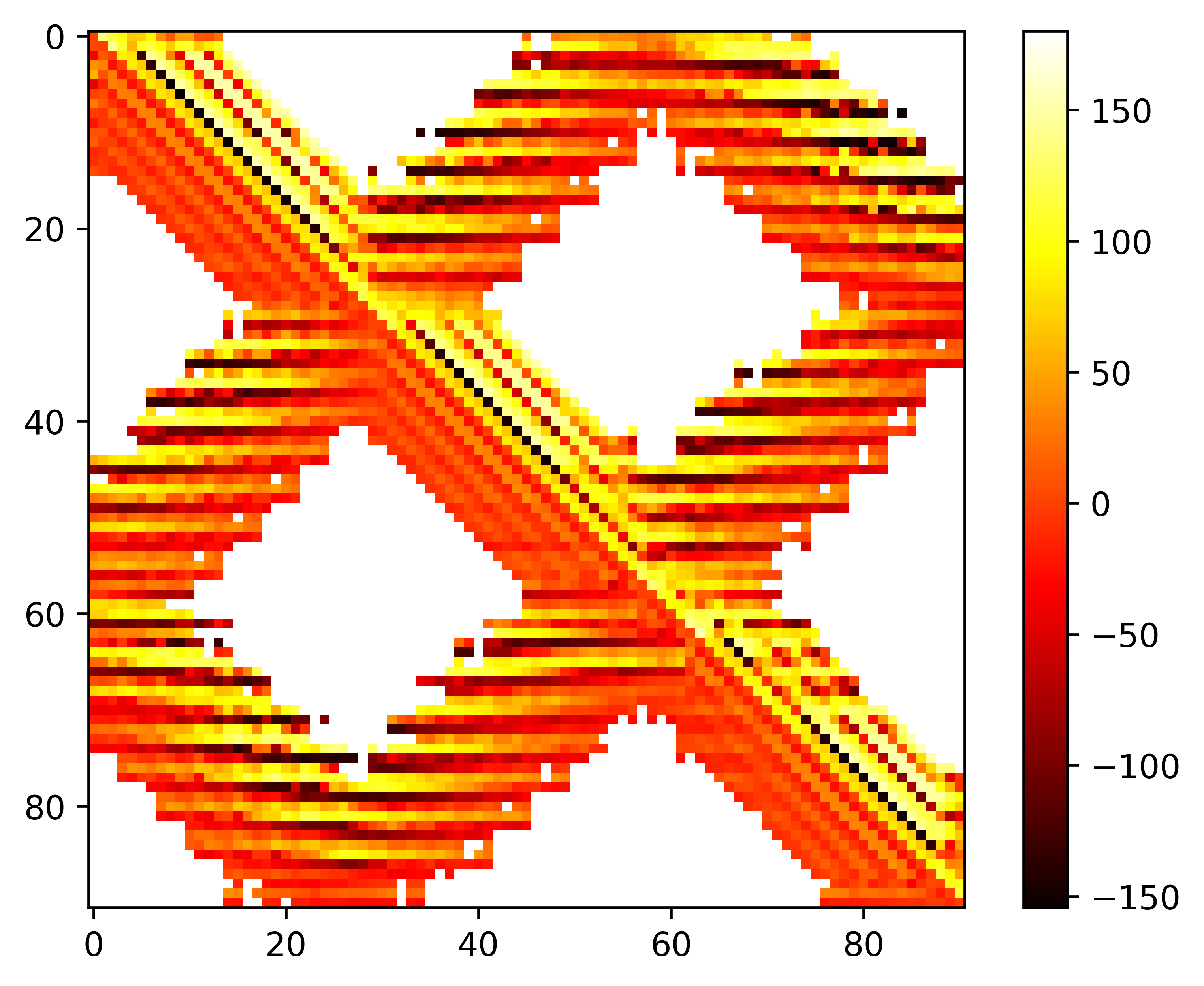

### seq.theta.png

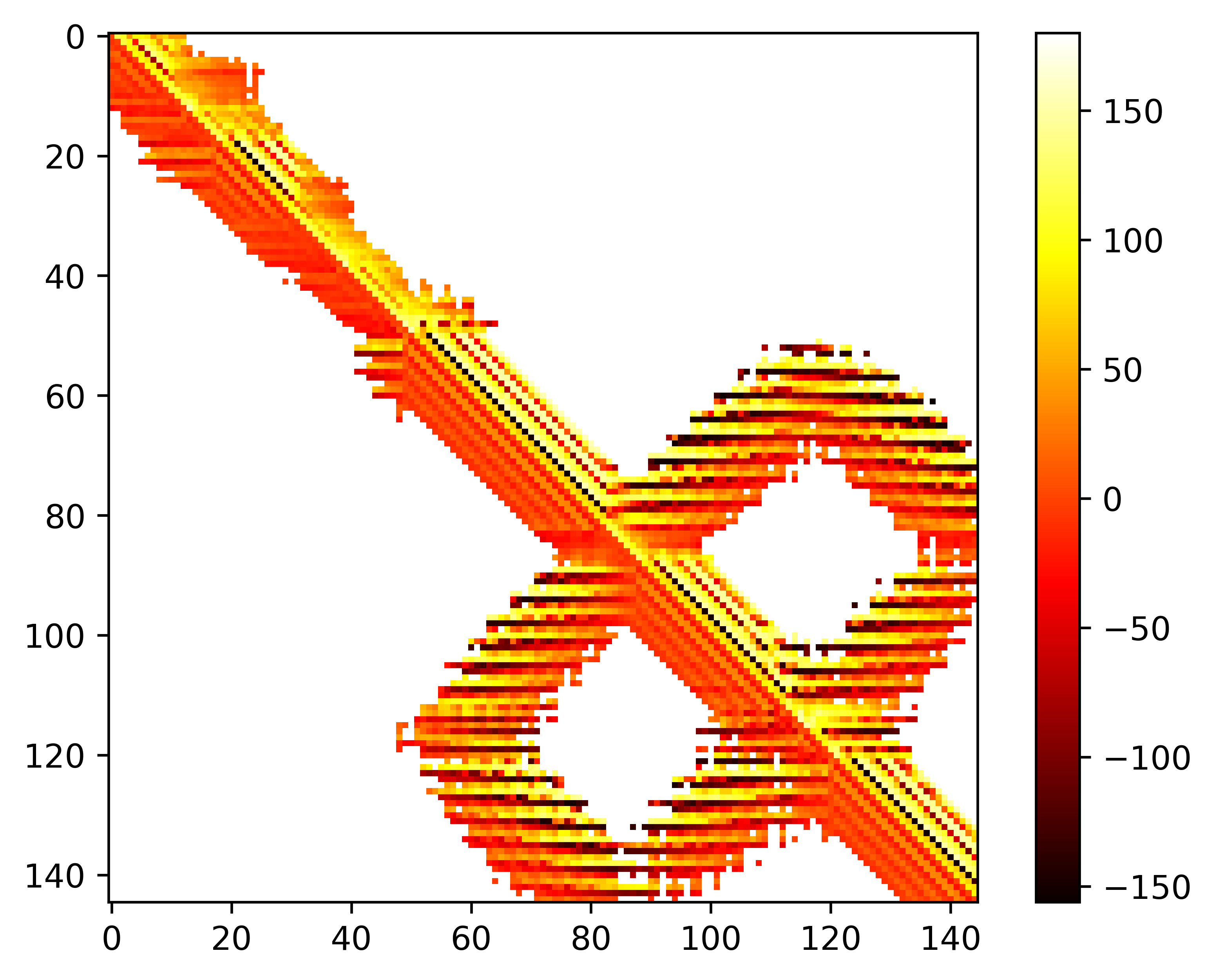

### seq.theta.png

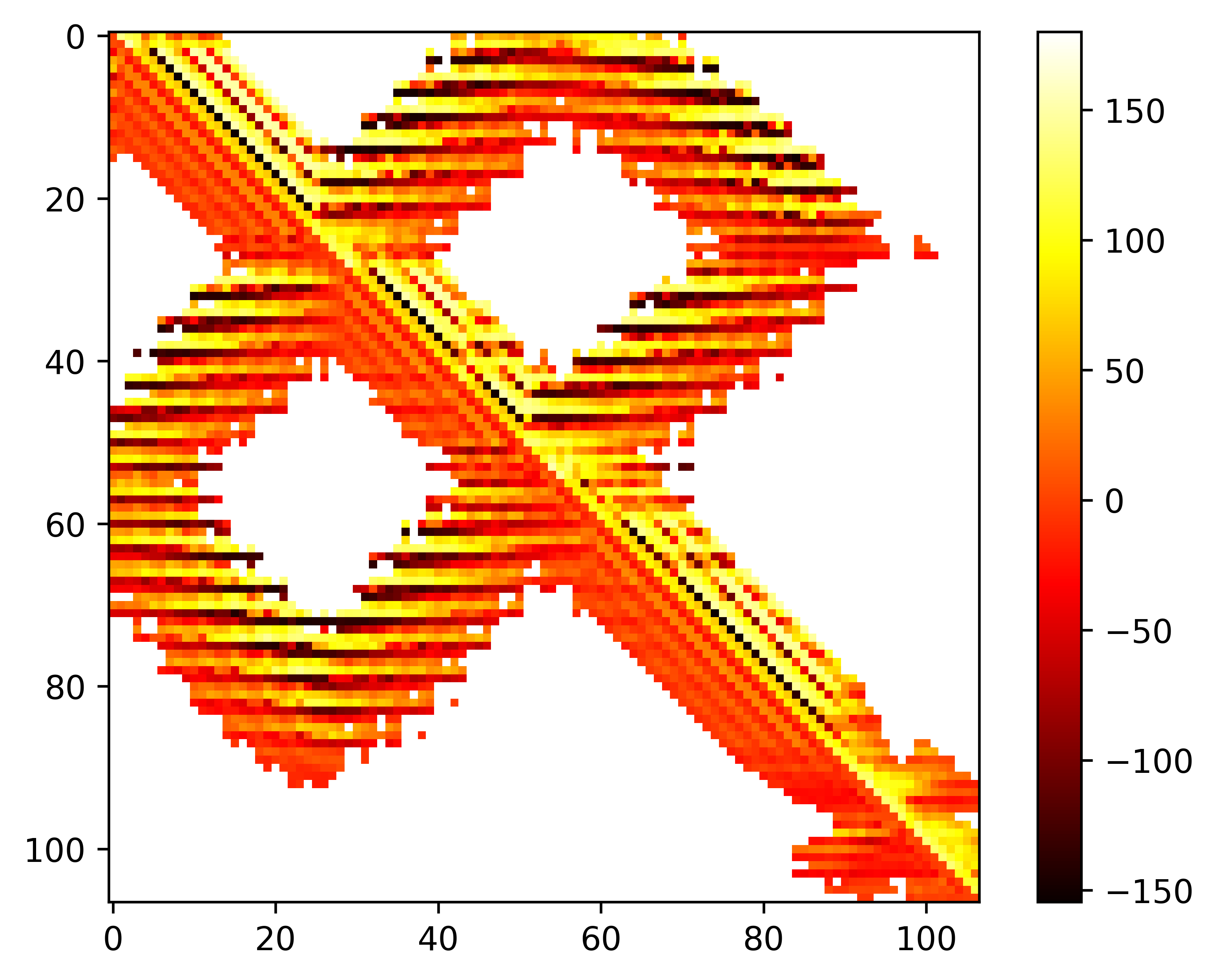

### seq.theta.png

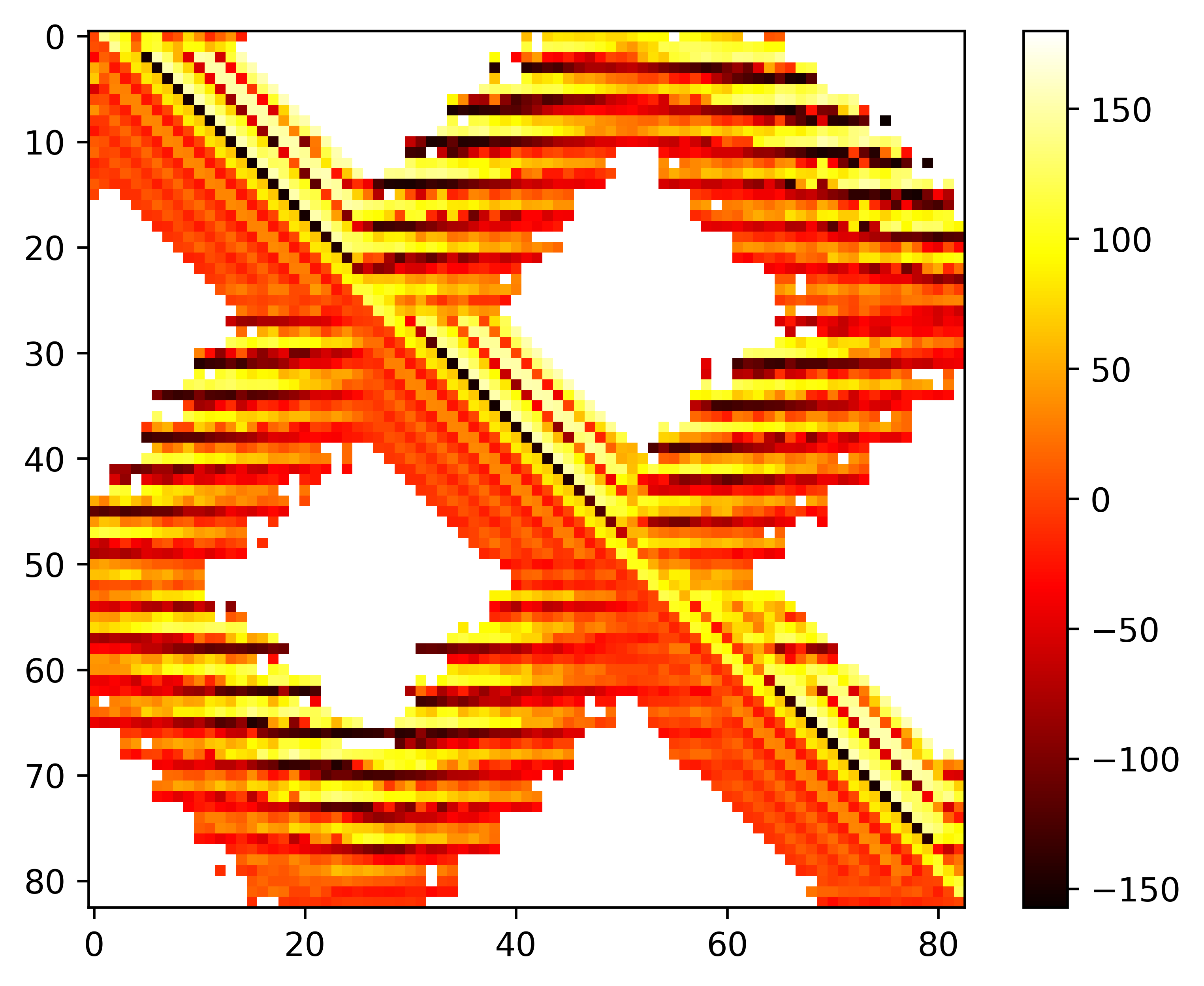

### seq.theta.png

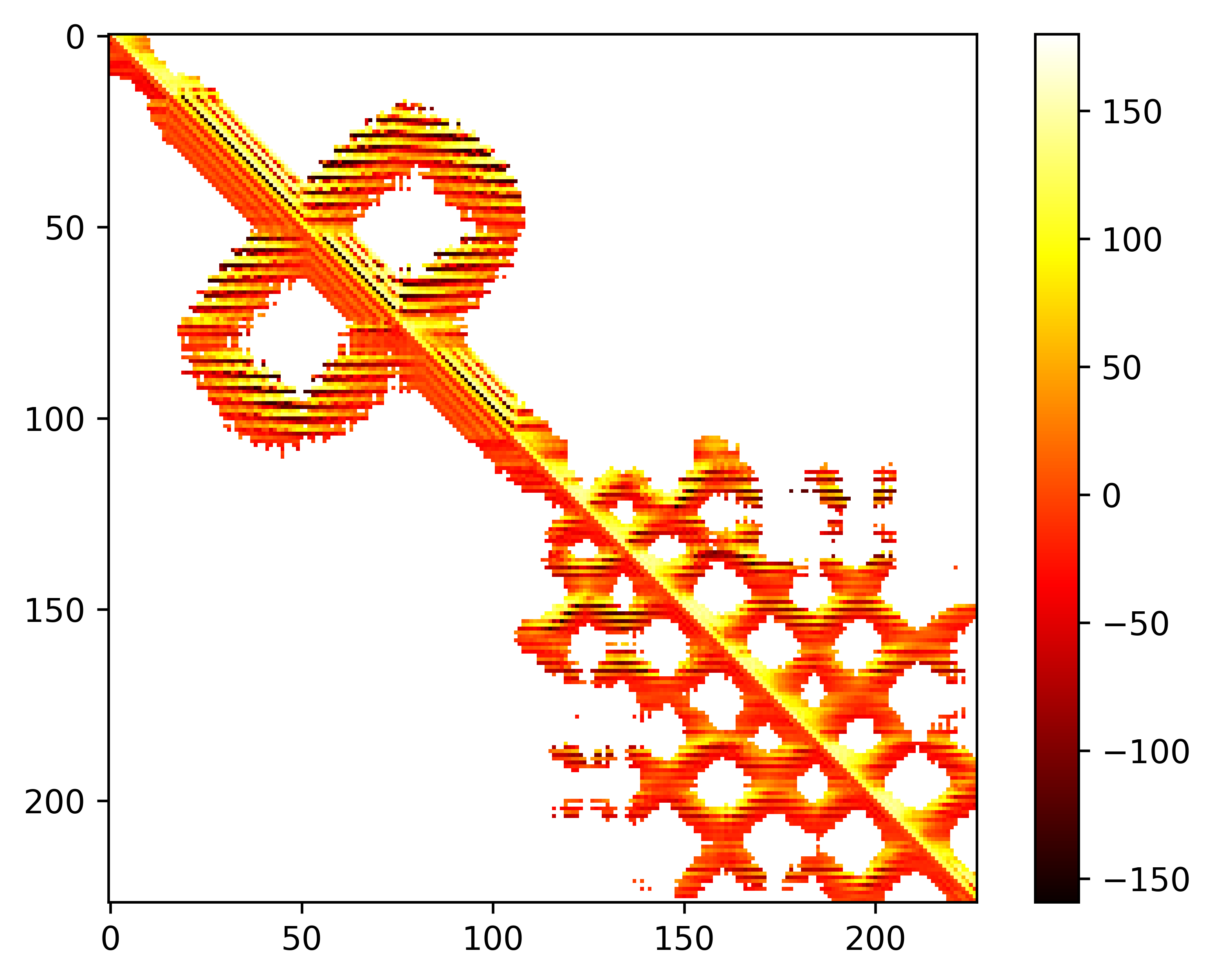
